# Evolutionary diversity of CXCL16*-*CXCR6: Convergent Substitutions and Recurrent Gene Loss in Sauropsids

**DOI:** 10.1101/2024.09.20.614078

**Authors:** Buddhabhushan Girish Salve, Sandhya Sharma, Nagarjun Vijay

## Abstract

The CXCL16-CXCR6 axis is crucial for regulating the persistence of CD8 tissue-resident memory T-cells (T_RM_). CXCR6 deficiency lowers T_RM_ cell numbers in the lungs and depletes ILC3s in the lamina propria, impairing mucosal defence. This axis is linked to diseases like HIV/SIV, cancer, and COVID-19. Together, these highlight that the CXCL16-CXCR6 axis is pivotal in host immunity. Previous studies of the CXCL16-CXCR6 axis found genetic variation among species but were limited to primates and rodents. To understand the evolution and diversity of CXCL16-CXCR6 across vertebrates, we compared ∼400 1-to-1 *CXCR6* orthologs spanning diverse vertebrates. The unique DRF motif of *CXCR6* facilitates leukocyte adhesion by interacting with cell surface-expressed *CXCL16* and plays a key role in G-protein selectivity during receptor signalling; however, our findings show that this motif is not universal. The DRF motif is restricted to mammals, turtles, and frogs, while the DRY motif, typical in other CKRs, is found in snakes and lizards. Most birds exhibit the DRL motif. These substitutions at the DRF motif affect the receptor─G_i/o_ protein interaction. We establish recurrent *CXCR6* gene loss in 10 out of 36 bird orders, including Galliformes and Passeriformes, Crocodilia, and Elapidae, attributed to segmental deletions and/or frame-disrupting changes. Notably, single-cell RNA sequencing of the lung shows a drop in T_RM_ cells in species with *CXCR6* loss, suggesting a possible link. The concurrent loss of *ITGAE, CXCL16* and *CXCR6* in chickens may have altered CD8 T_RM_ cell abundance, with implications for immunity against viral diseases and vaccines inducing CD8 T_RM_ cells.

## Introduction

Chemokines constitute a family of small proteins pivotal in orchestrating migration and maintaining homeostasis, thereby influencing the efficacy of immune defence (Zlotnik and Yoshie 2000; Hughes and Nibbs 2018). Specific cell surface receptors, i.e., chemokine receptors (CKRs), are required for chemoattractant activity (Allen et al. 2007). Together, chemokines and their receptors form the ligand-receptor axis.

*CXCR6* is a widely-conserved gene initially described as an HIV and SIV co-receptor (Alkhatib et al. 1997; Elliott et al. 2015; Wetzel et al. 2017). *CXCR6* belongs to class A G protein-coupled receptors (GPCRs), activated by its ligand through the N-terminal acidic residues and signals via G_i/o_ (Shimaoka et al. 2004a; Petit et al. 2008; Singh et al. 2015). The DRF motif in the cytoplasmic side of transmembrane 3 (TM3:H3C region) is unique to the CXCR6 protein, while other CKRs have a DRY motif (Koenen et al. 2017). The modification of DRY to DRF reduces calcium signalling and chemotactic response. However, the DRF to DRY affects adhesion capacity and indicates that the DRF motif in CXCR6 may be an adaptation for adhesion (Nomiyama and Yoshie 2015; Singh et al. 2015; Koenen et al. 2017). The DRF motif is also involved in the selective use of G_i/o_ in different cell types (Singh et al. 2010, 2015). *CXCL16*, the unique ligand of *CXCR6*, belongs to the CXC chemokine family and is involved in various immune and inflammatory processes (Matloubian et al. 2000). *CXCL16* plays a dual role in chemotaxis and adhesion with both soluble and transmembrane forms. The CXCL16- CXCR6 axis has also been implicated in various disease conditions such as cancer, HIV/SIV, and COVID-19 (Korbecki et al. 2021; Mabrouk et al. 2022).

Several factors influence how mallards and chickens respond to avian influenza virus (AIV), including variations in their mucosal defence mechanisms (Evseev and Magor 2019; Morris et al. 2023). Tissue-resident memory (T_RM_) cells are essential for mounting effective immunity against influenza infection (Hogan et al. 2001; Wu et al. 2014; Lee et al. 2024). In the context of compromised mucosal defence, the *CXCL16*-*CXCR6* signalling axis regulates the localisation of T_RM_ cells in different lung compartments and maintains airway T_RM_ cells. In *CXCR6*-/- mice, compromised cellular immunity to influenza infection has been observed (Wein et al. 2019). The *CXCL16*-*CXCR6* axis also regulates the diversity and function of innate lymphoid cells (ILC3). Specifically, the loss of *CXCR6* in mice resulted in the absence of NKp46+ ILC3s in the intestinal lamina propria (Satoh-Takayama et al. 2014). Notably, the lungs and intestines are primary sites for AIV infection in ducks (Evseev and Magor 2019). While *CXCR6* has been extensively studied in human and mouse model systems, its role in avian species remains largely unexplored. Hence, the function of this receptor may be different between birds and mammals, necessitating further functional characterisation of *CXCR6* across taxa.

T_RM_ cells are predominantly located in mucosal tissues, where pathogens and hosts commonly interact (Fan and Rudensky 2016). Previous studies have identified *CXCR6* as being expressed on T_RM_ cells and play a crucial role in their residency within tissues such as the lung, liver, skin, brain, and intestinal lamina propria (Sato et al. 2005; Tse et al. 2014; Chea et al. 2015; Zaid et al. 2017; Smolders et al. 2018; Wein et al. 2019; Rosen et al. 2022). Moreover, *CXCR6* serves as a core signature gene for CD8 T_RM_ cells, alongside other genes such as *CD69*, *ITGAE* (*CD103*), and *ITGA1* (*CD49a*) (Kumar et al. 2017; Zaid et al. 2017; Wein et al. 2019). T_RM_ cells play a significant role in pathogen immunity and tumour progression (Mueller and Mackay 2016; Kok et al. 2021; Yenyuwadee et al. 2022). This axis is involved in the activation and regulation (Germanov et al. 2008) and contributes to long-term MAIT cell retention (Yu et al. 2020).

Considering the crucial role of the *CXCL16*-*CXCR6* axis in the innate-adaptive immune system, comparative genomic analyses focusing on gene presence/absence patterns and adaptive evolution enable an understanding of the link between inter-species genetic differences and disease susceptibility. Previous comparative studies of *CXCL16*-*CXCR6* found genetic variation among species but were limited to primates and rodents only (Xu et al. 2018; He et al. 2020). Here, we leveraged the availability of large-scale genomic data and machine learning-based tools, such as Predicting Coupling probabilities of G-protein coupled receptors (PRECOGx) and Tool to infer Orthologs from Genome Alignments (TOGA), to conduct a comparative genomics analysis of ∼425 *CXCR6* gene orthologs to investigate the genetic diversification of *CXCR6* across vertebrates. Our findings unveil evidence of adaptive evolution, mainly through the DRF motif. Furthermore, our screening identified the recurring loss of the *CXCR6* and *CXCL16* within the Sauropsids lineage (including Aves) while being highly conserved across mammalian orders.

## Materials and methods

### Evaluation of conserved synteny to identify one-to-one orthologues

To explore the evolutionary trajectory of *CXCR6* and its ligand *CXCL16*, we conducted a comparative genomic analysis of ∼425 vertebrate species spanning fishes, amphibians, reptiles, birds, and mammals **(Supplementary Table 1)**. Our analysis encompasses the full spectrum of vertebrate diversity, encompassing over 50 orders. We used BLASTn (Camacho et al. 2009) to search for 1-to-1 orthologs of *CXCR6* in 22 representative species from key vertebrate clades using the coding sequence of the human gene (ENST00000458629) **(Supplementary Table 2)**. A similar search strategy was used to identify potential orthologs of *CXCR6* genes in Aves, Crocodilia, and Squamata using the closest relative species to the group, such as ostrich (*Struthio camelus*; ENSSCUT00000025423) and central bearded dragon (*Pogona vitticeps*; ENSPVIT00000003078).

Further, we relied on the conserved micro-synteny to distinguish BLASTn hits from other CKRs. The NCBI Genome Data Viewer (Rangwala et al. 2021) facilitated the visualisation of BLASTn hits, identifying genes flanking *CXCR6* and enabling comparisons of the genomic landscape across species and was further confirmed using ENSEMBL annotations (Martin et al. 2023). For species lacking genome annotations in the NCBI database, we used the TOGA (tools to infer orthologues from genome alignment) pipeline (Kirilenko et al. 2023) to annotate the *CXCR6* gene **(Supplementary Table 1)**. Additionally, the species with low-quality protein annotations in the NCBI dataset underwent reassessment using TOGA and verified by short-read Illumina data.

### Validation of Genome assembly and Finding of gene loss

To ensure the absence of the gene is not due to assembly artifacts, we employed long overlapping reads spanning the flanking region of *CXCR6* to validate the genome assembly. Publicly available long-read data (PacBio and Nanopore) validated the assemblies of chicken, mallard, zebra finch, pigeon, downy woodpecker, budgerigar, Northern carmine bee-eater, Anna’s hummingbird, kakapo, common swift, crocodiles, alligators, and the leatherback sea turtle **(Supplementary Table 3)**. Read mapping was done using BWA-MEM v0.7.17 (Jung and Han 2022) to their respective reference genomes. Repeats were annotated using RepeatMasker v4.1.5, and visualization was done using the UCSC genome browser (Kent et al. 2002).

Subsequently, in the species where the assembly was correct at the *CXCR6* gene locus, three scenarios are possible: (1) segmental deletion resulting in gene loss, (2) translocation to other genomic locations, and (3) frame-disrupting changes resulting in gene loss. In the case of segmental deletion, the closest species with intact *CXCR6* was used to estimate the deletion size. The segmental deletion was further confirmed using pairwise dot and circos plots (Krzywinski et al. 2009). The possibility of translocation to other locations was ruled out using BLASTn searches of available genome assemblies and raw read databases of the focal species. Finally, the frame-disruptive changes were identified using the TOGA (Kirilenko et al. 2023) pipeline and verified using the BLASTn search of the SRA database **(Supplementary Table 4)**.

### Patterns of Molecular Evolution: Conservation, Selection, and Convergence

Multiple sequence alignments (MSAs) of intact *CXCR6* from Aves and other key representatives from vertebrate clades were generated using ClusatalX v2.1 (Larkin et al. 2007). The MSAs were annotated for the transmembrane region, and conserved motifs were marked in pyMSAviz (https://moshi4.github.io/pyMSAviz/). Annotations of the conserved motifs and amino acid residues were done based on the study by Nomiyama and Yoshie et al. (Nomiyama and Yoshie 2015). EMBOSS v6.6 pepinfo, and pepstats were used to plot and calculate amino acid properties in protein sequences, respectively.

The MSA of *CXCR6* orthologs and the time-calibrated phylogenetic trees were used for molecular evolutionary analysis. The phylogenetic tree was obtained from the Timetree website (http://www.timetree.org/). The magnitude of GC-biased gene conversion (gBGC) was assessed using MapNH v1.3.0 and phastBias (**Supplementary Table 5**). To detect the selection pressure on the *CXCR6* gene across orders of vertebrate species, we used several dN/dS metric-based software tools. The alignment and tree are analyzed by Hyphy v2.3.14 packages site and branch-site models, i.e., FEL, MEME, aBSREL, BUSTED, and RELAX (Kosakovsky Pond et al. 2005) **(Supplementary Table 6)** and PAML (Yang 2007) using the codeML application **(Supplementary Table 7).** Multiple-testing correction was done using the FDR method.

To find out the sites that changed along with CXCR6’s DRF/DRY/DRL motif, we used CSUBST (Fukushima and Pollock 2023) to detect convergent substitution. The convergent substitutions were identified by specifying foreground branches, mapped to protein structure, and visualized using PyMOL (https://pymol.org/2/) (Schrödinger, LLC 2015).

### *CXCR6* G-protein coupling probability and MD simulation

To evaluate the effect of amino acid substitutions at the DRF motif of CXCR6 on G-protein coupling, we used the machine-learning predictor PRECOGx (Matic et al. 2022) and GPCRsignal (Miszta et al. 2021). Briefly, CXCR6 1-to-1 orthologs were used to find G-protein coupling probabilities for different types of G-proteins **(Supplementary Table 8)**. To identify which amino acid residues interact with G-proteins, we used the cryo-EM structure (PDB id: 6WWZ) of human CCR6 in a complex with CCL20 and G_o_ protein (Wasilko et al. 2020). Mutations were provided **(Supplementary Table 9)**, and MD simulations were run for 25 ns with 200 frames. Membrane thickness was kept at 33.8 Å.

### Analysis of the expression profile of the *CXCR6* gene

*CXCR6* is expressed in the spleen, lung, liver tissues, CD4 and CD8 cells (Latta et al. 2007; Chea et al. 2015; Hughes and Nibbs 2018; Wein et al. 2019; Mabrouk et al. 2022). Therefore, we screened the expression status of the *CXCR6* gene in chicken CD4 and CD8 cells (PRJNA713900), liver, and T cell (PRJEB27455/FAANG project) transcriptome (Dai et al. 2021; Kern et al. 2021). For the species where *CXCR6* gene remnants are present, the RNA-seq reads from public datasets **(Supplementary Table 10)** were aligned to the reference genomes using STAR v2.7.1a read mapper (Dobin 2019). The flanking exons of the *FYCO1* gene are positive controls within the species. The integrative genomics viewer (IGV) browser (Thorvaldsdottir et al. 2013) visualized the transcriptional status.

### Lung Single-cell RNA-seq analysis

To compare the effect of *CXCR6* gene loss on CD8 T_RM_ cells in the lung, we used mallard (*Anas platyrhynchos*) and pigeon (*Columba livia*) single-cell RNA-seq data available on the SPEED pan-species single-cell database (Chen et al. 2023). We used *CD69*, *CD103* (*ITGAE*), and *CD49a* (*ITGA1*) as marker genes for CD8 T_RM_ cells (Kumar et al. 2017; Zaid et al. 2017; Wein et al. 2019). Previously, it was discovered that these three genes controlled the expression of the *CXCR6* gene, which is critical for the retention of T_RM_ cells in the lung (Morgan et al. 2008; Wein et al. 2019; Mabrouk et al. 2022). We also retrieved lung single cell transcriptome sequencing data from the NCBI database (Accession ID: PRJNA747757; for mallard and pigeon)(Chen et al. 2021) and PRJNA834764 (for chicken) (Liu et al. 2023). All single-cell transcriptomic data were processed using Cell Ranger v7.2.0 (10× Genomics), and Single-cell analysis was conducted using Seurat v5. (Hao et al. 2024) (see **Supplementary Text S1** for Tipps and Tricks in methods).

## Results

### Gene collinearity ensures unambiguous ortholog identification

Comparing the gene order in the flanking regions of *CXCR6* across 22 representative species from major vertebrate clades, we found that *CXCR6* is situated between exon 14 and 15 of the *FYCO1* gene. In the analyzed vertebrate genomes (**Supplementary Table 1**), including humans, the left flank of the *FYCO1* gene contains the *CCR9* gene, and the right flank contains the *XCR1* gene **(Fig. 1A; Supplementary Table 2)**. In the case of the western clawed frog (*Xenopus tropicalis*), although the gene (*XCR1*) on the right flank is conserved, the gene order is changed as the *FYCO1* gene is missing, and the left flank contains the *GLB1* gene. However, in other species of Amphibia class, such as Gaboon caecilian (*Geotrypetes seraphini*), the *CXCR6* gene is present within the intron of the *FYCO1* gene. The conserved synteny observed at the *CXCR6* locus ensures the identification of 1-to-1 orthologs. Most mammal genomes screened for the *CXCL16* gene contain *ZMYND15* and *MED11* on the flanks, with altered synteny in other vertebrates, such as frogs (**Fig. 1A**).

**Fig. 1:**
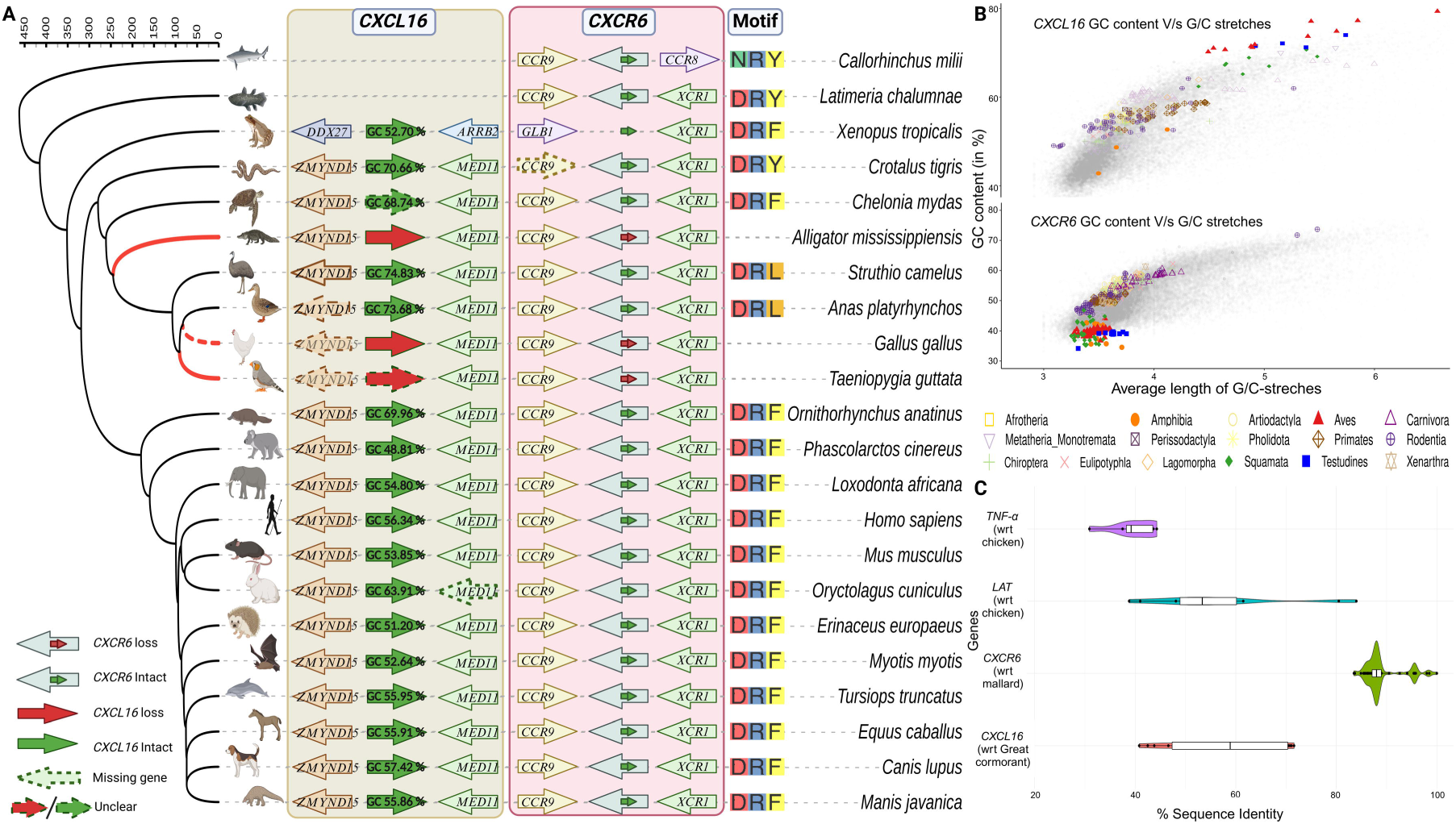
Conservation of *CXCR6* and *CXCL16* gene synteny across the vertebrates. **A.** Phylogenetic relationships among vertebrate species are shown with solid red lines for branches with the loss of the *CXCR6* due to gene-inactivating mutations and dashed red lines representing gene loss due to segmental deletions. The solid green arrows indicate the intactness of *CXCL16*, while the solid red arrows represent loss. The GC content of the *CXCL16* gene is mentioned above the arrow. The small red and green arrows within the big arrow represent *CXCR6* in the intron of the *FYCO1* gene (big arrow). The red and green colours indicate its absence and presence, respectively. Dotted arrow borders indicate the genes missing in the species with their names within the arrows. The red/green coloured arrows with dotted borders are the species in which the status of the gene is unclear. The direction of the arrows indicates gene orientation. The conserved DRF motif (species-specific changes like NRY, DRY, and DRL) is shown for that species. The time-calibrated phylogenetic tree was downloaded from the Timetree website. The conserved motifs are inferred based on the multiple sequence alignment of these representative species. The iTOLv6 was used to annotate branches of the phylogenetic tree. **B.** *CXCL16* gene has a high GC content with long stretches of G/C in Sauropsids. GC content is plotted against the mean length of G/C stretches. The background grey dots show the distribution of GC content and average G/C-stretch length in the chicken RefSeq genes >300 base pairs. **C.** Sequence divergence comparison between *CXCL16*, *CXCR6*, *LAT*, and *TNF-α*. The *CXCL16* shows high sequence divergence like *LAT* and *TNF-α*.

Distinguishing gene loss from "missing genes" remains a challenge, especially in bird genomes, due to the challenges of sequencing and assembly of GC-rich regions and those with repetitive content (for example, telomeres) (Hron et al. 2015; Rohde et al. 2018; Janusova et al. 2023). We found evidence of high GC content and prevalence of G/C stretches in the case of *CXCL16*, a gene whose syntenic locus could not be recovered in bird genomes (**Fig. 1B**). However, an intact *CXCL16* could be recovered in the harpy eagle (*Harpia harpyja*; GC content: 70.68%) and great cormorant (*Phalacrocorax carbo*; GC content: 71.02%). The *CXCL16* orthologs could be recovered in other birds, such as Eurasian goshawk (*Accipiter gentilis*), common buzzard (*Buteo buteo*), Cory’s shearwater (*Calonectris borealis*), White-tailed eagle (*Haliaeetus albicilla*), Darwin’s rhea (*Rhea pennata*), ostrich (*Struthio camelus)* (**Supplementary Figure S1**), mallard (*Anas platyrhynchos*) and swan goose (*Anser cygnoides*). However, in the case of chicken (*Gallus gallus*), the availability of high-quality genomes and multi-platform long-read datasets allowed us to assert the loss of *CXCL16* (**Supplementary Figure S2**). We identified the loss of *CXCL16* in at least one species in all three major lineages (i.e., birds, Crocodilia, and elapids) with *CXCR6* loss (**Supplementary Figure S3-S5**).

Our inability to identify remnants of *CXCL16* in several species suggests the possibility of a highly diverged copy may fall into the group of "missing genes" like *TNF-α* and *LAT* that have now been recovered as highly diverged orthologs. Upon comparing the percentage of nucleotide sequence identity among orthologs of *TNF-α*, *LAT*, *CXCR6,* and *CXCL16* (**Fig. 1C**), the *CXCL16* orthologs are highly diverged. Interestingly, *CXCL16* gene sequences were recovered in some mallard short-read data and were found to be identical to the human *CXCL16* sequence. However, upon verification, these short-read data were identified as contaminated with human DNA and did not match the actual sequence of *CXCL16* recovered from mallard long-read data (**Supplementary Figure 6**). Overall, *CXCL16* has substantial sequence divergence, GC-rich content, and prevalence of long G/C stretches, a pattern not seen in the case of *CXCR6* (see **Supplementary Text S2**).

### Convergence in the DRF motif may govern G-protein coupling

CXCR6 is a ∼342-amino-acid residue protein with seven transmembrane helices with conserved motifs such as TxP, DRF, and NPxxY(x)_5,6_F essential for GPCR receptor activation (**Fig. 2A**). However, N-terminal Cys residues and the CWxP motif found in other CKRs are not found in *CXCR6* (with only proline of this motif conserved) which has low sequence conservation in the terminal regions (**Fig. 2B**). Several amino-acid substitutions at conserved positions are specific to birds compared to mammals (**Fig. 2A**). Additionally, our comparison of representative species of mammal (human) and bird (mallard) found the terminal regions of CXCR6 have drastically different hydrophobicity (**Fig. 2C, Supplementary Figure 7-8**). A quantitative comparison of the acidic residues in the N-terminal regions among vertebrate clades found that Aves and Testudines have fewer acidic residues compared to Amphibia, Lepidosauria, and Mammalia (median difference Aves: p-Value < 0.0001 and Testudine: p-value <0.05, using Kruskal–Wallis test) **(Fig. 2D)** (see **Supplementary Figure 9**, for all pairwise comparisons). The H3C region also has bird-specific substitutions (**Fig. 2E**).

**Fig. 2:**
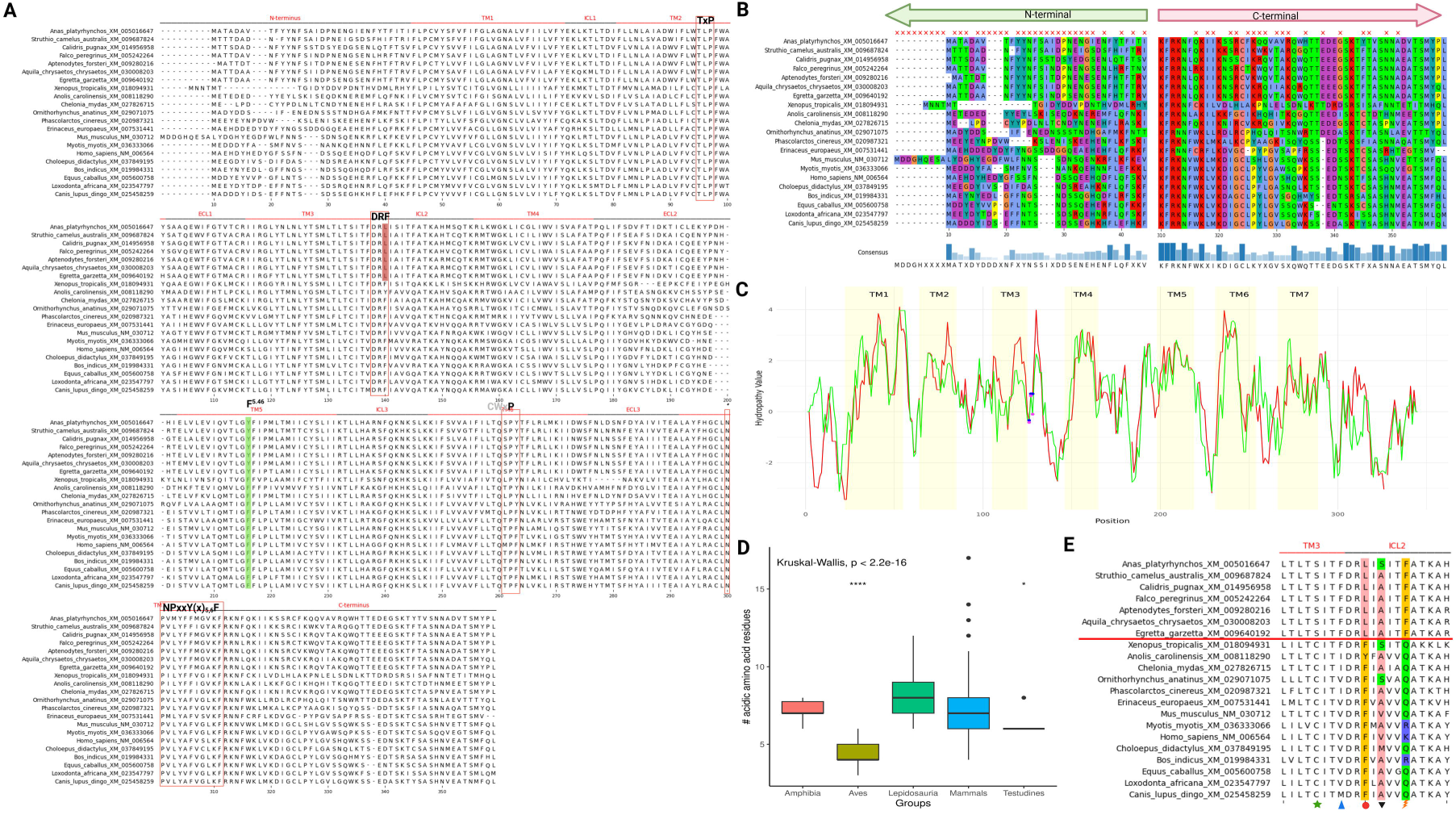
Evolutionary diversification of CXCR6 protein-coding sequences. **A.** CXCR6 protein sequences from 21 vertebrate species, consisting of birds and other key representative species. The conserved motifs (The TxP, DRF, and NPxxY(x)5,6F) are shown with rectangular red boxes. The only conserved residue proline from the CWxP motif is shown with the rectangular red box in the TM-6 region. Changes in DRF, i.e., F to L, are highlighted in red. The conserved residues F^5.46^ in TM-5 are shown with a light-green highlight. TM regions (I-VII) are labelled as annotated in the UniProt database for the human CXCR6 (O00574). pyMSAviz is used for visualisation (https://github.com/moshi4/pyMSAviz). **B.** Terminal sequence conservation. The N-terminal and C-terminal regions show less amino acid conservation. The red crosses on the alignment indicate conservation of less than 50%. The bottom blue bars represent the consensus of the sequence. The amino acids coloured are based on the Clustal format. **C.** A comparative Kyte-Doolittle hydropathy character of the human (red line) and mallard (green line). Human and mallard hydropathy was obtained by giving amino acid sequences in EMBOSS Pepinfo. The hydropathy value (y-axis) and residue position (x-axis) are plotted in R using ggplot2. Transmembrane regions are shown in yellow highlights. DRF and DRL residues in humans and mallards are shown with blue and pink dots, respectively. **D.** Comparison of N-terminal acidic amino acid residues across vertebrate groups. The x-axis represents key vertebrate groups such as Amphibia, Aves, Mammals, Lepidosauria, and Testudines. The y-axis represents the number of acidic residues. A comparison of medians is made with the Kruskal-Wallis test using ggplot2 in R. **E.** Bird-specific changes in the H3C region of CXCR6. The horizontal red line separating birds from other vertebrate species shows the changes in the amino acid of the cytoplasmic site of TM-III. The change of C to S (green star), V to F (blue triangle), F to L (red circle), V to A (inverted black triangle), and K to F (orange thunderbolt).

The DRF motif of the H3C region found exclusively in CXCR6 is widely conserved across mammals, frogs, and turtles. Notably, several species of snakes and lizards have the DRY, while birds have the DRL **(Fig. 1A** and **Fig. 2E, Supplementary Figure 10)**. Vertebrate species can be categorized based on their DRY/DRF/DRL status **(Fig. 3A).** With this classification, we identified three groups with the mammal-like DRF motif (placentals, frogs, and turtles), the squamate-like DRY motif (squamates and *Geotrypetes seraphini*), and aves-like DRL motif (aves and *Sphenodon punctatus*). Additionally, the mammal-like DRF motif is found among monotremes and most marsupials. However, the Didelphimorphia species (Agile gracile opossum (*Gracilinanus agilis*) and Gray short-tailed opossum (*Monodelphis domestica*)) have the aves-like DRL motif (**Supplementary Figure S11**). A systematic search for convergent substitutions correctly identified the F128L (DRF to DRL substitution) in aves-like and F128Y (DRF to DRY substitution) in squamate-like lineages **(Fig. 3B)**. Aves-like lineages have another convergent substitution, F200Y **(Fig. 3C)** (position F^5.47^, according to Ballesteros–Weinstein numbering scheme). In the case of squamates-like lineages, we found a convergent substitution at F299L (F^8.54^). While other birds, including Accipitriformes species, have a DRL motif, the harpy eagle (*Harpia harpyja*) has DRF with evidence of site-specific positive selection at the third residue (MEME: LRT = 3.308, p = 0.0908. FEL: LRT = 3.306, p = 0.0690) **(Supplementary Figure 12-13)**. The most common substitution detected by the MEME model is ctT(L)>ctC(L) compared to the Ctt(L)>Ttt(F) substitution in harpy eagle.

**Fig. 3:**
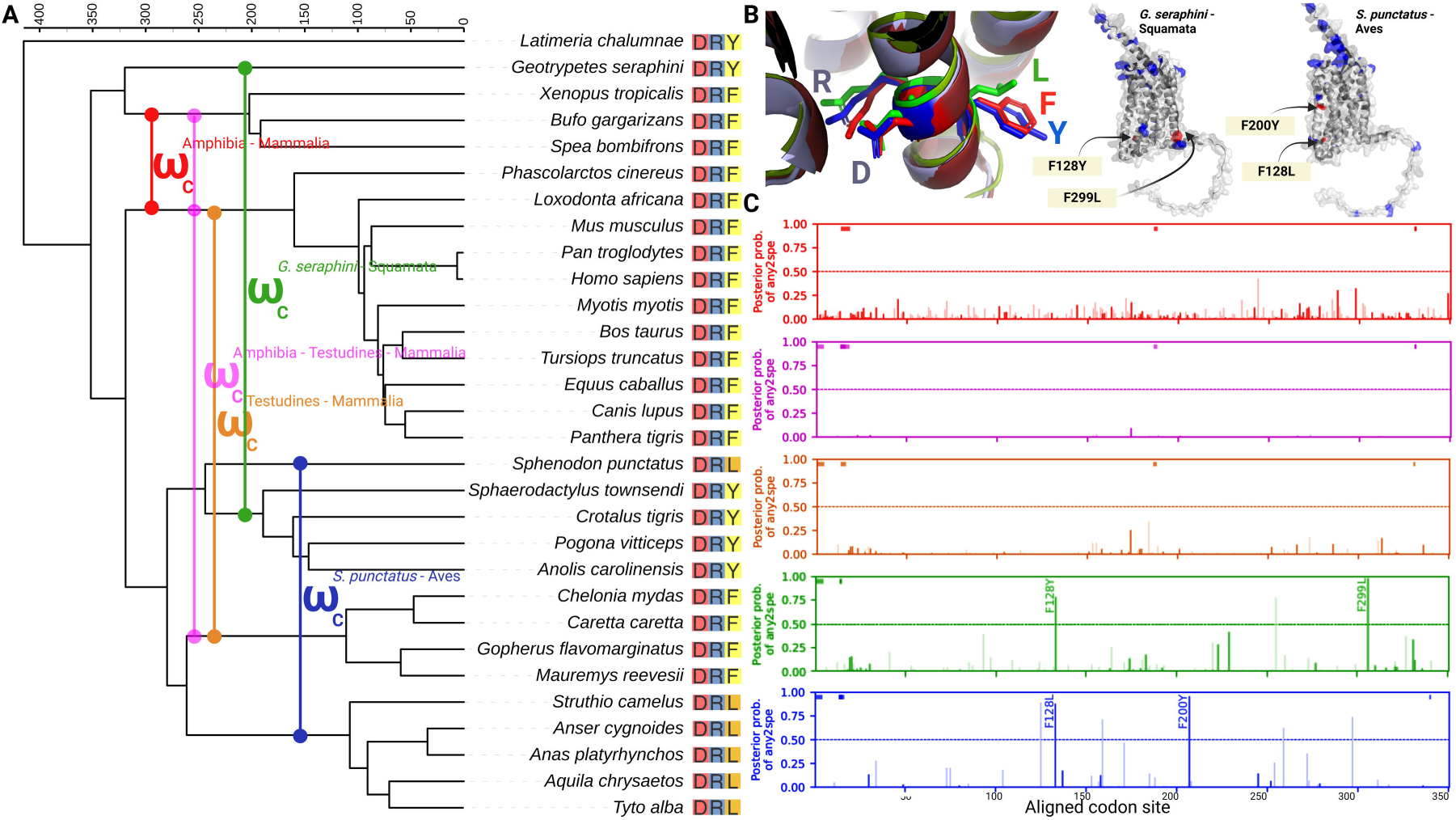
Molecular convergence in vertebrate CXCR6. **A.** The 30 vertebrate species were selected based on DRY/DRF/DRL and evolutionary time. Foreground branches (labelled on the phylogenetic tree with different colours □_C_) are decided based on the DRY/DRF/DRL motif and used to find convergent substitution (any2spe) using CSUBST. **B.** The DRY/DRF/DRL motif change is shown on the 3D structure of CXCR6, whereas the convergent substitutions mapped to human protein structure in "Squamates-like" and "Aves-like" foreground lineages. **C.** The convergent sites are shown for foreground branches with their posterior probabilities of any amino acid to specific amino acid substitution (coloured according to foreground branch combination). The Y-axis represents the site-specific magnitude of convergence in the foreground branches, and the X-axis represents the site numbers in the input alignment. The dark- and light-coloured bars represent the non-synonymous and synonymous substitutions. The time-calibrated phylogenetic tree was downloaded from the Timetree website. The time scale is in million years. PyMol is used for the visualisation of 3D protein structures.

Our short (200 ns) molecular dynamics (MD) simulations suggest that residues from the H3C region interact with G-proteins and the third amino acid residue (DRY/DRF/DRL) differentially influences the interaction frequency of receptors with G-proteins (**Fig. 4**). Coupling probabilities estimated using PRECOGx, between vertebrate CXCR6 protein orthologs and Gi/o was higher than those with other G-protein families **(Supplementary Figure 14, Supplementary Table 8)**. Among the Gi/o proteins, the coupling probabilities were highly heterogeneous for GoA, with Aves having lower probabilities than other vertebrates **(Supplementary Figure 14)**. One major difference in the C-terminal between GoA and GoB occurs at residue 346, with N in GoA vs K in GoB. Replacing the N in GoA with the GoB-like K in MD simulations changed the interaction frequency at several important positions in the H3C region (**Supplementary Figure 15**). Sequence convergence at the functionally important DRF motif, changes in the coupling probabilities estimated using PRECOGx, and differential interactions predicted by MD simulations highlight the importance of the DRF containing the H3C region.

**Fig. 4:**
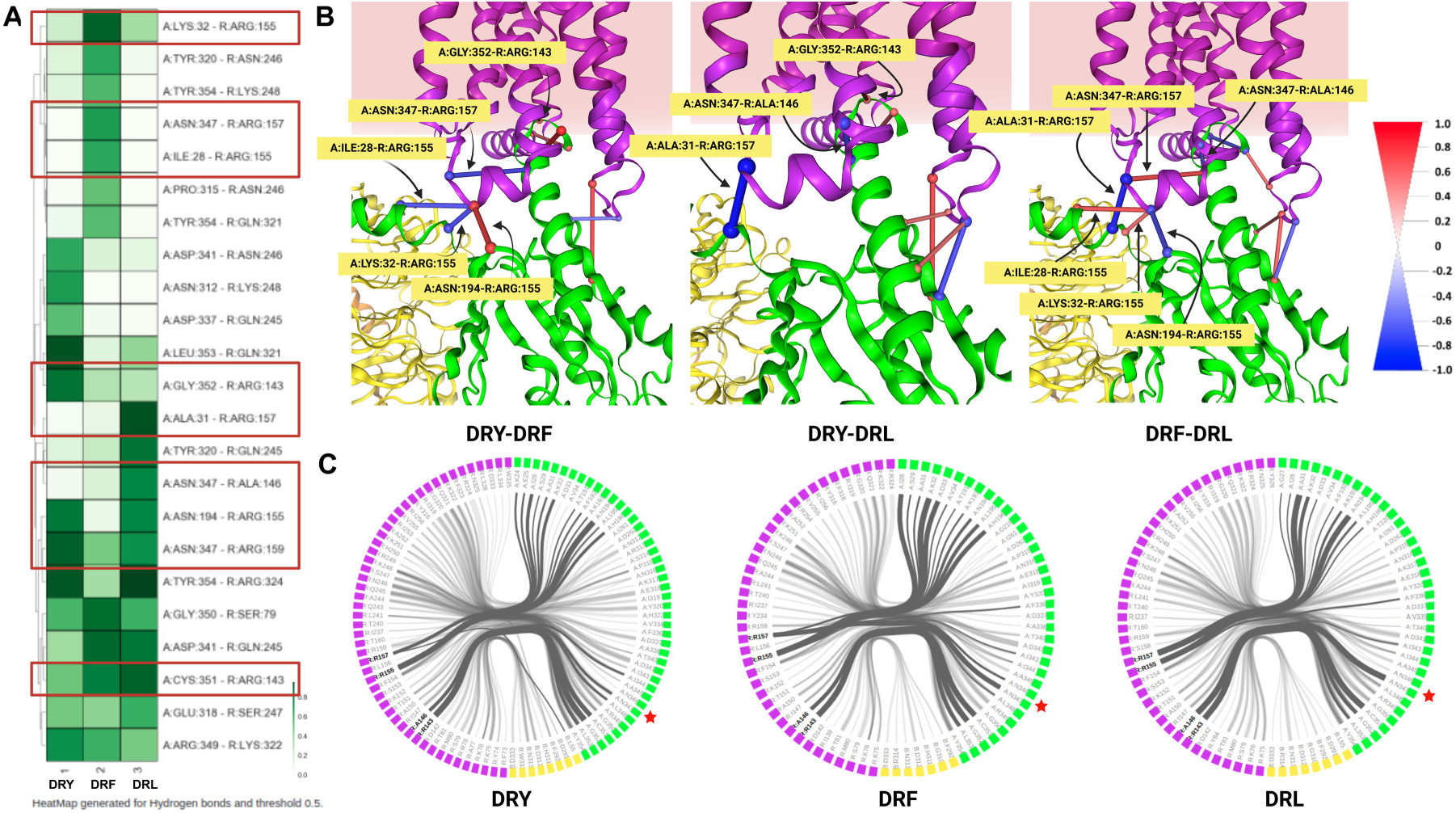
Interaction frequency difference analysis. **A.** The heatmap for hydrogen bonds compares interaction frequency estimated by MD simulations for DRY/DRF/DRL mutations. The frequency difference cutoff was set to 0.5 to show meaningful interaction. Red rectangular boxes show the residues from the H3C region on the heatmap. **B.** The frequency difference between each pair is shown with sticks. The stick cutoff is 0.5. the intensity of colour and the stick thickness are proportional to the average frequency difference between each combination. The colour and thickness scale is shown on the top-right side. **C.** The FlarePlot shows the residue-residue interactions for DRY/DRF/DRL-specific changes. All interactions (Hydrogen bonds, Aromatic, Salt bridges, and Van der Waals) are shown. Pink boxes show the receptor residue, whereas the green and yellow boxes represent the residues from Gα and Gβ G-proteins, respectively. The crucial residues of receptors ARG143, ALA146, ARG155, and ARG157 are shown in bold. The line thickness corresponds to the frequency of residue-residue interaction and the number of times such interaction is found. The asparagine (N), the site where GoA and GoB differ, is shown with a red star.

### Recurrent pseudogenisation of *CXCR6* gene in birds

Of the 36 bird orders (122 species) examined, the *CXCR6* gene has acquired frame-disrupting changes in 23 species from 3 bird orders **(Fig. 5A, Supplementary Table 1)**. In addition to these species that contain gene remnants, the *CXCR6* gene is completely missing from the genomes of 31 species spanning nine orders. In most of these species which are missing the gene, the gene is potentially lost due to segmental deletions, as the flanking exons of the *FYCO1* gene and the intervening intron are always found in the genome assembly. We could verify the reliability of the genome assembly at the *CXCR6* locus using long-read data in the case of chicken, zebra finch, pigeon, downy woodpecker, budgerigar, Northern carmine bee-eater, common swift, Anna’s hummingbird, and Kakapo (**Fig. 6A, Supplementary Figure 16-22**). In contrast to the species with missing *CXCR6*, species with the intact gene at the correct syntenic location, such as mallard (*Anas platyrhynchos*), have clear expression in lung and liver tissue **(Supplementary Figure 23, Supplementary Table 10)**.

**Fig. 5:**
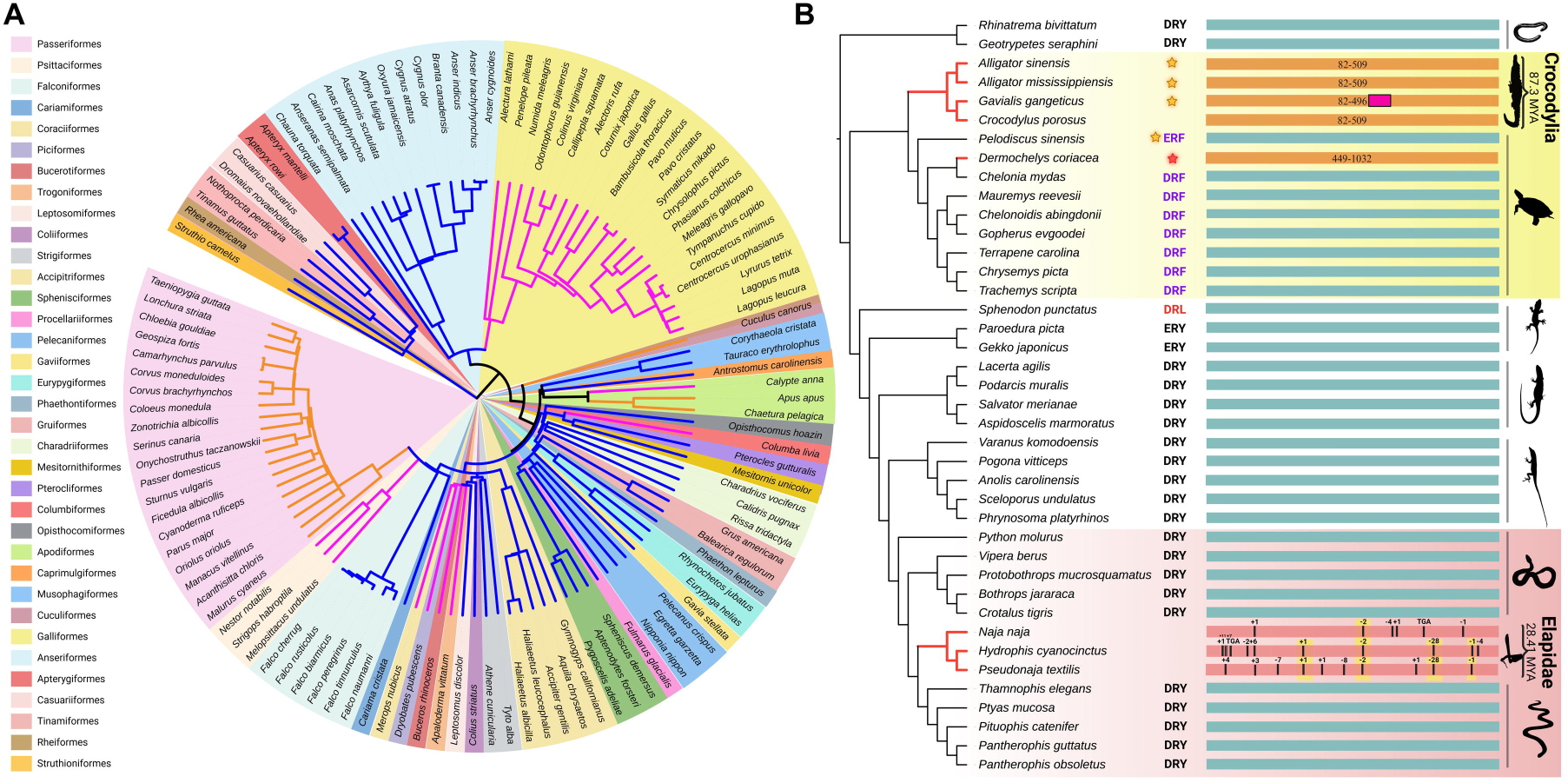
Recurrent gene loss in Sauropsids. **A.** Summary figure of *CXCR6* across different orders of class Aves. The solid blue colour branches of the circular phylogenetic tree represent the intact *CXCR6* gene. The solid orange colour branches indicate loss of the *CXCR6* gene due to frame-disrupting changes, whereas solid pink phylogenetic branches show *CXCR6* gene loss due to segmental deletion. The bird orders are mentioned on the left side, and the phylogenetic tree is coloured accordingly. **B.** Status of *CXCR6* gene in reptiles. Red branches show loss of the *CXCR6* gene, whereas solid black branches indicate intactness of the *CXCR6* gene. The glowing orange star represents relaxed selection in that species using HYPHY. The DRF/DRY/ERY motif in the H3C region is shown for that species. The orange horizontal bar shows partial remains of *CXCR6*. The start and end positions of partial hits are mentioned on the horizontal bars. The pink rectangular box on the top of the orange bar represents the insertion of repeat in the Gharial (*Galivialis gangeticus*). Repeats were identified using RepeatMasker v4.1.5. A light-red horizontal bar with events indicated with black lines represents *CXCR6* gene pseudogenisation events. The yellow highlighted events are the shared inactivating mutations (SIMs). The time- calibrated phylogenetic tree is downloaded from the Timetree website. The iTOLv6 was used to annotate branches of the phylogenetic tree. The species with yellow and red gradient background highlights are used in the selection analyses for that clade. The Silhouette Images are taken from the PhyloPic (https://www.phylopic.org/).

**Fig. 6:**
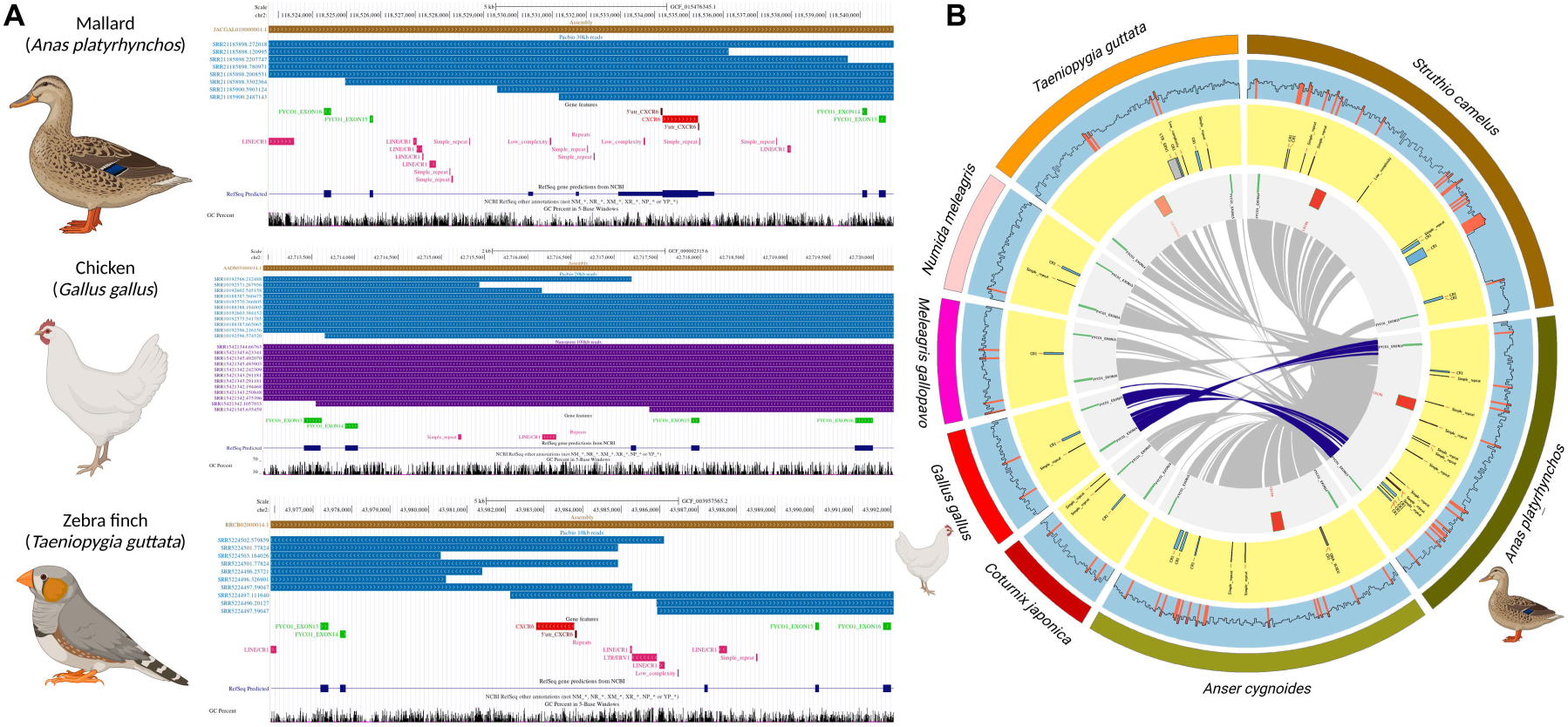
Galliformes specific *CXCR6* gene loss. **A.** Assembly verification at the syntenic location of the *CXCR6* in mallard (top panel), chicken (middle panel), and zebrafinch (bottom panel). Royal blue represents the PacBio reads of longer than 10 Kb, whereas reads in violet are of Nanopore (>100Kb), with their read IDs on the left side. The direction of arrows within reads indicates the read’s orientation. The gene feature tract shows the *FYCO1* gene exons (light green colour), *CXCR6* (red colour), and the 5’ and 3’ UTR region of *CXCR6* (brown boxes). The type of repeats occurring in this region are shown in pink in the Repeats track. **B.** Circos plot summarising GC content, repeat region, and sequence comparison of *CXCR6* region. The line within the sky blue background shows GC content with red-coloured bars. In the yellow track, repeats are shown. The innermost track, *CXCR6* (red coloured), and *CXCR6*-flanking *FYCO1* exons are shown in light green. The grey ribbons indicate the corresponding sequence of the respective species compared to mallard (for chickens, the blue ribbon shows the same).

### **(a)** Galliformes specific deletion

The *CXCR6* gene is not annotated in the chicken or any other galliform genomes available on NCBI RefSeq **(Supplementary Figure 24)**. However, the anseriform genomes consistently annotate *CXCR6* between the 14^th^ and 15^th^ exons of *FYCO1* (**Supplementary Table 2**, **Fig. 6A**). The pairwise genome alignment between mallard and chicken suggests complete gene loss through segmental deletion (**Supplementary Figure 24**). Further, a blast-based search of sixteen galliform genome assemblies failed to identify the *CXCR6* gene or its remnants. Nonetheless, the flanking exons of *FYCO1* could be identified in all the genome assemblies of galliforms. The completeness of the genome assembly at the *CXCR6* locus in the chicken genome was validated using long-read sequencing (PacBio and Nanopore) (**Fig. 6A**). A BLASTn search of short-read (Illumina) databases of galliform birds using the closest outgroup species with intact *CXCR6* (ostrich and mallard) sequence as a query failed to recover an intact gene in these species. The search of short-read datasets has the limitation of not being able to identify genes with extreme GC content. Despite searching the long-read database of chicken using the ostrich and mallard *CXCR6* queries, none of the spurious hits (from other CKRs) covered the entire ORF and were extremely short. Therefore, it is implausible that *CXCR6* has translocated to another genomic region in chicken.

BLASTn search of the RNA-seq database of chickens (*Gallus gallus*) from diverse tissues/cell types, such as CD4, CD8 cells, liver, and T cells, failed to find evidence for *CXCR6* transcripts. The consistent absence of the *CXCR6* gene region across all Galliformes in contrast to its presence in Anseriformes allowed us to deduce the loss of *CXCR6* in Galliformes via segmental deletion. A comparison of the mallard *CXCR6* locus with the corresponding galliform loci found CR1 repeats at the junction of alignment blocks **(Fig. 6B)**. While the GC content at the *CXCR6* locus is heterogenous, regions with extreme GC do not occur within the gene or its immediate vicinity. The boundaries of the segmental deletion in all galliform genomes occur in the same region as chickens, and the approximate deletion size is ∼10.5 Kb compared to mallard **(Fig. 7A, Supplementary Table 11)**. Pairwise dot plots of mallard with galliform and anseriform species depict the contrast and confirm this galliform order-specific segmental deletion **(Fig. 7B)**. Therefore, multiple lines of evidence strongly support the complete loss of *CXCR6* in galliform birds.

**Fig. 7:**
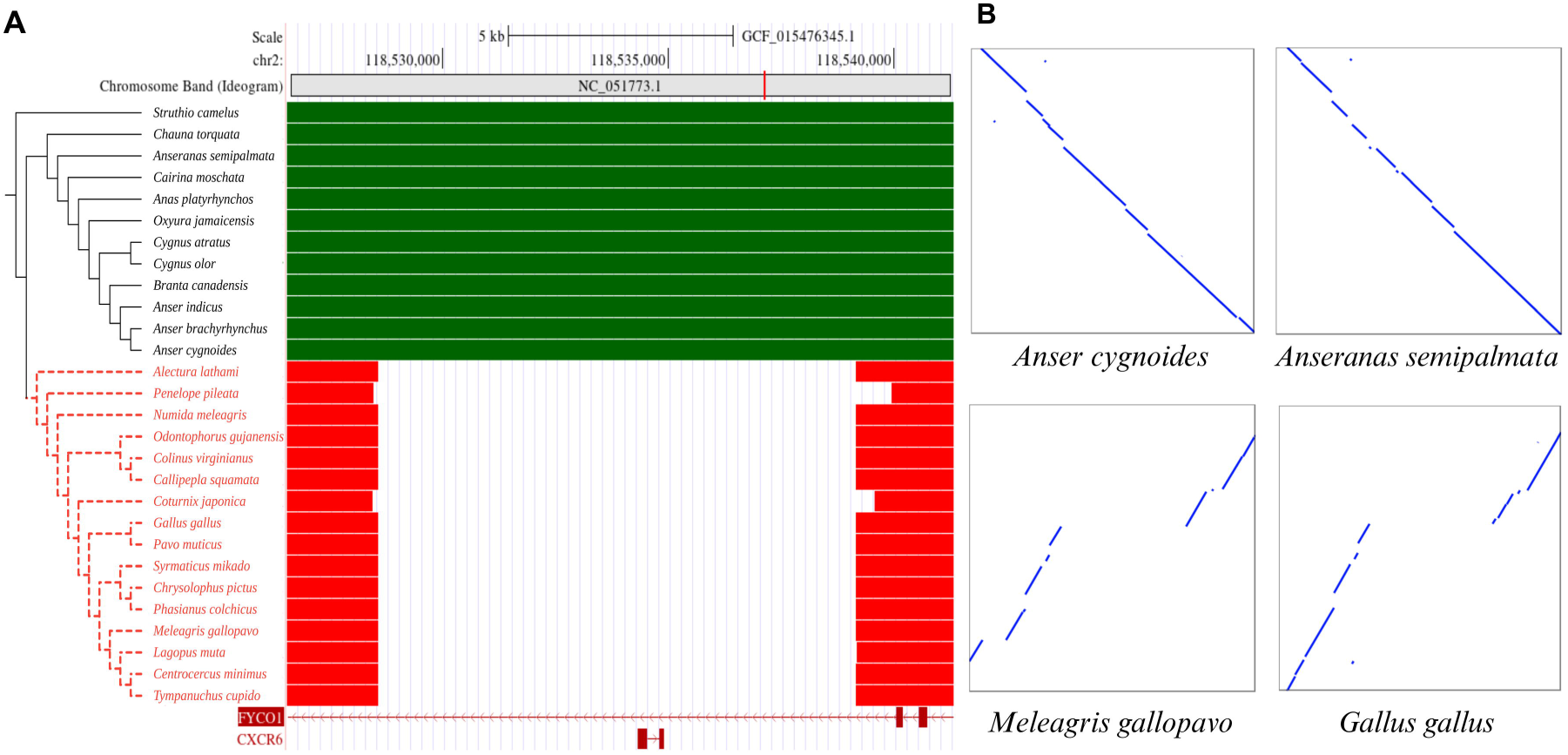
Galliform-specific segmental deletion. **A.** The phylogenetic branches of Anseriformes species are shown with solid black lines, whereas dashed phylogenetic branches represent segmental deletion of the *CXCR6*. The species with green boxes have an intact *CXCR6* gene. In comparison, the species with red colour are Galliformes with *CXCR6* deletion. **B.** Pairwise dot plot comparison to mallard (*Anas platyrhynchos*). The top panel shows the intactness of *CXCR6* in Swan goose (*Anser cygnoides)* and magpie goose *(Anseranas semipalmata)*. In contrast, the bottom panel shows segmental deletion in turkey (*Meleagris gallopavo*) and chicken (G*allus gallus)*. Repeats were identified using RepeatMasker v4.1.5. The UCSC browser is used for visualisation. The iTOLv6 is used to annotate branches of the phylogenetic tree.

### **(b)** Promoter deletion, relaxed selection, and frame-disrupting changes in Passeriformes

The only two passeriform species (white-rumped snowfinch; *Onychostruthus taczanowskii* and common starling; *Sturnus vulgaris*) with *CXCR6* refseq gene annotation are tagged as a low-quality protein hints at the possibility of gene loss in Passeriformes (**Supplementary Table 1**). Although the genomic region containing the *CXCR6* exon is present in the genomes of Passeriformes species, an upstream region of ∼4Kb (compared to the golden eagle (*Aquila chrysaetos*)) which potentially contains regulatory elements is deleted in all Passeriformes **(Fig. 8A, Supplementary Table 11)**. Long-read (PacBio) based validation of the *CXCR6* gene locus rules out the possibility of assembly artifact in the zebra finch (*Taeniopygia guttata*) **(Fig. 6A)**. The high-confidence annotation of gene regulatory elements in the human genome suggests that the missing region in Passeriformes may have contained the *CXCR6* gene enhancers and promoters **(Fig. 8B)**. The bird species with *CXCR6* gene loss, such as zebra finch (*Taeniopygia guttata*), lacks expression **(Supplementary Figure 25, Supplementary Table 10)**.

**Fig. 8:**
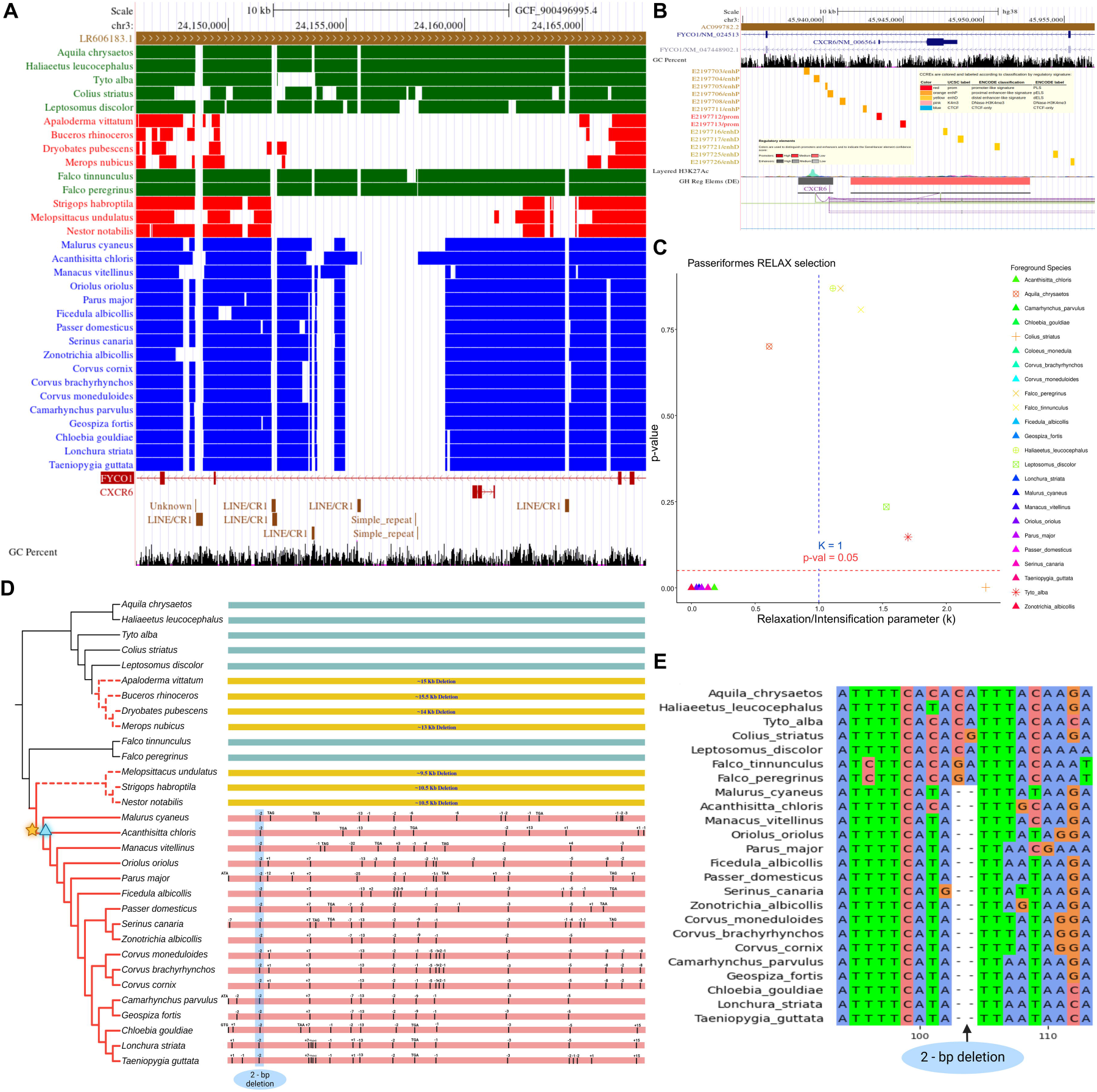
*CXCR6* gene loss in Passeriformes and other independent bird orders. **A.** Deletion size comparison of *CXCR6* gene-containing segments across the Passeriform species compared to the golden eagle (*Aquila chrysaetos*). The species with green boxes have an intact *CXCR6* and do not show the deletion. In comparison, the species with red colour have *CXCR6* deletion (the large gap between the boxes). The species with blue boxes has a deletion of the promoter/enhancer region of the *CXCR6* gene in Passeriformes. **B.** The gene regulatory regions, such as promoter/enhancer of the *CXCR6* gene in humans, are shown. UCSC browser is used for visualisation (http://genome.ucsc.edu). **C.** Relaxed selection in Passeriformes. The X-axis represents the k-value, and the y-axis represents the p-value obtained using the RELAX model of HYPHY. Filled triangles represent the Passerine species. **D.** Red-colored solid branches represent the loss of the *CXCR6* gene due to frame-disrupting changes in the phylogenetic tree. The events are shown on top of a single exonic *CXCR6* gene with black lines at corresponding locations. The branches with a dashed red colour represent species with segmental deletion. The approximate size of segmental deletion is mentioned on the yellow horizontal bars. The glowing orange star and sky blue triangle on the phylogenetic tree represent relaxed selection using HYPHY and codeML, respectively. The iTOLv6 was used to annotate branches of the phylogenetic tree. **E.** The shared inactivating mutation of two bp deletions of CA across all the Passeriformes.

Strong signatures of relaxed selection (HYPHY RELAX: FDR-corrected p-value < 0.05; k-value < 1 and PAML’s codeML) are pervasive among passeriform birds **(Fig. 8C, Supplementary Table 12)**. In contrast, relaxed selection signatures are not found in closely related bird species. The Passeriform birds share multiple frame-disrupting changes validated using short-read sequencing **(Fig. 8D)**. The two-base deletion of CA at 83-84^th^ position and the seven-base insertion of TGATGGT at the 202^nd^ position is shared by most Passeriformes **(Fig. 8E)**. The two-base deletion of GT at 413-414^th^ position is shared among Passeriform species except for golden-collared manakin (*Manacus vitellinus*). The signatures of relaxed selection, multiple frame-disrupting changes, lack of gene expression, and the putative loss of the gene regulatory region in Passeriformes conclusively establish gene loss and may have resulted from the deletion of the promoter region.

### **(c)** Independent order-specific gene loss

Similar to the complete loss of the *CXCR6* gene in galliform species, our search of bird genomes and short-read datasets is indicative of independent segmental deletions of ∼13-15 Kb in bar- tailed trogon (*Apaloderma vittatum*; Trogoniformes), rhinoceros hornbill (*Buceros rhinoceros*; Bucerotiformes), downy woodpecker (*Dryobates pubescens*; Piciformes), and northern carmine bee-eater (*Merops nubicus*; Coraciiformes) **(Fig. 5A**, **Fig. 8, Supplementary Table 11)**. Among Psittaciformes species, evidence of segmental deletions is found in the budgerigar (*Melopsittacus undulatus*; ∼9.5 Kb), kakapo (*Strigops habroptila*; ∼10.5 Kb), and kea (*Nestor notabilis*; ∼10.5 Kb) **(Supplementary Figure 19-20)**. Shorter segmental deletions that span the entire coding region are found in domestic pigeon (*Columba livia* ∼6 Kb; Columbiformes) and common cuckoo (*Cuculus canorus* ∼8.5 Kb; Cuculiformes) **(Supplementary Figure 26)**. Genome assembly validation using long-read sequencing in downy woodpecker, northern carmine bee-eater, budgerigar, kakapo, common swift, and pigeon rules out the possibility of assembly artefacts **(Supplementary Figure 16-22)**. Gene loss due to both segmental deletion and frame-disrupting changes can be seen among species of Apodiformes with (1) Galliform-like complete gene loss through segmental deletion in anna’s hummingbird (*Calypte anna*), (2) Passeriform-like gene loss (deletion of upstream regulatory regions and frame-disrupting changes) occurs in the common swift (*Apus apus*) and chimney swift (*Chaetura pelagica*) **(Supplementary Figure 27)**. Hence, loss of the *CXCR6* gene through segmental deletion and/or frame-disrupting changes is frequent among bird species.

### *CXCR6* gene loss in class reptilia

Crocodilians are the closest living relatives of birds (class Aves). The annotation of the *CXCR6* gene is missing for Crocodilia. Further, the *CXCR6* gene annotation is tagged as a low-quality protein in the eastern brown snake (*Pseudonaja textilis*) of the Elapidae family of snakes. Our search of the genomes and short-read datasets recovered remnants of the *CXCR6* gene in Crocodilia and identified shared inactivating mutations (SIMs) in Elapidae snakes **(Fig. 5B)**. Based on the patterns of lineage-specific gene loss, the approximate time (in million years ago, MYA) of *CXCR6* pseudogenisation is ∼87.3 MYA and ∼28.41 MYA in the Crocodilian order and Snakes Elapidae family, respectively.

### **(a)** Fragmentary *CXCR6* remnants establish gene loss in Crocodilia

Genomes of all four Crocodilia species, namely, the American alligator (*Alligator mississippiensis*), Chinese alligator (*Alligator sinensis*), saltwater crocodile (*Crocodylus porosus*), and gharial (*Gavialis gangeticus*), have only partial remains of *CXCR6* gene at the syntenic location **(Supplementary Table 13)**. Searching the short-read datasets using the coding sequences of Chinese softshell turtle (*Pelodiscus sinensis*) and green sea turtles (*Chelonia mydas*) *CXCR6* gene as queries recovered the same remnants. Assembly verification using long-read (PacBio) sequencing eliminates the possibility of assembly artifacts **(Supplementary Figure 28-33)**. Moreover, several repetitive sequences are found in the vicinity of *CXCR6* remnants. In the gharial (*Gavialis gangeticus*), a simple repeat is inserted into the truncated *CXCR6* gene sequence **(Fig. 5B** and **Supplementary Figure 30**). Although we found strong evidence of relaxed selection in the Chinese alligator (*Alligator sinensis*), American alligator (*Alligator mississippiensis*), and the gharial (*Gavialis gangeticus*), no such signatures could be identified in the saltwater crocodile (*Crocodylus porosus*) **(Fig. 5B, Supplementary Table 12)**. The genome assembly and short-read dataset of leatherback sea turtles (*Dermochelys coriacea*; Testudine) contains only a partial sequence of the *CXCR6* gene, suggesting an independent gene loss in this species. Long-read sequencing of the *CXCR6* locus in the leatherback sea turtle validates the genome assembly (**Supplementary Figure 33**).

### **(b)** SIMs in the Elapidae family of snakes

The BLASTn search of the genomes of the Indian cobra (*Naja naja*), blue-banded sea snake (*Hydrophis cyanocinctus*), and eastern brown snake (*Pseudonaja textilis*) using the *CXCR6* gene of western terrestrial garter snake (*Thamnophis* elegans; ENSTELG00000009216) as query found truncated sequences at the syntenic location in all three species. Short-read sequencing supports all the frame-disrupting changes identified among Elapidae snakes (**Fig. 5B**). We found a two-base (AA nucleotide) SIM at position 552-553^rd^ across the Elapidae family. Another SIM of 28-base deletion from the 773-800^th^ position was found in the blue-banded sea snake (*Hydrophis cyanocinctus*) and eastern brown snake (*Pseudonaja textilis*) **(Fig.5B)**. The *CXCR6* gene is intact and expressed in the green sea turtle (*Chelonia mydas*; spleen) **(Supplementary Figure 34)**. However, the RNA-seq datasets screened lack evidence of gene expression in the Chinese alligator (*Alligator sinensis*; spleen), blue-banded sea snake (*Hydrophis cyanocinctus*; venom gland), and Indian cobra (*Naja naja*; venom gland) **(Supplementary Figure 35-37)**.

### Pigeon lung has lower CD8 T_RM_ cell abundance than mallard

We hypothesized that the lineage-specific loss of *CXCR6* may have consequences for the localization of CD8 T_RM_ cells to the lungs, which could have implications for mucosal immune defense. Consistent identification of CD8 T_RM_ cells among species with (mallard) and those without (chicken, pigeon) *CXCR6* is based on marker genes (*CD69*, *ITGAE*, and *ITGA1*). However, in the case of chicken and other galliforms, the *ITGAE* (*CD103*) gene is not annotated on RefSeq, in contrast to the consistent annotation among anseriform species **(Supplementary Figure 38)**. The genome assembly, short and long-read sequencing-based validation, and lack of gene expression for all exons in diverse tissues confirm the loss of the *ITGAE* gene in chicken (**Supplementary Figure 39**). A few exons of *ITGAE* were expressed in chickens and were recognized as long non-coding RNA (**Supplementary Figure 39**, **Supplementary Table 14**). Putative candidate bird species with *ITGAE* gene loss identified by TOGA (which needs further validation) overlapped several species with *CXCR6* gene loss (**Supplementary Figure 40, Supplementary Table 14**).

Concurrent loss of *CXCR6* and *ITGAE* in chicken raises the question of T_RM_ cell identity and simultaneously makes it challenging to pinpoint T_RM_ cells in single-cell transcriptomes. Moreover, single-cell transcriptomes of birds are still scarce and not always comparable due to methodological differences. Fortunately, a single experimental study of lung single-cell transcriptomes of mallard (intact *CXCR6*) and pigeon (*CXCR6* gene loss) is available on the SPEED database. These single-cell RNA-seq datasets allowed the evaluation of differences in T_RM_ cell abundance between mallard and pigeon lungs. The SPEED database cell-type annotation identifies the *ITGAE*-expressing cells as a sub-population of fibroblasts and ciliated cells. In contrast to the small fraction of *ITGAE*-expressing cells, *ITGA1*-expressing cells are more broadly expressed across different cell types. We looked for cells that co-express *ITGAE* and *ITGA1* to narrow down the T_RM_ cells. The mallard (183 cells) single-cell transcriptome appears to have more T_RM_ cells than pigeons (2 cells) **(Supplementary Figure 41-42)**. This observation suggests that species lacking *CXCR6* exhibit diminished populations of CD8 T_RM_ cells compared to those with intact *CXCR6*. Among the six chicken lung single-cell transcriptomes compared, the expression patterns of *CD69* and *ITGA1*-expressing cells are heterogeneous. Moreover, *CD69*-expressing cells consistently do not form clusters and represent a small fraction of the cells. However, *ITGA1*-expressing cells are abundant and form clusters in all six conditions **(Supplementary Figure 43)**. Our detailed analysis of the limited single-cell lung transcriptomes suggests a role for species-specific *CXCR6* gene presence/absence in determining T_RM_ cell identity, abundance, and localization.

## Discussion

Our study explores the evolution and diversification of the *CXCL16*-*CXCR6* genes from over 400 vertebrate genomes. Evolutionary trace analysis of the CXCR6 DRF motif identified heterogeneity among key vertebrate lineages with evidence of recurrent gene loss in the Sauropsid lineages concurrent with the loss of *CXCL16*. The DRF motif is restricted to mammals, turtles, and frogs, while the DRY motif, typical in other CKRs, is found in snakes and lizards. Birds exhibit the DRL motif, except for the harpy eagle, which possesses DRF. Convergent amino acid substitutions were identified at the DRF motif and two other positions in Aves- and Squamate-like lineages. MD simulations reveal that DRY/DRF/DRL site variations affect G-protein interaction. The Aves-like DRL displayed reduced GoA G-protein coupling probability, as estimated using a machine learning-based tool PRECOGx. Our comparative study identified recurrent loss of the *CXCR6* gene in 10 out of 36 bird orders attributed to segmental deletion and/or frame-disrupting changes. The species from the order Crocodilia retain partial gene relics at the syntenic location, while the Elapidae family of snakes lost *CXCR6* due to frame-disrupting changes. A prominent phenotype associated with the absence of *CXCR6* is a decrease in airway CD8 T_RM_ cells (Wein et al. 2019). Interestingly, species lacking the *CXCR6* gene show decreased T_RM_ cells, as evidenced by comparisons of single-cell RNA-seq datasets. The concurrent loss of two core T_RM_ cell marker genes (*CXCR6* and *ITGAE*) in chicken may require revisiting the definition of T_RM_ cells and their abundance as T cells differ between birds and mammals (Burkhardt et al. 2022).

### Comparative analysis of *CXCR6* unveils lineage-specific diversification

The distinctive features of the CXCL16-CXCR6 axis include the receptor’s unique DRF motif, the presence of its ligand in both membrane-bound and soluble forms, and the lack of promiscuous interactions with other CKRs/chemokines. The evolutionary expansion of the CKRs gene family is driven by whole and segmental genome duplication events and substitutions in key residues (Nomiyama et al. 2010; Hughes and Nibbs 2018). The combination of gene collinearity and sequence similarity analysis allowed us to confidently identify vertebrate *CXCR6* and *CXCL16* orthologs. While previous comparative studies of *CXCL16-CXCR6* found minor variation among species of the murine and primate orders (Xu et al. 2018; He et al. 2020), our study highlights the lineage-specific changes in CXCL16-CXCR6 in vertebrates. The basic residues in CXCL16 and acidic residues in the N-terminus of CKRs (including CXCR6) are important for ligand-receptor interaction and downstream signalling (Monteclaro and Charo 1996, 1997; Shimaoka et al. 2004b; Petit et al. 2008; Wasilko et al. 2020; Zhang et al. 2021). However, not all N-terminus residues are critical for productive interaction (Petit et al. 2008). We found a reduction in N-terminus acidic residues in birds and turtles compared to other vertebrates. The N-terminus regions of other CKRs are heterogeneous, with changes in the abundance of acidic residues (Szpakowska et al. 2012).

The convergent substitution F200Y (F^5.47^) found in Aves-like lineages interacts with the CWxP motif of CKRs (Nomiyama and Yoshie 2015). In some CKRs, the F → L substitution was reported in tetrapods and teleosts (Nomiyama and Yoshie 2015). The steric hindrance mutation at this site resulted in constitutive G_αi_ activity of CCR5 (Steen et al. 2014). The other convergent substitution, F299L (F^8.54^), is in the C-terminal and close to the NPxxY(X)_5,6_F motif. These convergent substitutions may affect the signalling pathway and intensity, with the DRF motif being an adaptation for adhesion (Koenen et al. 2017). Structural and functional studies have found that the H3C region is involved in G-protein signalling in Class A GPCRs (Zhou et al. 2019; Jones et al. 2020). The DRF motif and H3C region are involved in the selective use of G_i/o_ proteins. Specifically, the third position affects G_o_ G-protein coupling (Singh et al. 2015). The substitution at the third residue of DRF (DRF or DRL) may not significantly impact the interaction with G-protein signalling as both have large, nonpolar side chains. Similar to these inter-species differences, the Corazonin GPCR receptor gene in the tick parasite has splice variants CRZ-Ra and CRZ-Rb with DRF and DQL motifs, respectively (Hauser et al. 2023). These splice variants are reported to differ in G-protein coupling, highlighting the importance of this motif. While our MD simulation found a difference in interaction frequency between DRY and DRF/DRL, no such difference was found between DRF and DRL at a threshold of 0.5. Therefore, species with DRF/DRL containing CXCR6 may have impaired chemotaxis and adaptation towards adhesion. Moreover, the higher probability of coupling between CXCR6 and G_i/o_, as predicted by PRECOGx, could be explained by the role of G_i/o_ in immune system functions such as migration and cellular adhesion of immune cells (Neptune and Bourne 1997; Villaseca et al. 2022). Earlier studies have used CCR6 (Singh et al. 2015) and CX3CR1 (Koenen et al. 2017) to study the importance of the DRF motif in CXCR6. Future studies can use CXCR6 orthologs from different species with heterogeneity at the DRF motif.

### Recurrent loss of *CXCR6* among Sauropsid lineages

Our study confirms the presence of *CXCR6* in elephant sharks, with subsequent gene loss in ray-finned fish such as zebrafish (Bajoghli 2013; Nomiyama and Yoshie 2015). However, *CXCR6* is intact in coelacanth, lungfish, and amphibians (with an MRCA (Most Recent Common Ancestor) of ∼350 MYA) (Boehm et al. 2012). The *CXCR6* gene is widely conserved among mammals and functionally characterized mainly in human and murine models. Gene perturbation studies have established that *CXCR6* is vital for retaining CD8 T_RM_ cells at mucosal sites and has a key role in immune defence (Mueller and Mackay 2016; Zaid et al. 2017; Wein et al. 2019). A crucial evolutionary transition in function and diversification of *CXCR6* appears to have occurred with the emergence of birds and reptiles. Among reptiles (with an MRCA of ∼300 MYA), we found the loss of *CXCR6* in Crocodilia and an independent loss in the Elapidae family of snakes, while it is intact in lizards and turtles. The co-expression of sialic acid α2,3-Gal and α2,6-Gal receptors in snake respiratory and digestive tracts has been linked with potential susceptibility to avian and human influenza A viruses (e Silva et al. 2024). Similarly, experimental evidence of susceptibility to influenza in alligators (Temple et al. 2015) and prevalence in crocodiles (Davis and Spackman 2008) is notable, given their phylogenetic proximity to birds (Short et al. 2015).

In the bird lineage, we establish the recurrent loss of the *CXCR6* gene in 10 out of 36 examined bird orders via segmental deletion and/or frame-disrupting changes. Overall, in species with intact *CXCR6,* we can recover *CXCL16* when the genome assembly quality is good. Therefore, we don’t anticipate a role for alternative ligands.

Our analysis of single-cell RNA-seq datasets found a difference in T_RM_ cell abundance between bird species with intact (mallard) and lost (pigeon) *CXCR6*. While we cannot rule out the possibility of other inter-species genomic differences altering the T_RM_ cell abundance, future single-cell studies with large sample sizes in multiple species can better evaluate the role of *CXCR6*.

The *CXCR6* gene loss in all galliform species, compared to an intact gene in all anseriform species, is a striking pattern. In ducks, CD8 T cells demonstrate efficacy against AIV (Dai et al. 2022), while in chickens, T cells play a crucial role in combating AIV infection (Dai et al. 2019). Furthermore, the adoptive transfer of AIV-specific CD8 T cells into chickens reduces morbidity and improves survival against subsequent lethal AIV challenges (Seo and Webster 2001; Seo et al. 2002). The loss of T_RM_ cell-specific genes such as *CXCR6* and *ITGAE* in chickens could have implications for dealing with pathogen invasions, susceptibility to infection, and limiting transmission (Uddbäck et al. 2023). Similarly, galliform-specific loss of mucosal defence genes (Salve et al. 2023) and other immune genes (Sharma et al. 2020, 2022; Krchlíková et al. 2023) have been identified. Together, these findings suggest a very different mucosal defence system in chickens, potentially explaining their differing susceptibility to AIV and guiding the development of a T_RM_ cell-based vaccine.

Loss-of-function mutations in chemokines/CKRs such as *CCR5* (Albalat and Cañestro 2016) and *CCL8* (van der Loo et al. 2016) lead to adaptive benefits. Similarly, the *CXCR6* deficiency is beneficial as it improves host defence against tuberculosis and influenza using alternative receptors (Ashhurst et al. 2019) and has a cardio-protective function (Zhao et al. 2013). The convergent and independent loss of *CCL27*, which interacts with skin-infiltrating T cells expressing a specific *CCR10*, has been associated with changes in skin physiology (Lopes-Marques et al. 2018, 2019; Espregueira Themudo et al. 2020). The recurrent loss of *CCL27* has been hypothesized to result from "an attenuated inflammatory response in the skin", and a similar scenario involving compromised mucosal defence may explain *CXCR6* and *CXCL16* gene loss. Additionally, the knockout of *XCR1*, a chemokine receptor, has been shown to affect IBV vaccine efficacy in chickens (Glendinning et al. 2024). Since chickens are naturally knocked out for *CXCR6*, a similar impact on vaccine effectiveness, especially for mucosal pathogens, is plausible. *CXCR6* recruits immune cells and regulates immune responses similar to *XCR1*. Therefore, the absence of *CXCR6* may alter immune cell dynamics and influence vaccine efficacy.

### Implications of gene loss for vaccine development

The mucosa is routinely used to vaccinate many bird species as a cost-effective and established method (Nochi et al. 2018). However, vaccines developed for one species might not be effective for another due to differences in the immune gene repertoire. *CXCR6* deficiency in mice leads to impaired cancer vaccine efficiency (Karaki et al. 2021). Therefore, lineage-specific loss of genes like *CXCR6* would similarly lead to reduced vaccine effectiveness. Similar to vaccines, the efficiency of other therapeutic interventions, such as chimeric antigen receptor (CAR)-T cell therapy, can also differ between species. For instance, improved antitumoral activity can be achieved by adding *CXCR6* (Lesch et al. 2021; Mabrouk et al. 2022). Hence, species with *CXCR6* gene loss can be targeted for improvised immunotherapies and vaccination. Nonetheless, the effect of therapeutically introducing lost genes into a species to improve vaccine efficiency needs further evaluation.

## Conclusion

The interaction of chemokines with chemokine receptors (CKRs) guides immune cells to target sites, impacting immune defence efficacy. The unique DRF motif in CXCR6 characterizes the specificity of the CXCL16-CXCR6 ligand-receptor axis. The DRF motif is crucial in cell adhesion and interaction with G-proteins. Our evolutionary trace analysis provides insight into variations in the DRF motif across vertebrate *CXCR6* 1-to-1 orthologs and other lineage-specific convergent substitutions, influencing receptor interaction frequencies with G-protein. The CXCL16-CXCR6 ligand-receptor duo is vital for the localization and maintenance of tissue-resident memory T-cells (T_RM_), of which *CXCR6* serves as the core T_RM_ marker gene along with *CD69*, *ITGAE*, and *ITGA1*. T_RM_ cells are critical for effective immunity against pathogenic infections such as influenza, as deficiencies in *CXCR6* lead to impaired immune responses. Here, by doing a comparative genomic study for *CXCR6* gene presence/absence, we establish recurrent loss of the *CXCR6* gene in Saurpsid lineages via segmental deletion and/or frame-disrupting changes. The reduced T_RM_ cell abundance in species which have lost the *CXCR6* gene further emphasizes its importance in immune cell localization and function. Our results highlight the evolution and diversification of CXCL16-CXCR6 in Sauropsids through the repeated concurrent loss of *CXCR6* and *CXCL16*. Furthermore, species with *CXCR6* gene loss may be targeted for therapeutic approaches to enhance T_RM_ retention and improve vaccination outcomes.

## Acknowledgements

This article was funded by the Department of Biotechnology, Ministry of Science and Technology, India (grant no. BT/11/IYBA/2018/03). The Science and Engineering Research Board (grant no. ECR/2017/001430) provided funds for procuring computational resources (i.e., Har Gobind Khorana Computational Biology cluster) used. We thank the Council of Scientific & Industrial Research for a fellowship to BGS and the Ministry of Human Resource Development for a fellowship to SS. We used BioRender (https://biorender.com) to arrange figures and images of species.

## Authorship Contribution Statement

BGS: Conceptualization, Investigation, Data curation—formal analysis, Writing—original draft, Writing—review, editing, and revision; SS: Investigation, Data curation—formal analysis, Writing-original draft; NV: Conceptualization, Investigation, Data curation—formal analysis, Project Administration/oversight, Writing—original draft, Writing—review, editing, and revision.

## Data archiving

Data and relevant code for this research work are stored in GitHub: https://github.com/CEGLAB-Buddhabhushan/CXCL16-CXCR6.git.

