## Supplementary Figures for "Evolutionary diversity of CXCL16*-*CXCR6: Convergent Substitutions and Recurrent Gene Loss in Sauropsids"

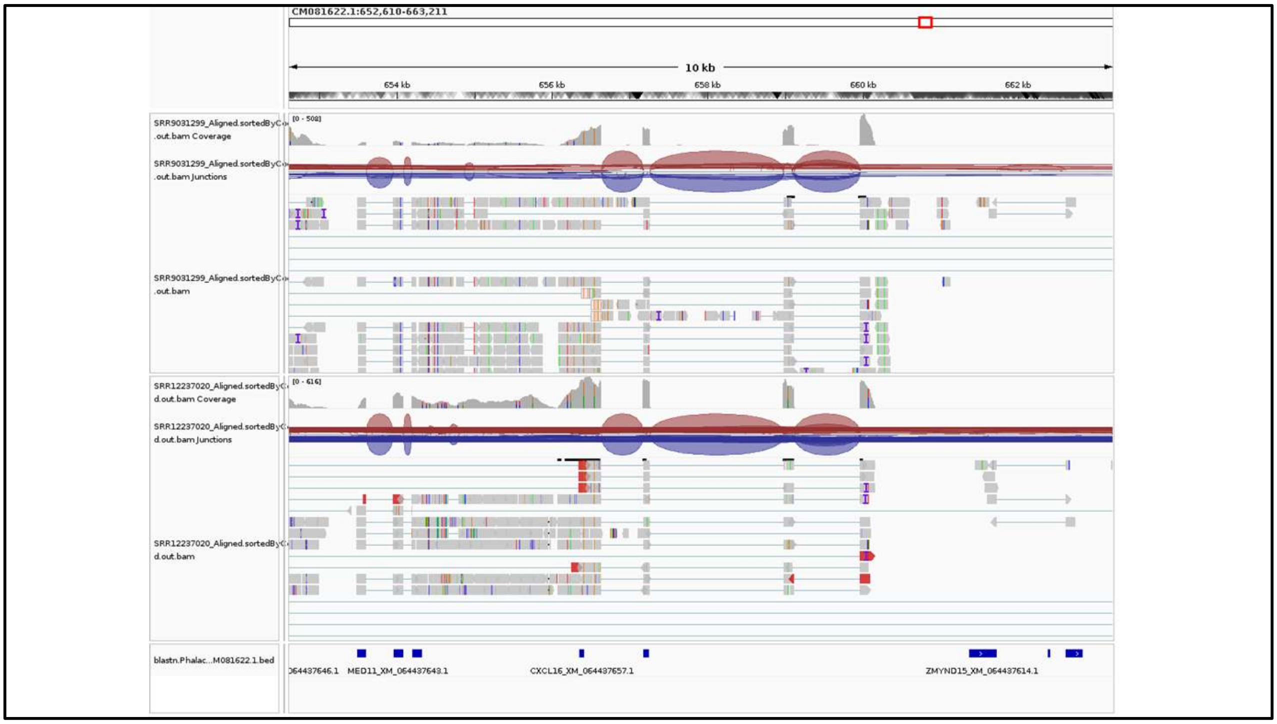


**Supplementary Figure 1:** Expression of *CXCL16* in ostrich. The *CXCL16* is expressed in ostrich at its syntenic locus. The *CXCL16* flanked by *MED11* and *ZMYND15* on choromosome CM081622.1. The SRA study (SRR9031299 and SRR12237020) mapped to reference genome using STAR read mapper.

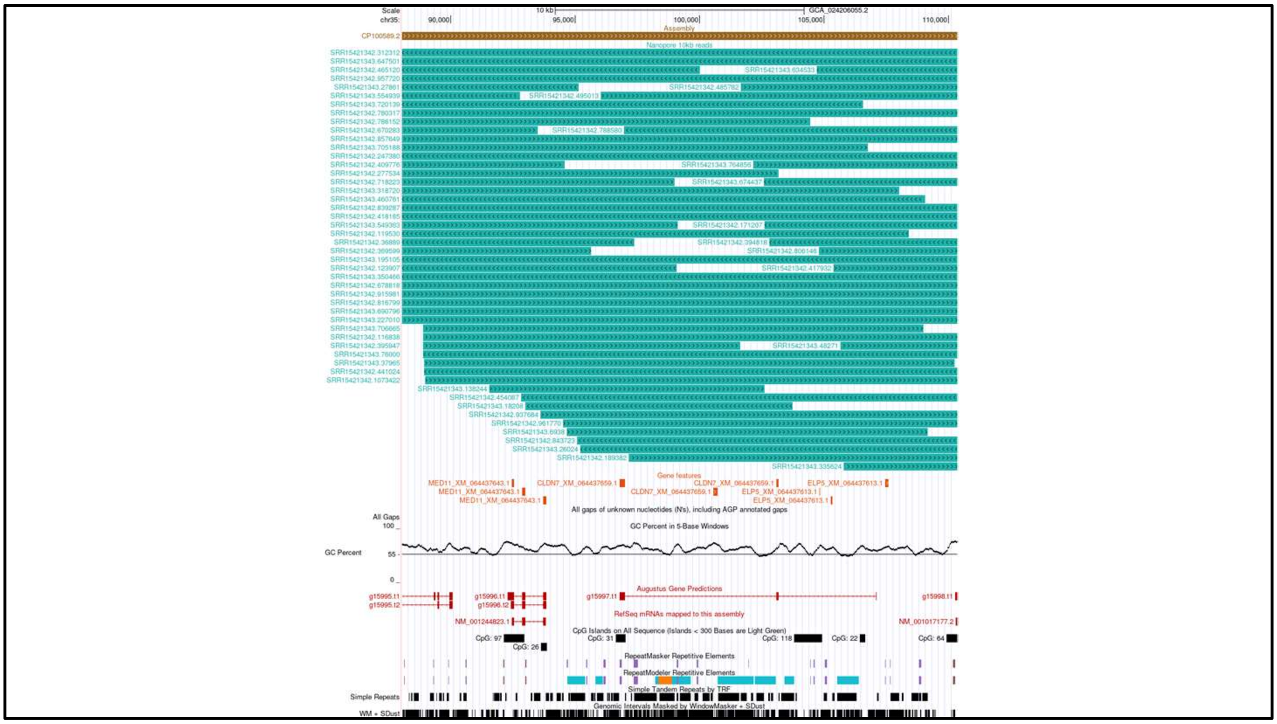


**Supplementary Figure 2:** Assembly verification at the *CXCL16* syntenic locus in chicken. The top track displays nanopore reads longer than 10 kb with their corresponding read IDs, spanning the region between the *MED11* and *CLDN7* genes. The Gene feature track highlights BLASTn hits for *MED11* and *CLDN7* in orange. No BLASTn hits were detected for *CXCL16* at its syntenic locus. The UCSC GC content track illustrates the GC content across this region.

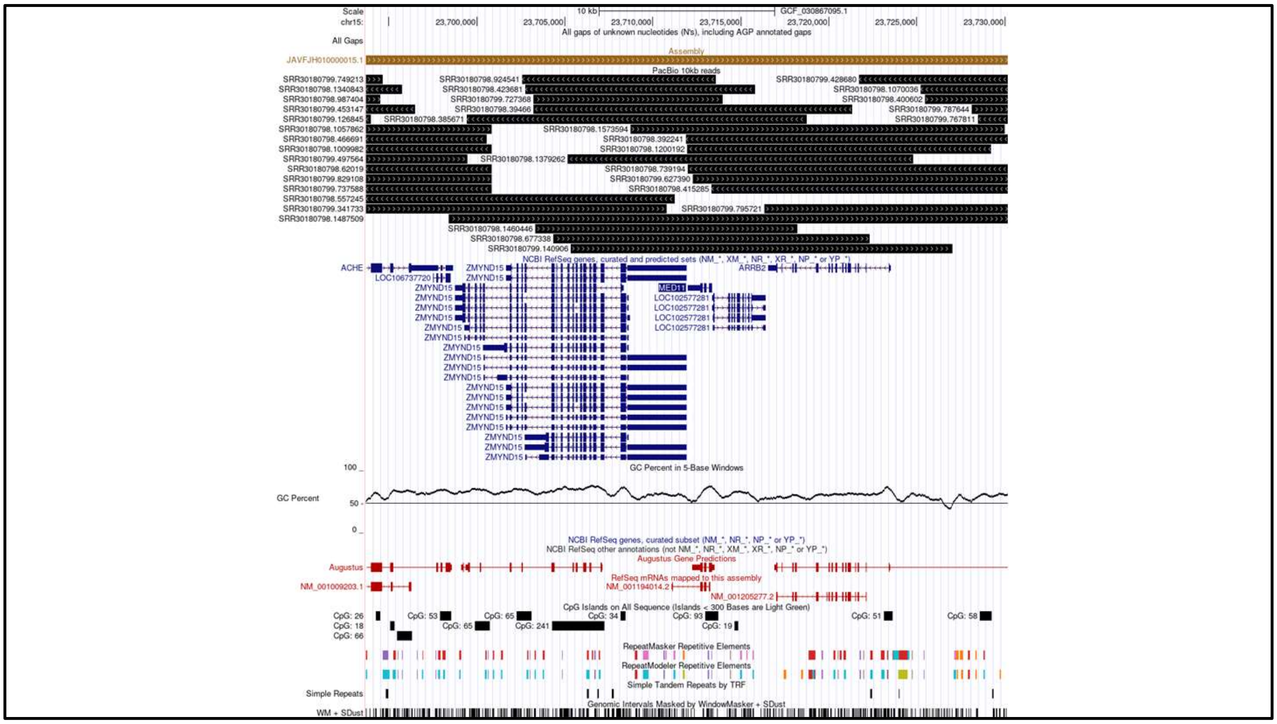


**Supplementary Figure 3:** Assembly verification at the *CXCL16* syntenic locus in *Alligator_mississippiensis.* The top track displays PacBio reads longer than 10 kb with their corresponding read IDs, spanning the region between the *MED11* and *ZMYND15* genes on chromosome 15 (NC_081838.1). The NCBI's Gene feature track shows the annotation for *ZYMND15* and *MED11*. The UCSC GC content track illustrates the GC content across this region.

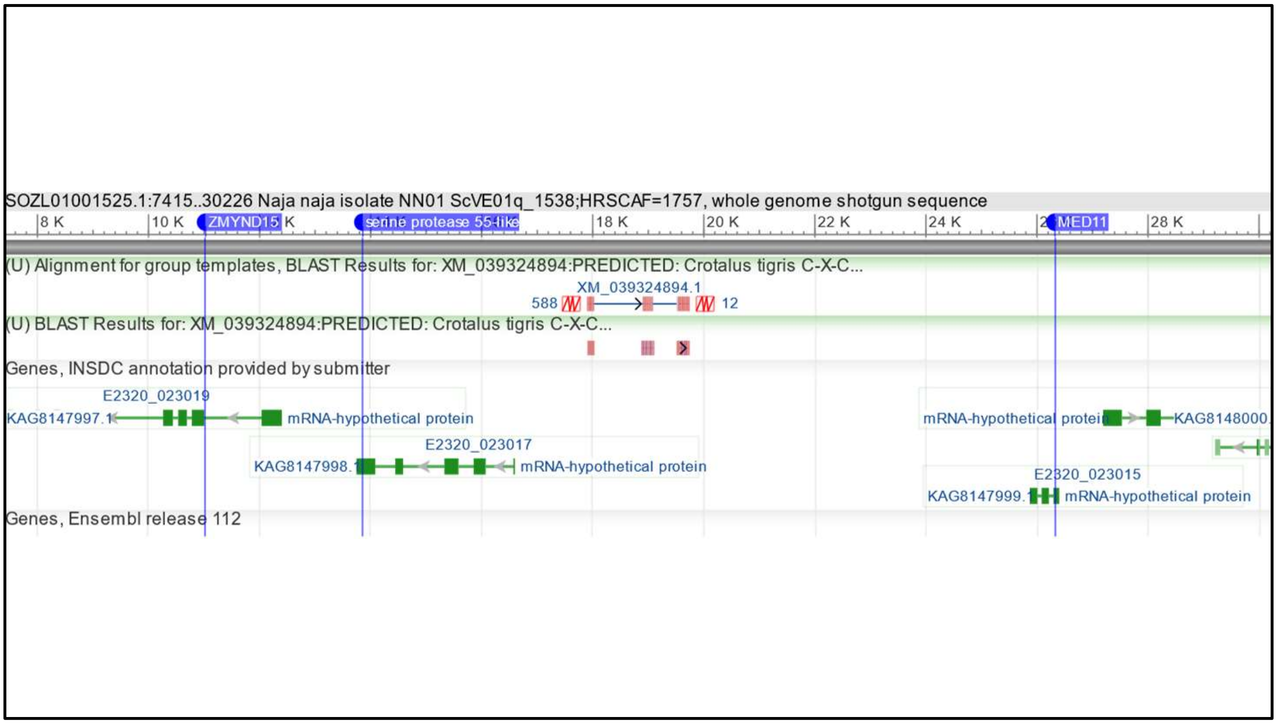


**Supplementary Figure 4:** Conserved gene synteny at the *CXCL16* locus in Indian cobra (Naja naja). BLASTn hits for Crotalus tigris *CXCL16* (XM_039324894.1) are shown in red on the SOZL01001525.1 scaffold of the Indian cobra genome. The *ZMYND15* and *PRSS55* genes are located on the left flank of *CXCL16*, while *MED11* is present on the right flank.

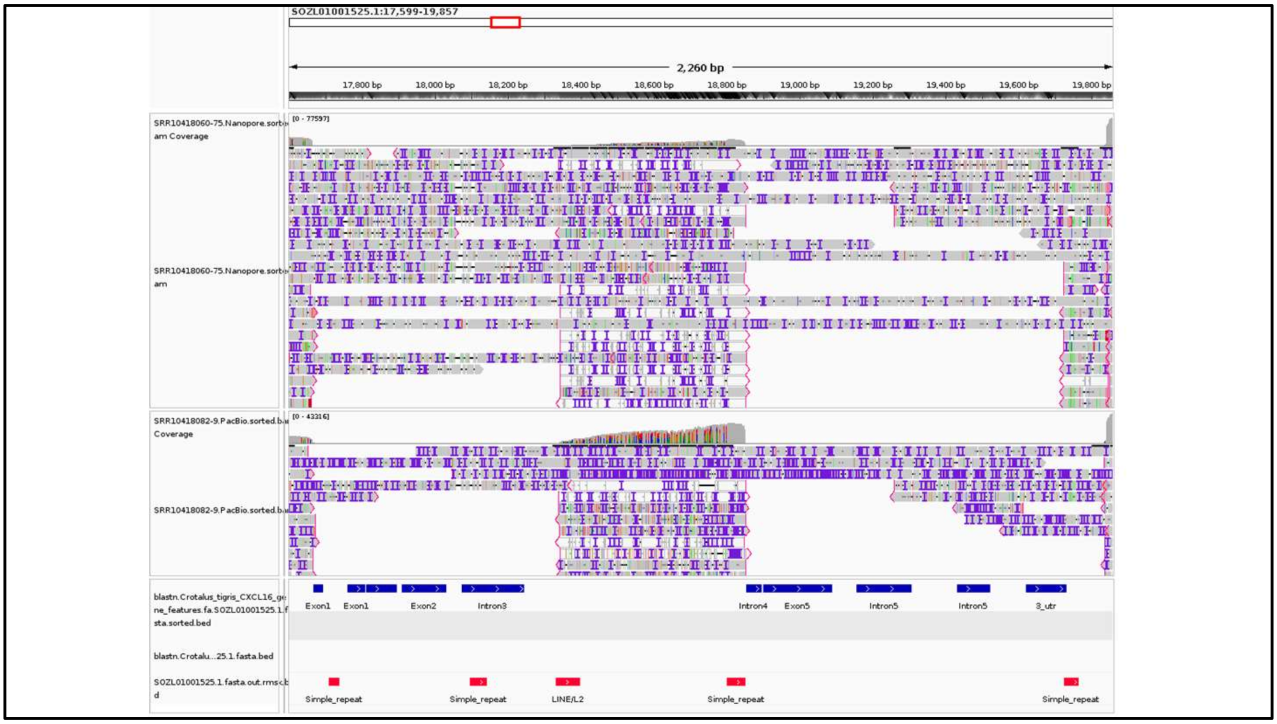


**Supplementary Figure 5:** Assembly verification at the *CXCL16* syntenic locus in Indian cobra (Naja naja). The IGV screenshot displays Nanopore and PacBio reads spanning the entire *CXCL16* locus. Exons and introns of CXCL16 are highlighted in blue. The bottom red bed track indicates the locations of repetitive sequences. No BLASTn hits were identified for intron 2 and exon 3 of *CXCL16*.

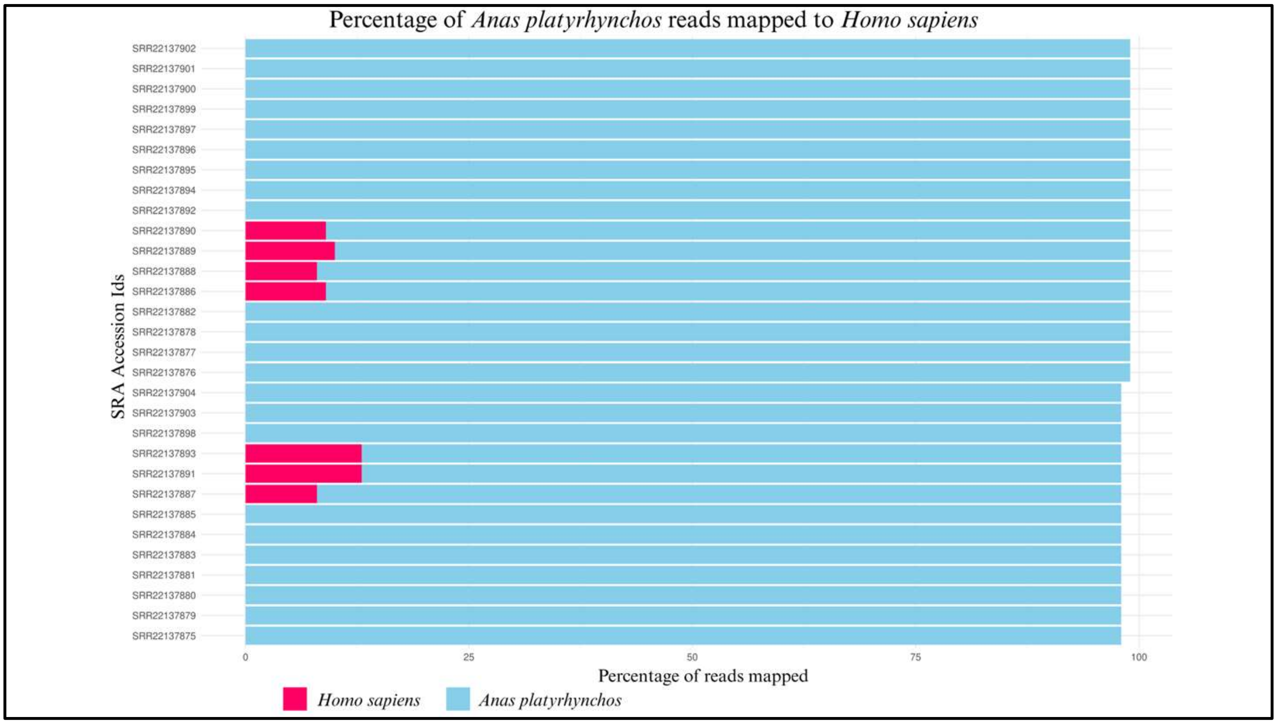


**Supplementary Figure 6:** Sample contamination in mallard SRA. The SRA study (NCBI accession no PRJNA896757) has an identical copy of human *CXCL16*. Read mapping to mallard and human genomes reveals contaminated samples. The Y-axis represents the SRA accession IDs, while the X-axis displays the percentage of reads mapped to human and mallard genomes, indicated by pink and sky-blue colours, respectively.

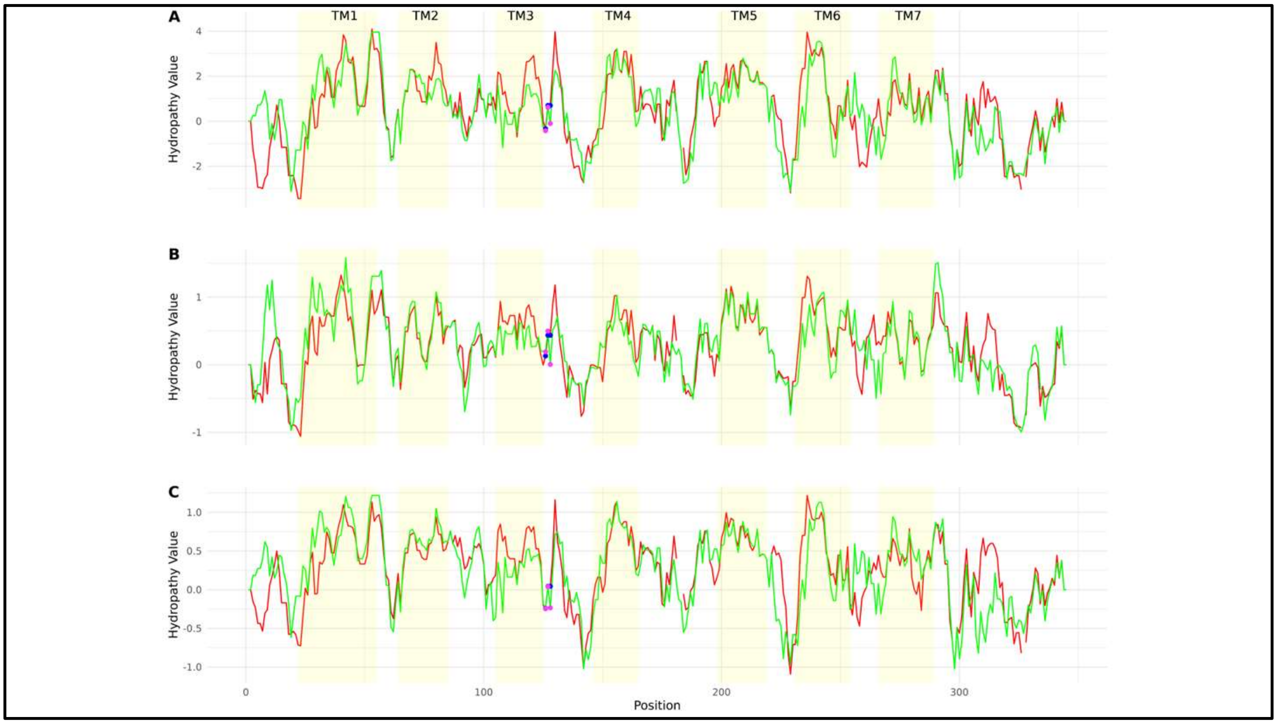


**Supplementary Figure 7:** Hydropathic properties of CXCR6 in humans and mallards. Hydropathy plot for humans (NP_006555.1) - red line and mallard (XP_005016704.3) - green line. Human and mallard hydropathy was obtained by giving amino acid sequences in EMBOSS Pepinfo with Kyte & Doolittle hydropathy parameters (**A**), OHM hydropathy parameters-Sweet & Eisenberg (**B**), Consensus parameters-Eisenberg et al.(**C**). The hydropathy value (y-axis) and residue position (x-axis) are plotted in R using ggplot2. DRF and DRL residues in humans and mallards are shown with blue and pink dots, respectively. The yellow portion represents the transmembrane.

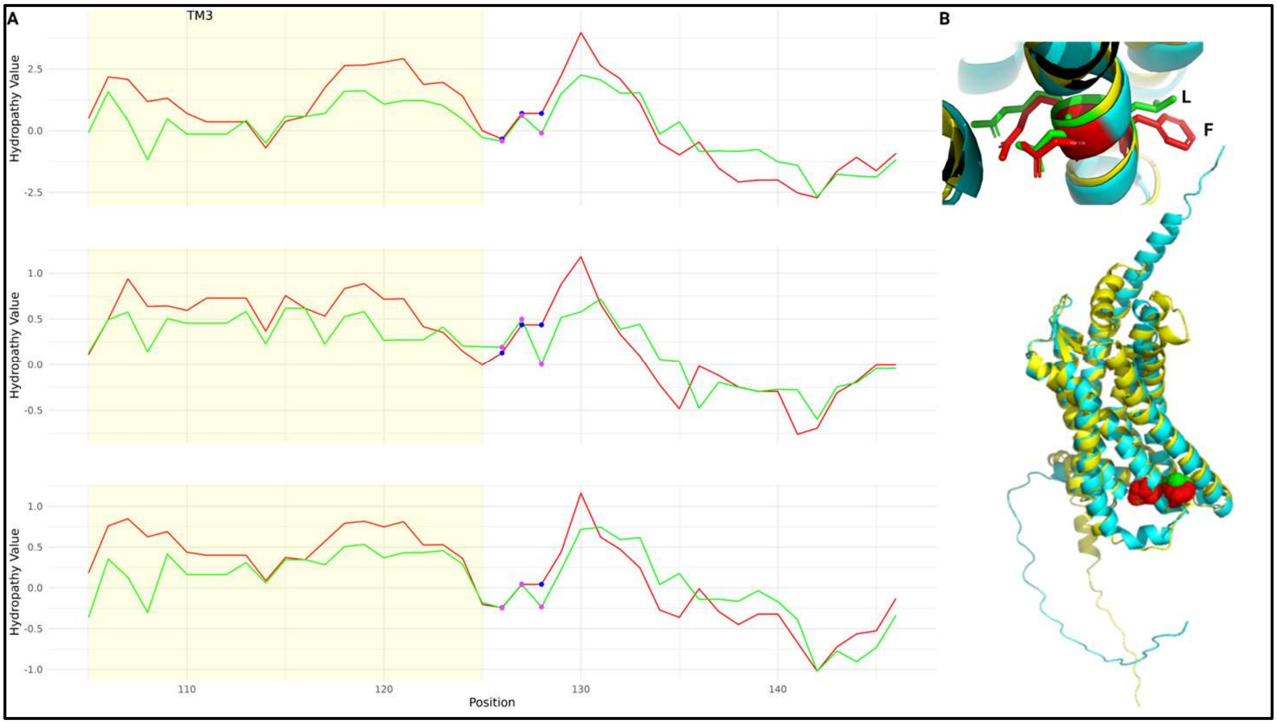


**Supplementary Figure 8:** Change in hydropathic properties in the DRF/DRL region. **A.** Hydropathy plot for humans (NP_006555.1) - red line and mallard (XP_005016704.3) - green line. Human and mallard hydropathy was obtained by giving amino acid sequences in EMBOSS Pepinfo with Kyte & Doolittle hydropathy parameters (top panel), OHM hydropathy parameters-Sweet & Eisenberg (middle panel), Consensus parameters-Eisenberg et al.(bottom panel). DRF and DRL residues in humans and mallards are shown with blue and pink dots, respectively. The hydropathy value (y-axis) and residue position (x-axis) are plotted in R using ggplot2. The yellow portion represents transmembrane three. **B.** 3D-structure comparison of human and mallard. The top panel shows DRF (red colour amino acids) and DRL (green colour amino acids) motif. The bottom panel shows the alignment of the 3D structure between human (cyan) and mallard (yellow).

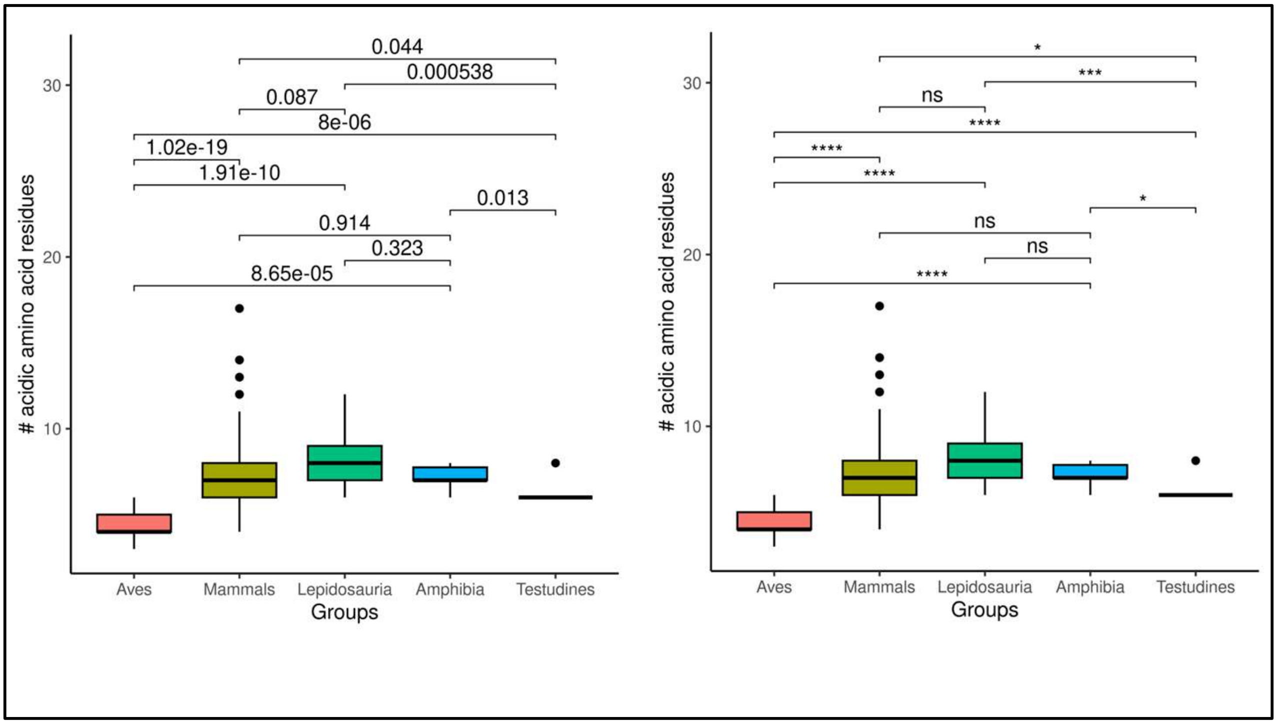


**Supplementary Figure 9:** Pairwise comparison of the number of acidic amino acid residues in the N-terminal. The x-axis represents various groups such as Amphibia, Aves, Mammals, Lepidosauria, and Testudines. The y-axis represents the number of acidic residues. Medians are compared with the Wilcoxon test using ggplot2 in R. The left-side boxplot shows the p-Values (above the inverted brackets for the groups), whereas the right-side boxplot shows the significance.

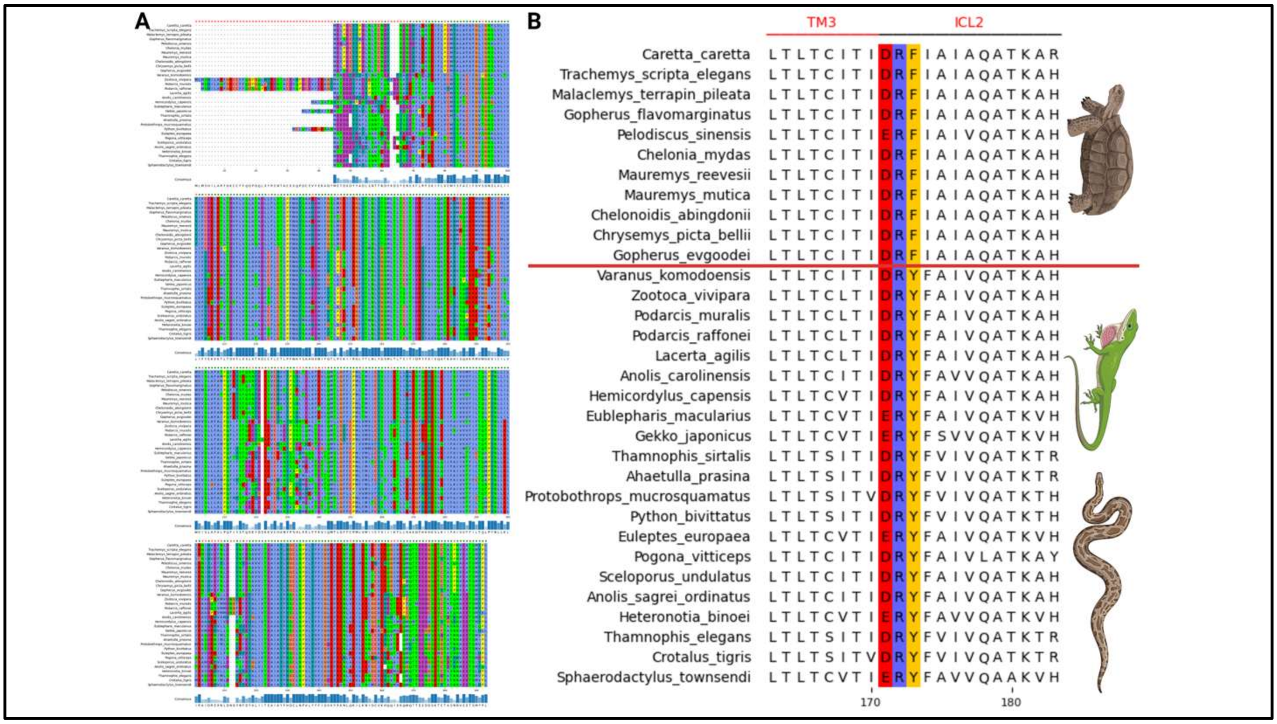


**Supplementary Figure 10:** CXCR6 gene conservation and comparison of DRF/DRY motif in lizards, snakes, and turtles. **A.** The MSA of lizards, snakes, and turtles. The red crosses on the alignment indicate conservation of less than 50%. The bottom blue bars represent the Consensus of the sequence. The amino acids coloured are based on the Clustal format. **B.** comparison of H3C region of reptiles. The red horizontal line separates turtles from other reptiles. The DRF/DRY motif amino acid residues are highlighted in Clustal format.

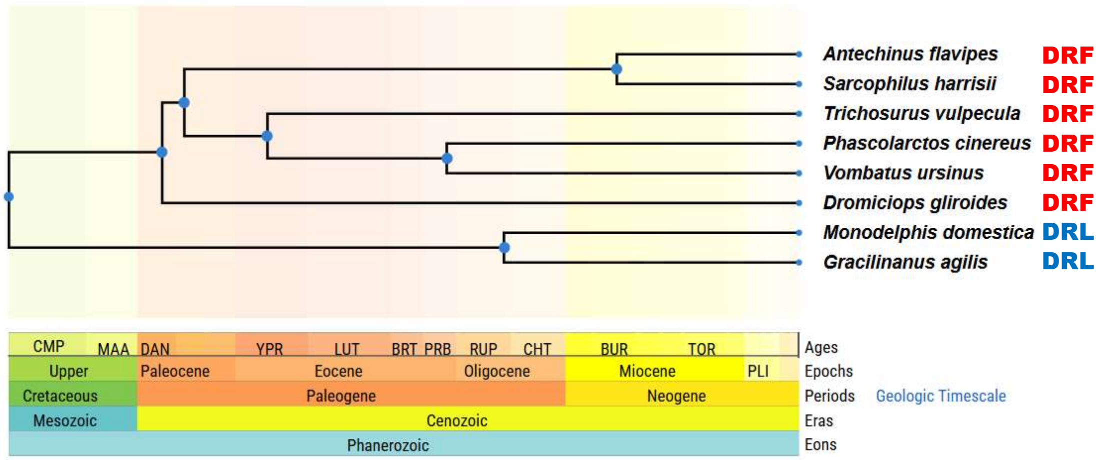


**Supplementary Figure 11:** Variation in DRF/DRL motif of marsupials. The Didelphimorphia species (Agile gracile opossum (*Gracilinanus agilis*) and Gray short-tailed opossum (*Monodelphis domestica*)) have the aves-like DRL motif, while other species have mammals-like DRF motif.

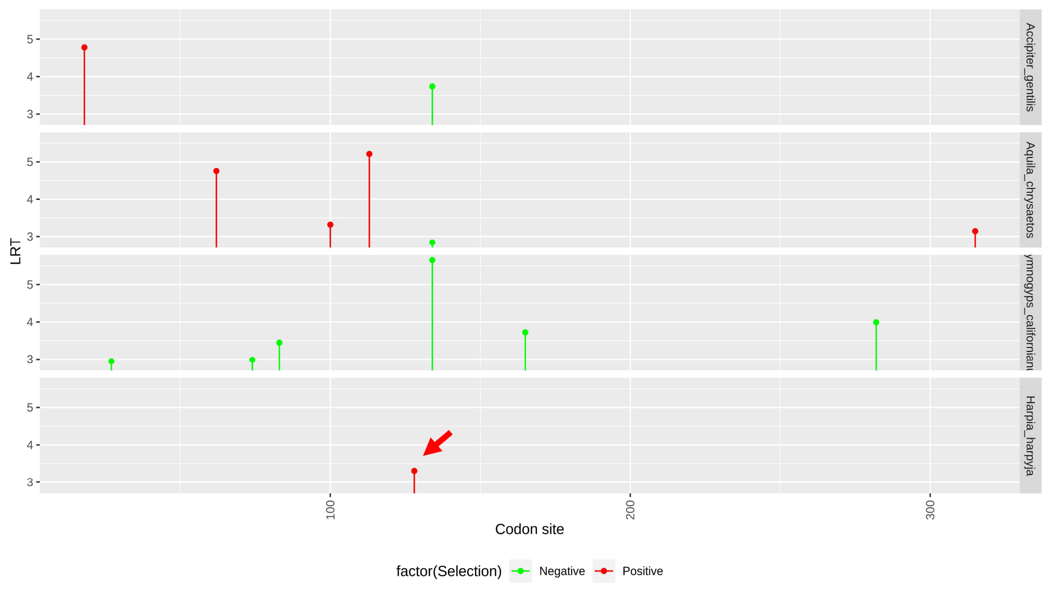


**Supplementary Figure 12:** Positive selection in the harpy eagle (*Harpia harpyja*)'s DRF motif. Positive selection at the DRF motif's third position (red arrow) was estimated using HYPHY's Fixed Effects Likelihood (FEL) model. The Y-axis represents Likelihood Ratio Test (LRT) values, and codon sites are on the X-axis. The lollipop plot was generated using ggplot in R programming.

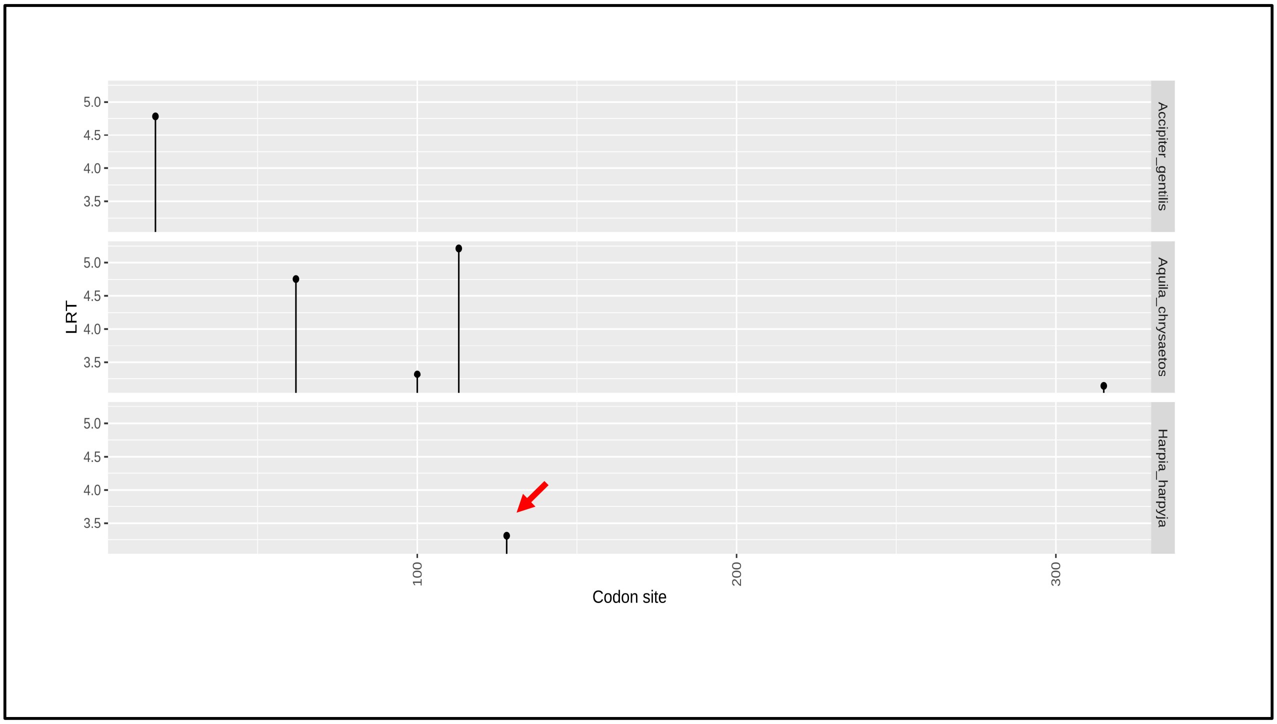


**Supplementary Figure 13:** Positive selection in the harpy eagle (Harpia harpyja)'s DRF motif. Positive selection at the DRF motif's third position (red arrow) was estimated using the MEME (Mixed Effects Model of Evolution) model of HYPHY. MEME is used to find sites that have experienced episodic diversification. The Y-axis represents Likelihood Ratio Test (LRT) values, and codon sites are on the X-axis. The lollipop plot was generated using ggplot in R programming.

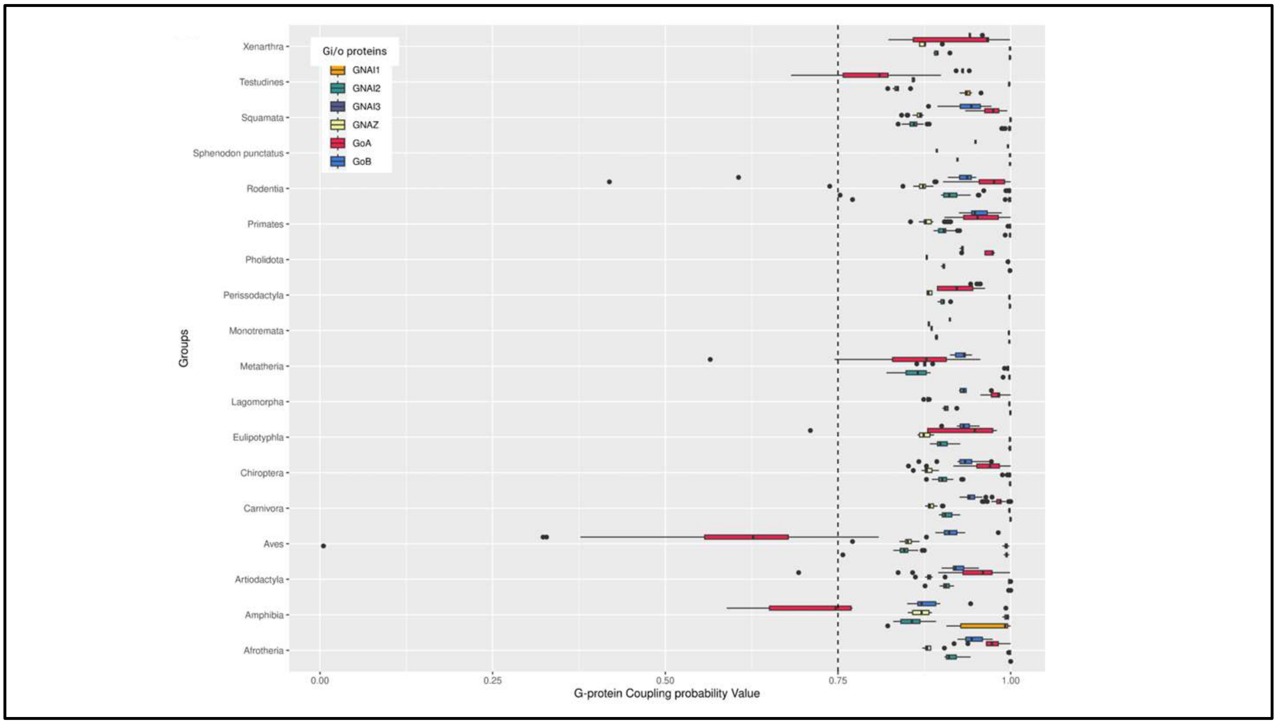


**Supplementary Figure 14:** PRECOGX predicted G-protein coupling probabilities for CXCR6 orthologs across vertebrates. The X-axis displays the coupling probabilities for Gi/o G-protein, while the Y-axis represents diverse vertebrate lineages. The vertical dotted line indicates probabilities above 75%. PRECOGx was used to predict coupling probabilities, and a boxplot plot was generated using ggplot in R programming.

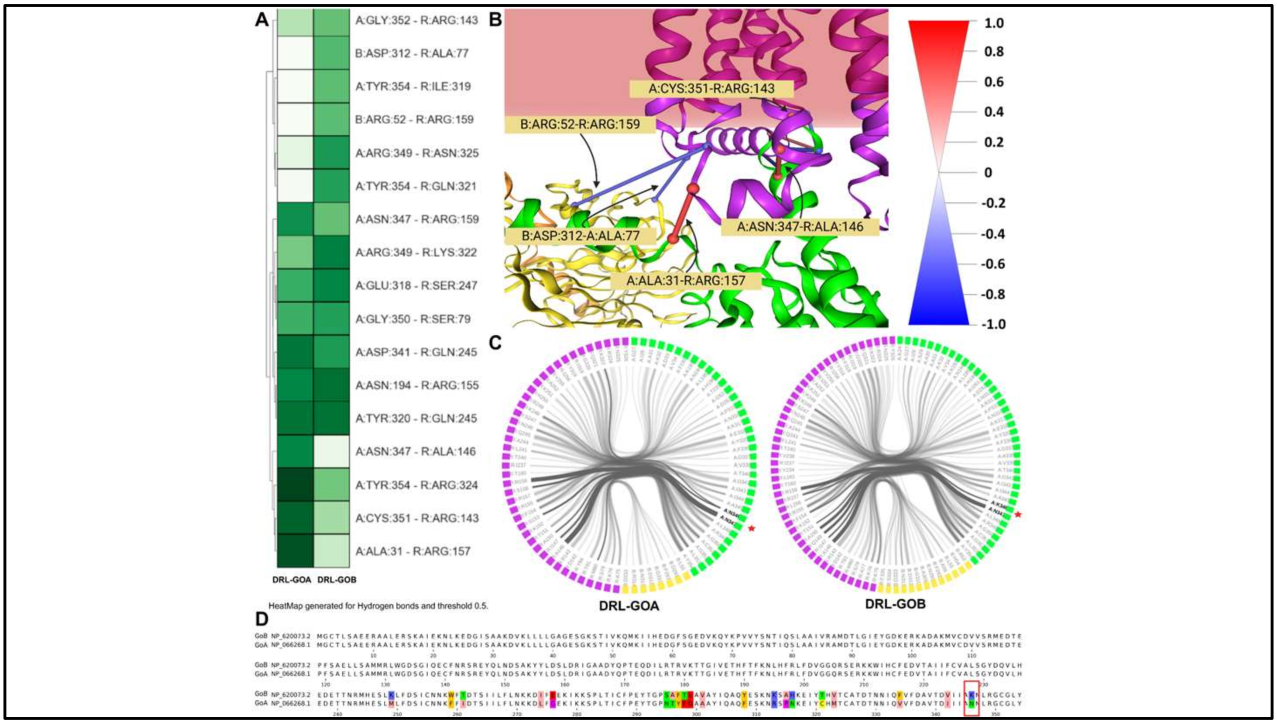


**Supplementary Figure 15: A.** The heatmap for hydrogen bonds compares MD simulations for DRL-GOA and DRL-GOB-like proteins, showing changes in residue interaction frequency due to the substitution of N with K in the GOA protein. **B.** Important sites implicated in the difference in interaction frequency are visualized in the 3D structure. **C.** The flareplot represents the interaction of N or K (highlighted by a red star) with the H3C region of CCR6 (DRL), with black connecting lines indicating the interaction between the receptor (pink) and GoA/GoB-like protein (green). **D.** Multiple sequence alignment between human GoA and GoB proteins reveals the N to K change at position 346, highlighted in the red box. The frequency difference cutoff was set to 0.5 to show meaningful interaction.

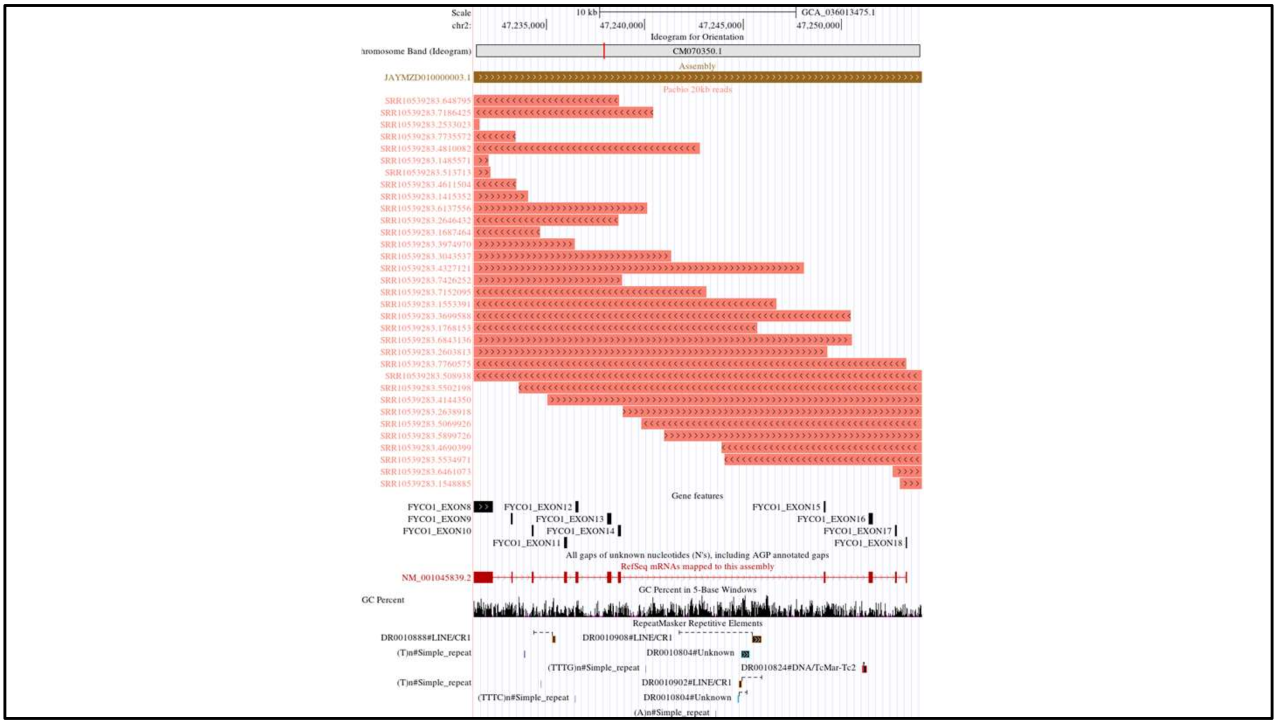


**Supplementary Figure 16:** Verification of the *Columba livia* (pigeon) genome assembly using long reads at the *CXCR6* gene locus. The UCSC genome browser image showing the alignment of ≥20 kb long reads generated using PacBio (SRR10539283) sequencing (salmon coloured), aligned to the pigeon (GCA_036013475.1) at the syntenic location of the *CXCR6*. The Gene feature track shows the *FYCO1* gene exons. The RepeatMasker BED track shows the repeats in that region. The accession IDs of the reads are specified on the left side of the reads beside them. Overlapping reads span the entire region at the syntenic location of the *CXCR6* gene, including the flanking exons of the *FYCO1* gene (Exon-14 and Exon-15).

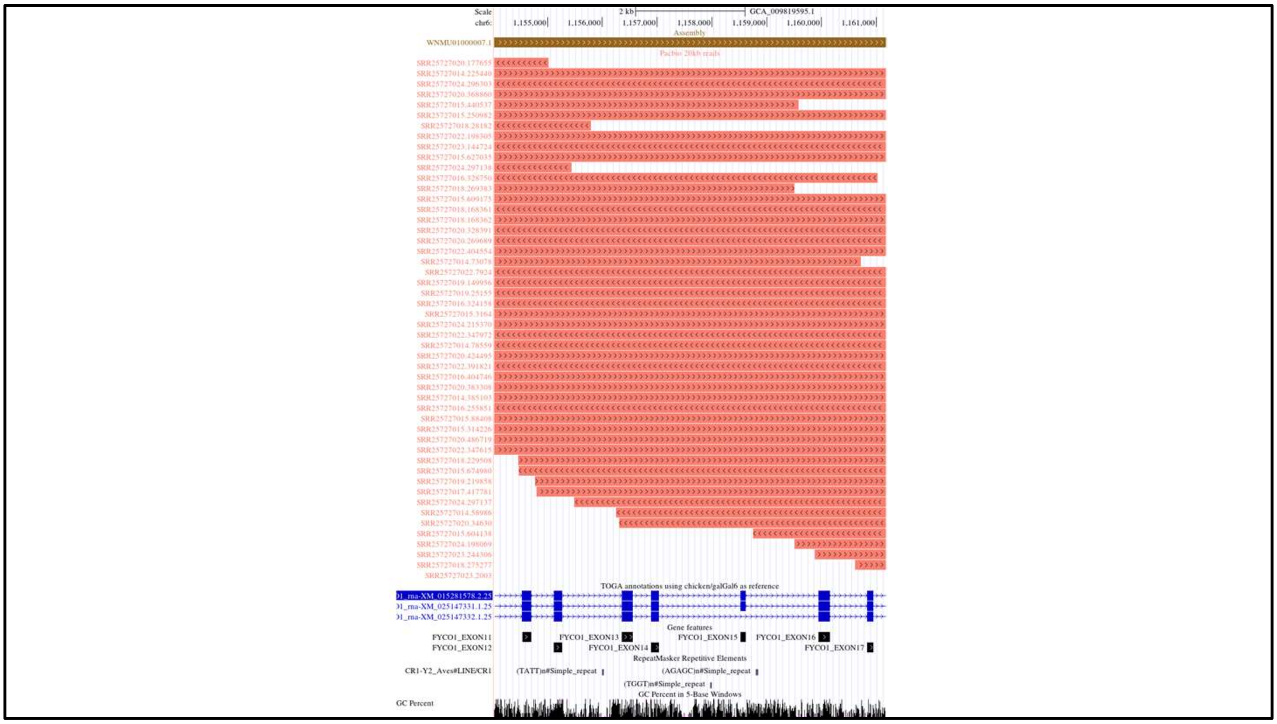


**Supplementary Figure 17:** Verification of the *Merops nubicus* (Northern carmine bee-eater)genome assembly using long reads at the *CXCR6* gene locus. The UCSC genome browser image showing the alignment of ≥20 kb long reads generated using PacBio (SRR25727014, SRR25727015, SRR25727016, SRR25727017, SRR25727018, SRR25727019, SRR25727020, SRR25727022, SRR25727023, and SRR25727024) sequencing (salmon coloured), aligned to the Northern carmine bee-eater (GCA_009819595.1) at the syntenic location of the *CXCR6*. The Gene feature track shows the *FYCO1* gene exons. The RepeatMasker BED track shows the repeats in that region. The accession IDs of the reads are specified on the left side of the reads beside them. Overlapping reads span the entire region at the syntenic location of the *CXCR6* gene, including the flanking exons of the *FYCO1* gene (Exon-14 and Exon-15).

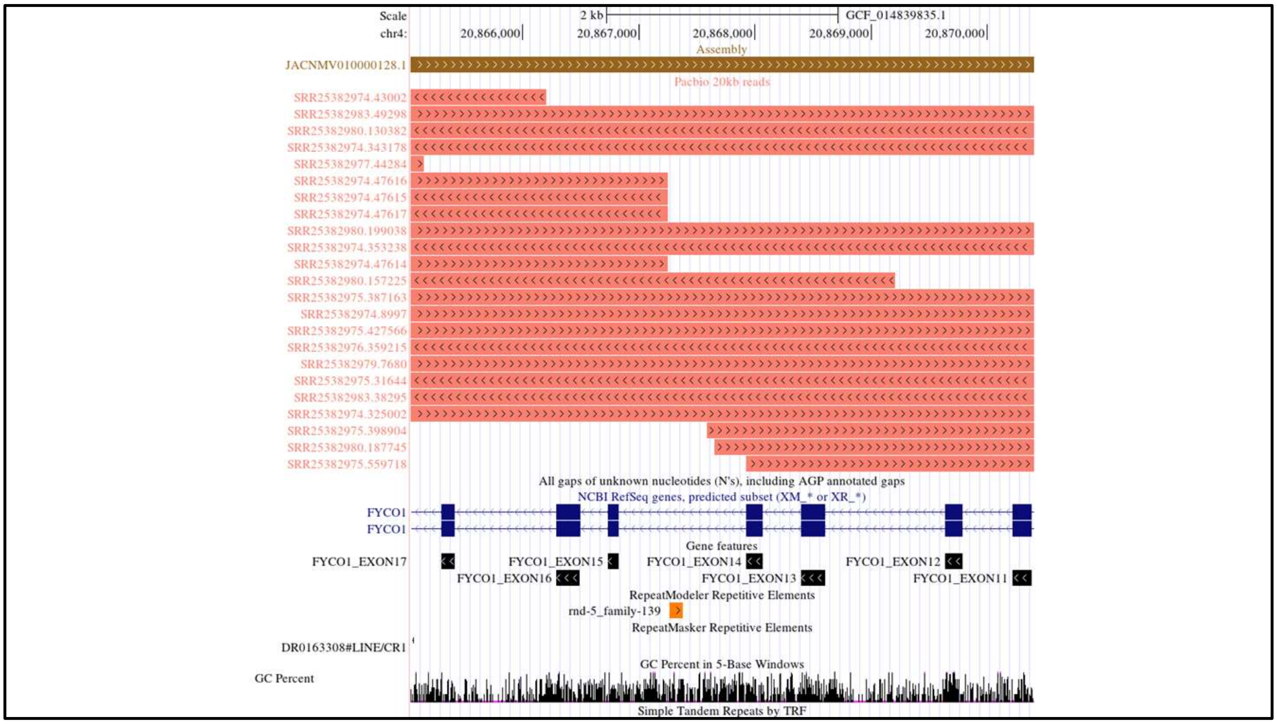


**Supplementary Figure 18:** Verification of the *Dryobates pubescens* (downy woodpecker) genome assembly using long reads at the *CXCR6* gene locus. The UCSC genome browser image showing the alignment of ≥20 kb long reads generated using PacBio (SRR25382974, SRR25382975, SRR25382976, SRR25382977, SRR25382979, SRR25382980, SRR25382982, and SRR25382983) sequencing (salmon coloured), aligned to the downy woodpecker (GCF_014839835.1) at the syntenic location of the *CXCR6*. The Gene feature track shows the *FYCO1* gene exons. The RepeatMasker BED track shows the repeats in that region. The accession IDs of the reads are specified on the left side of the reads beside them. Overlapping reads span the entire region at the syntenic location of the *CXCR6* gene, including the flanking exons of the *FYCO1* gene (Exon-14 and Exon-15).

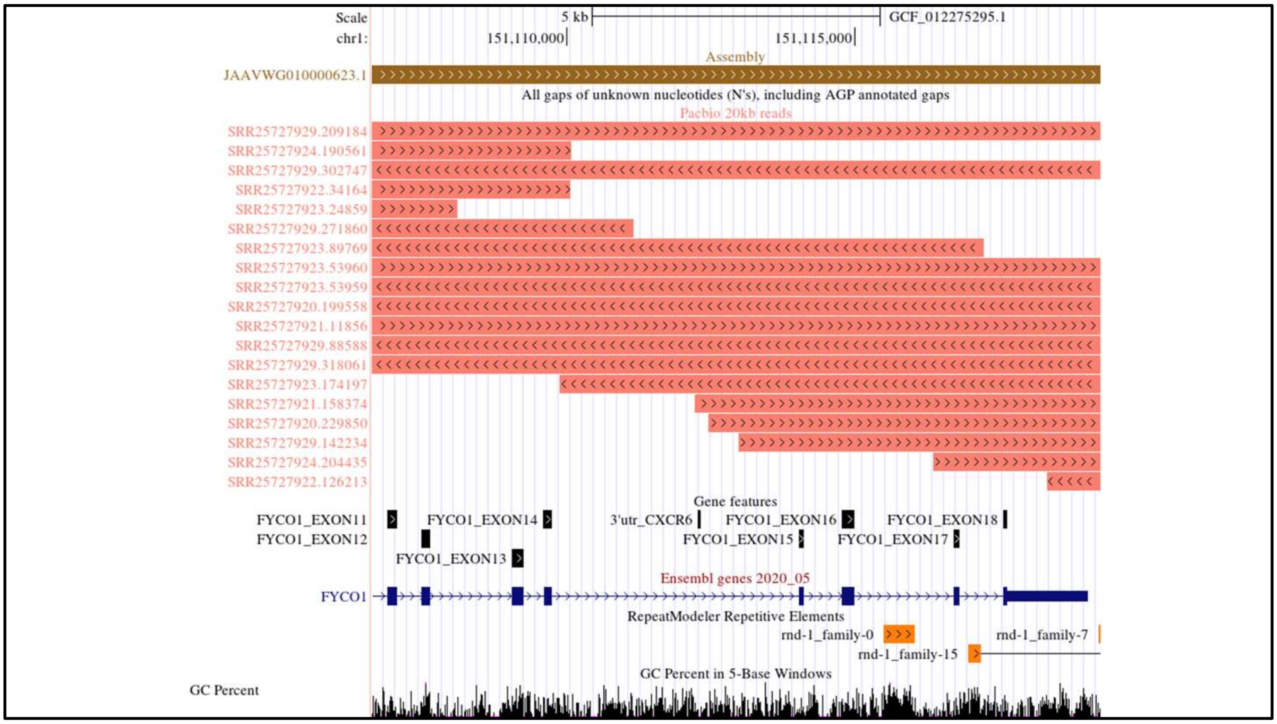


**Supplementary Figure 19:** Verification of the *Melopsittacus undulatus* (budgerigar) genome assembly using long reads at the *CXCR6* gene locus. The UCSC genome browser image showing the alignment of ≥20 kb long reads generated using PacBio (ERR244164, ERR244165, ERR244166, SRR25727920, SRR25727921, SRR25727922, SRR25727923, SRR25727924, SRR25727928, and SRR25727929) sequencing (salmon coloured), aligned to the budgerigar (GCF_012275295.1) at the syntenic location of the *CXCR6*. The Gene feature track shows the *FYCO1* gene exons. The RepeatMasker BED track shows the repeats in that region. The accession IDs of the reads are specified on the left side of the reads beside them. Overlapping reads span the entire region at the syntenic location of the *CXCR6* gene, including the flanking exons of the *FYCO1* gene (Exon-14 and Exon-15).

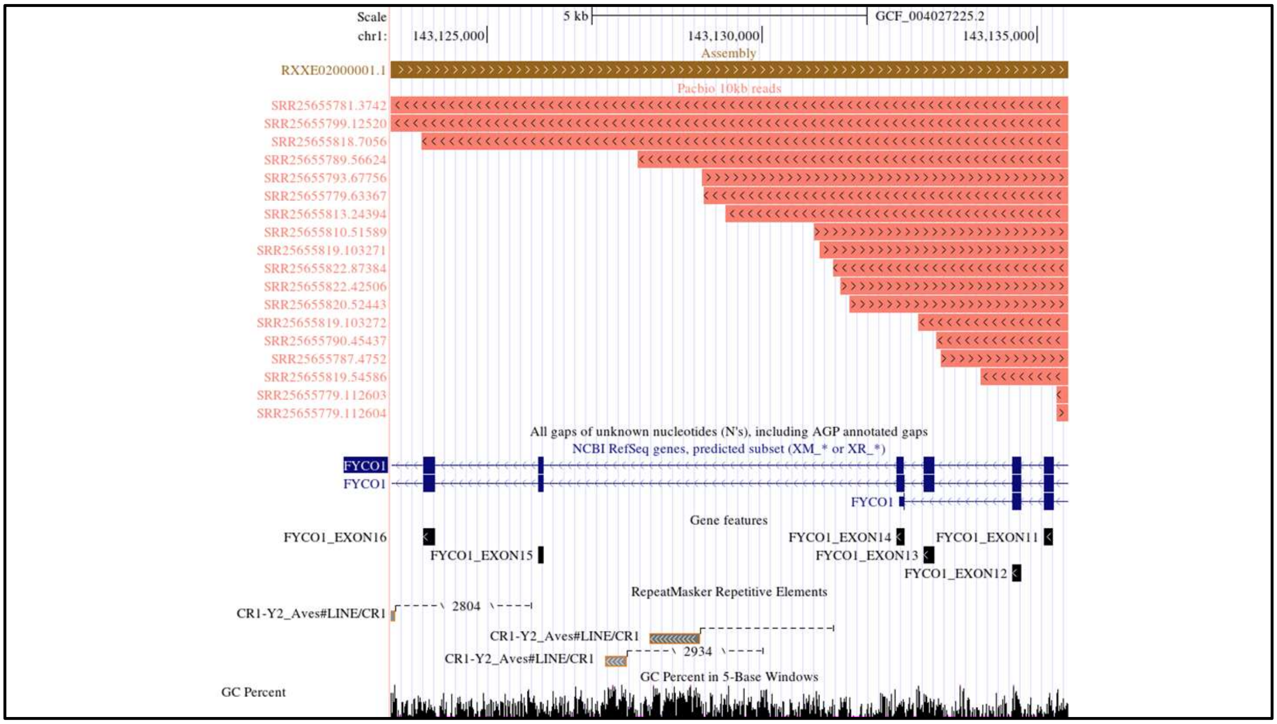


**Supplementary Figure 20:** Verification of the *Strigops habroptila* (kākāpō) genome assembly using long reads at the *CXCR6* gene locus. The UCSC genome browser image showing the alignment of ≥20 kb long reads generated using PacBio (SRR25655775 to SRR25655825) sequencing (salmon coloured), aligned to the kākāpō (GCF_004027225.2) at the syntenic location of the *CXCR6*. The Gene feature track shows the *FYCO1* gene exons. The RepeatMasker BED track shows the repeats in that region. The accession IDs of the reads are specified on the left side of the reads beside them. Overlapping reads span the entire region at the syntenic location of the *CXCR6* gene, including the flanking exons of the *FYCO1* gene (Exon-14 and Exon-15).

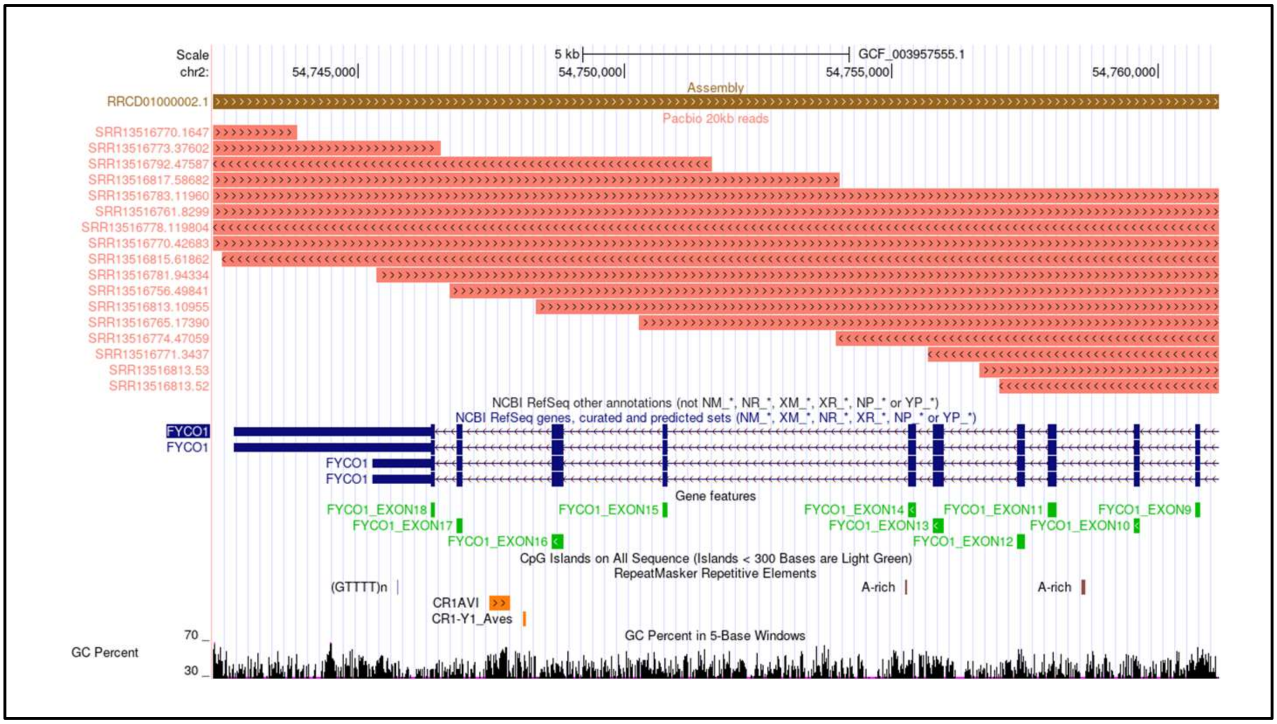


**Supplementary Figure 21:** Verification of the *Calypte anna* (Anna's hummingbird) genome assembly using long reads at the *CXCR6* gene locus. The UCSC genome browser image showing the alignment of ≥20 kb long reads generated using PacBio (SRR13516755 to SRR13516817) sequencing (salmon coloured), aligned to Anna's hummingbird (GCF_003957555.1) at the syntenic location of the *CXCR6*. The Gene feature track shows the *FYCO1* gene exons. The RepeatMasker BED track shows the repeats in that region. The accession IDs of the reads are specified on the left side of the reads beside them. Overlapping reads span the entire region at the syntenic location of the *CXCR6* gene, including the flanking exons of the *FYCO1* gene (Exon-14 and Exon-15).

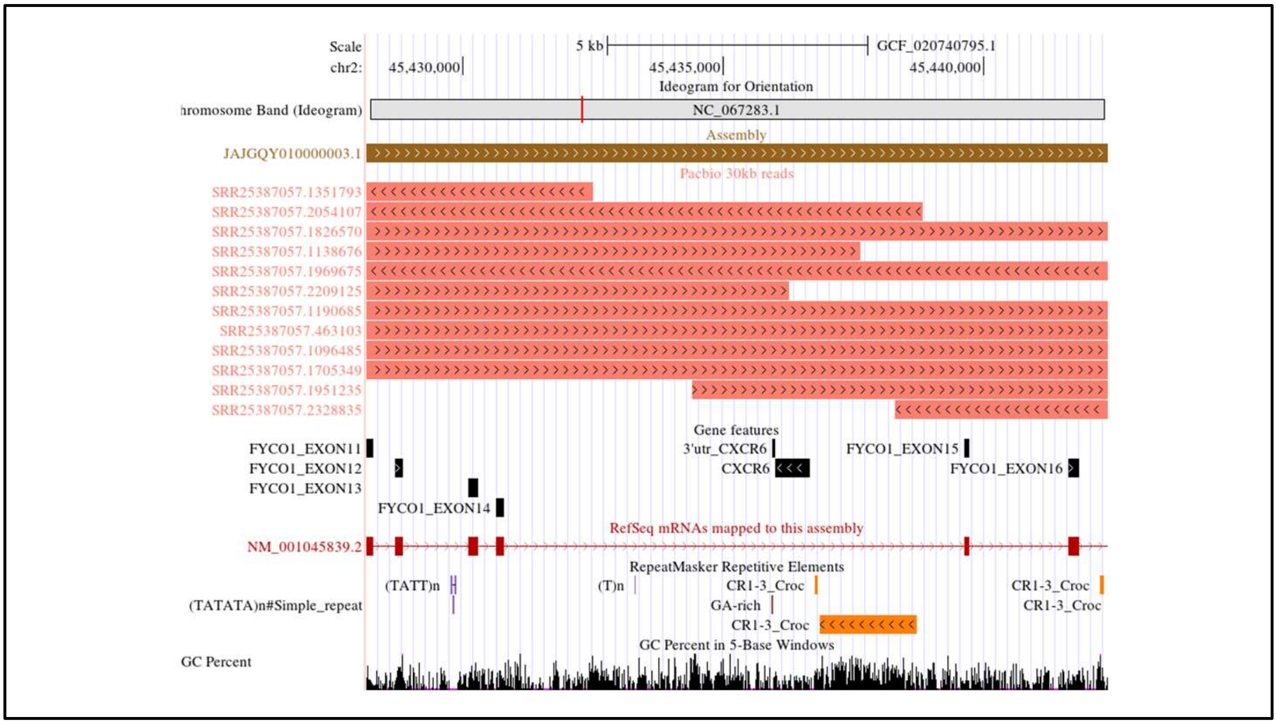


**Supplementary Figure 22:** Verification of the *Apus apus* (common swift) genome assembly using long reads at the *CXCR6* gene locus. The UCSC genome browser image showing the alignment of ≥30 kb long reads generated using PacBio (SRR25387057) sequencing (salmon coloured), aligned to the common swift (GCF_020740795.1) at the syntenic location of the *CXCR6*. The Gene feature track shows the *FYCO1* gene exons and *CXCR6* gene relics. The RepeatMasker BED track shows the repeats in that region. The accession IDs of the reads are specified on the left side of the reads beside them. Overlapping reads span the entire region at the syntenic location of the *CXCR6* gene, including the flanking exons of the *FYCO1* gene (Exon-14 and Exon-15).

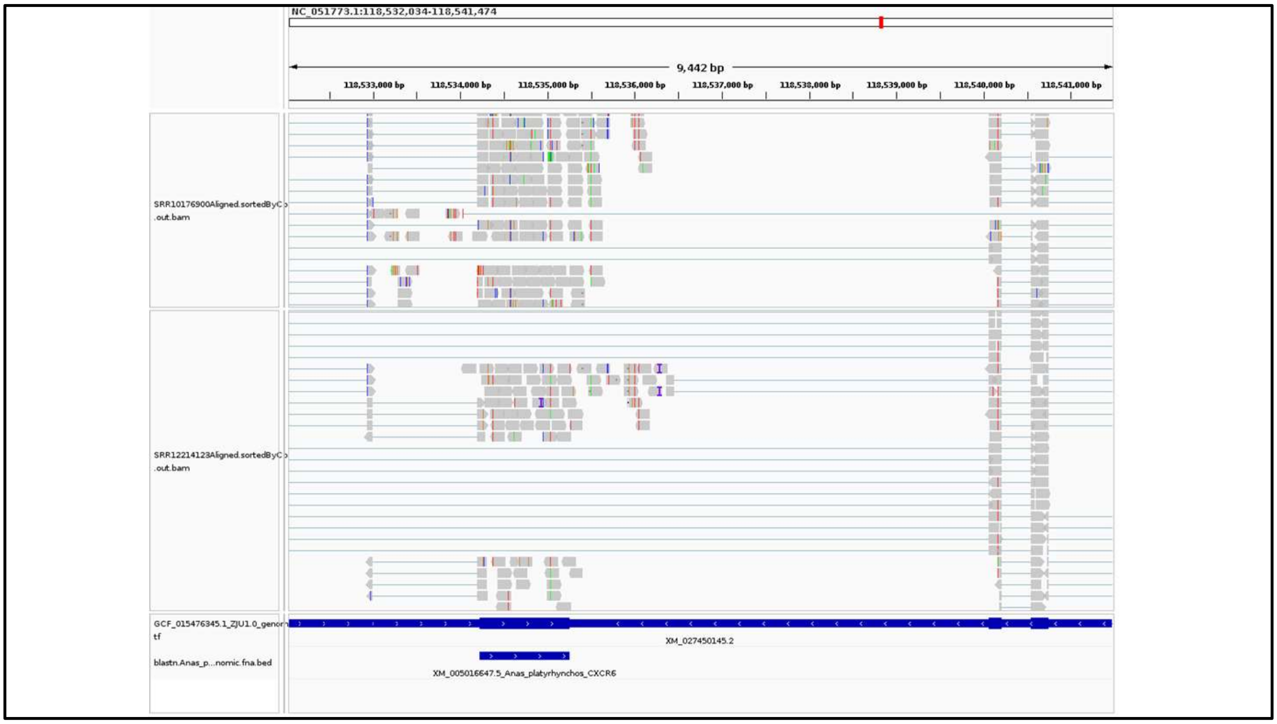


**Supplementary Figure 23:** The *CXCR6* gene is expressed in the spleen and liver of the mallard (*Anas platyrhynchos*). IGV screenshot for *CXCR6* expression for spleen (SRR10176900) and liver (SRR12214123) mapped to GCF_015476345.1 genome assembly of mallard (*Anas platyrhynchos*), using STAR mapper.

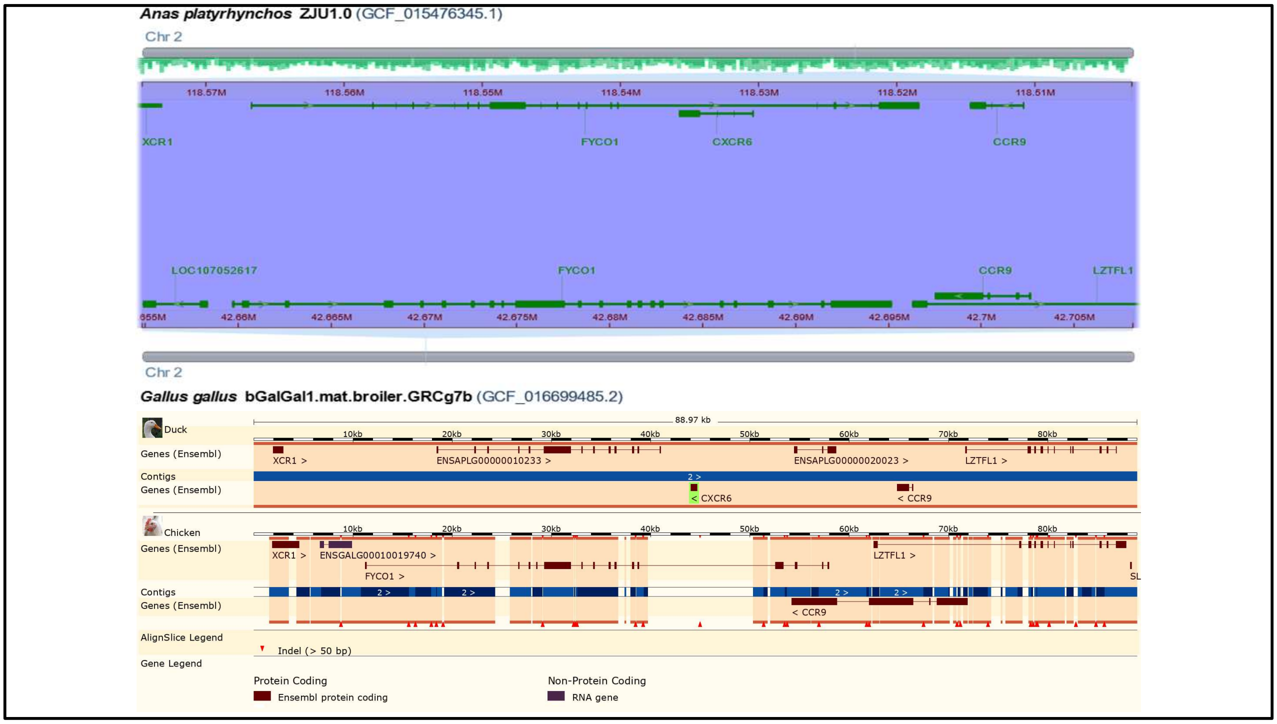


**Supplementary Figure 24:** The pairwise alignment between mallard and chicken shows the *CXCR6* is not present in chicken (top-panel: Image, taken from NCBI Comparative Genome Viewer) and the corresponding region is missing from chicken (bottom-panel: Image from ENSEMBL pairwise alignment).

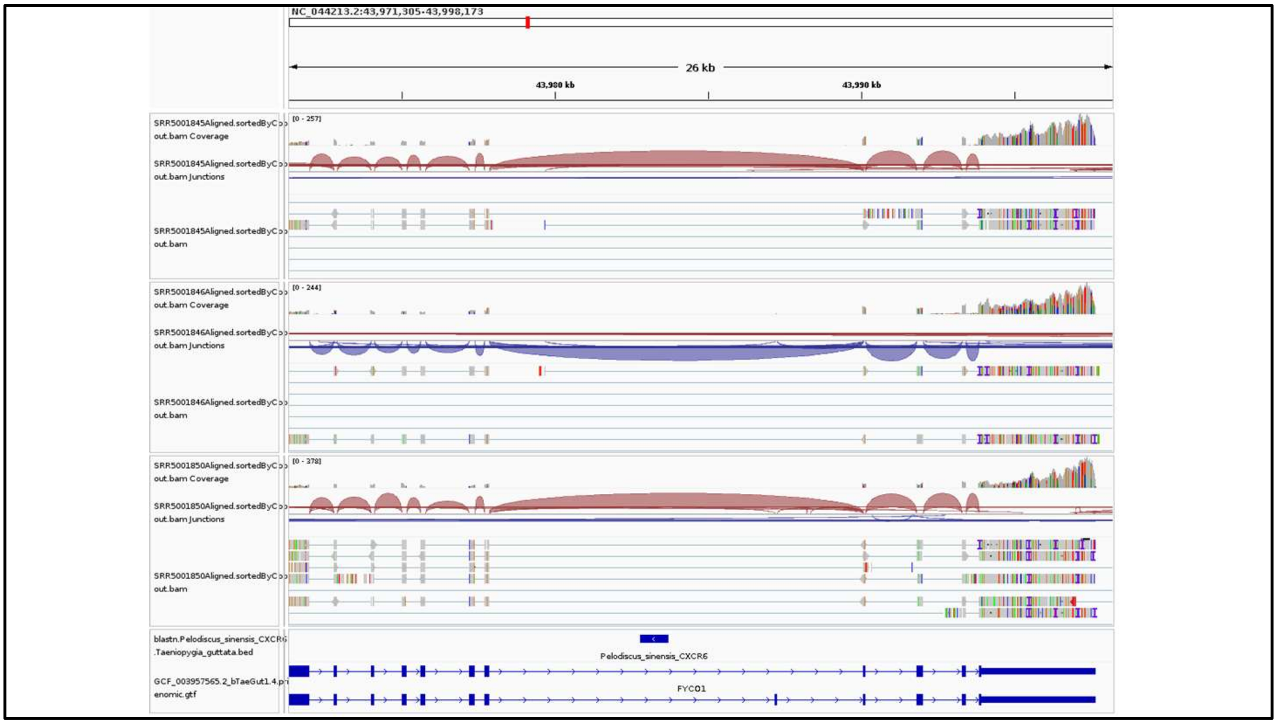


**Supplementary Figure 25:** The *CXCR6* gene lacks expression in the spleen of zebra finch (*Taeniopygia guttata*). IGV screenshot for *CXCR6* expression for spleen (SRR5001845, SRR5001846, SRR5001850) mapped to GCF_003957565.2 genome assembly of the zebra finch (*Taeniopygia guttata*), using STAR mapper.

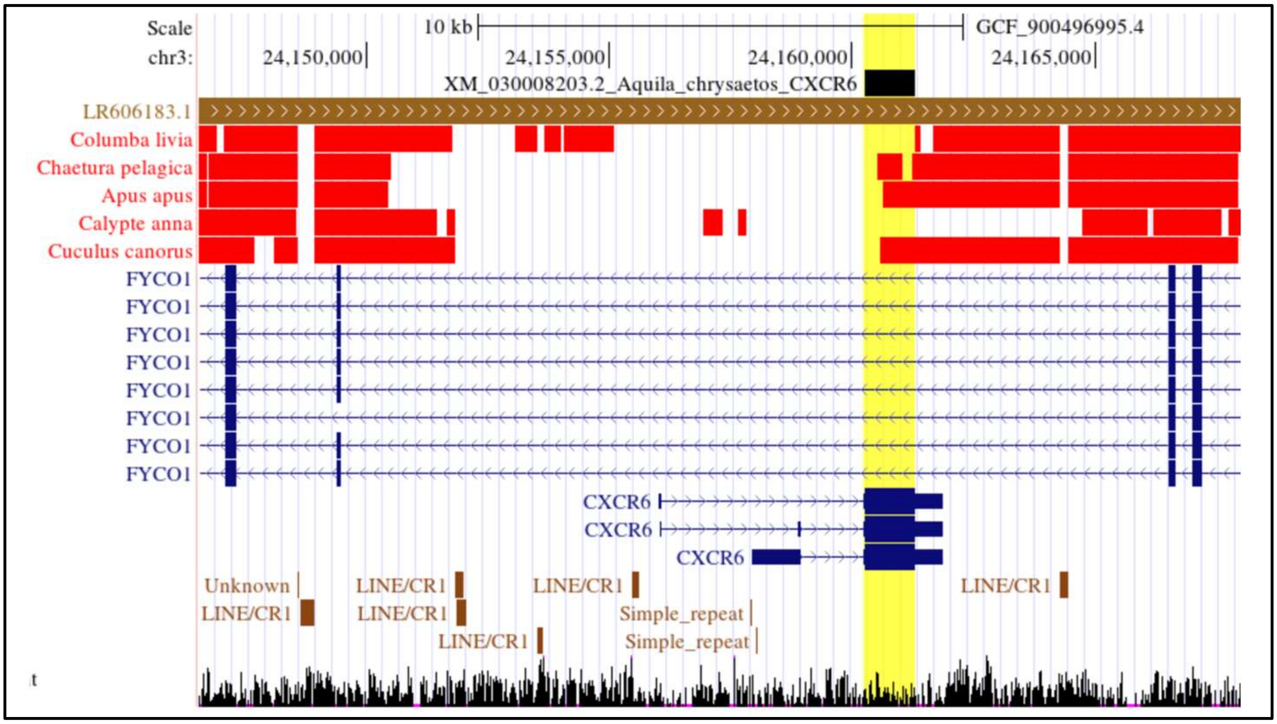


**Supplementary Figure 26:** Independent gene loss in birds. Deletion size comparison of *CXCR6* gene-containing segments with the species compared to the golden eagle (*Aquila chrysaetos*). The species with red colour have *CXCR6* gene deletion. The yellow highlight shows the *CXCR6* gene region. For deletion size estimation, the BLASTn of the shown species was performed with golden eagle repeat masked chromosome and corresponding blast hits of that species. The deletion boundaries are not the same, and their sizes also differ.

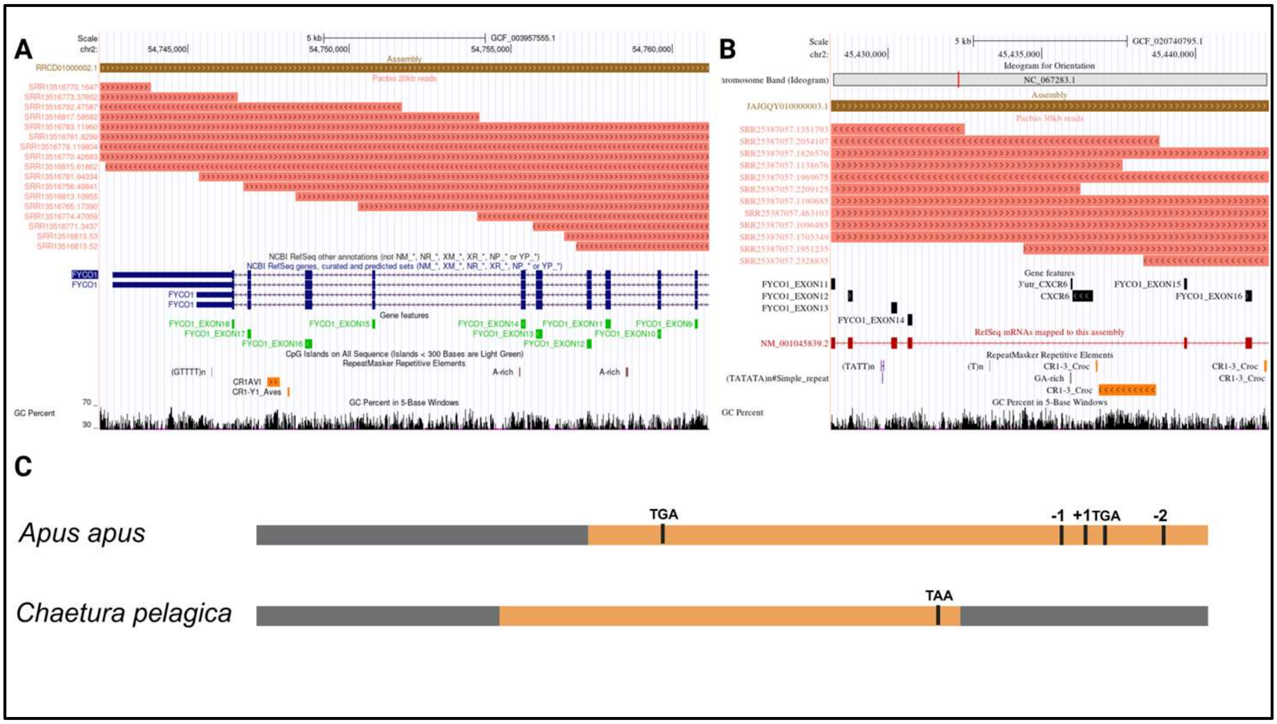


**Supplementary Figure 27**: Segmental deletion and *CXCR6* gene pseudogenization in Apodiformes. **A.** Verification of the *Calypte anna* (Anna's hummingbird) genome assembly using long reads at the *CXCR6* gene locus. The UCSC genome browser image shows the alignment of ≥20 kb long reads generated using PacBio sequencing (salmon coloured), aligned to Anna's hummingbird (GCF_003957555.1) at the syntenic location of the *CXCR6*. The Gene feature track shows the *FYCO1* gene exons. The RepeatMasker BED track shows the repeats in that region. The accession IDs of the reads are specified on the left side of the reads beside them. Overlapping reads span the entire region at the syntenic location of the *CXCR6* gene, including the flanking exons of the *FYCO1* gene (Exon-14 and Exon-15). **B.** Verification of the *Apus apus* (common swift) genome assembly using long reads at the *CXCR6* gene locus. The UCSC genome browser image showing the alignment of ≥30 kb long reads generated using PacBio (SRR25387057) sequencing (salmon coloured), aligned to the common swift (GCF_020740795.1) at the syntenic location of the *CXCR6*. The Gene feature track shows the *FYCO1* gene exons and *CXCR6* gene relics. The RepeatMasker BED track shows the repeats in that region. The accession IDs of the reads are specified on the left side of the reads beside them. Overlapping reads span the entire region at the syntenic location of the *CXCR6* gene, including the flanking exons of the *FYCO1* gene (Exon-14 and Exon-15). **C.** frame-disrupting changes in

*Apus apus* (common swift) and *Chaetura pelagica* (chimney swift). The frame-disrupting events in *Chaetura pelagica* (chimney swift) are only based on genome assembly.

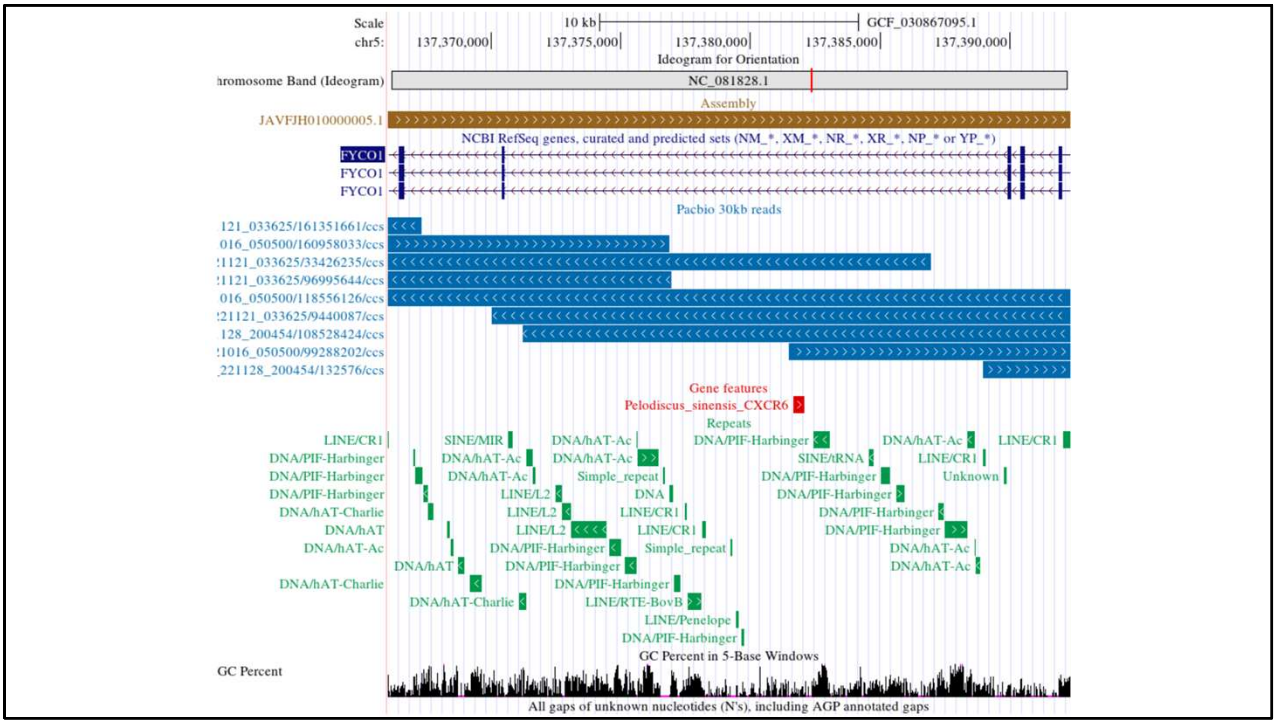


**Supplementary Figure 28:** Verification of the American alligator (*Alligator mississippiensis*) genome assembly using long reads at the *CXCR6* gene locus. The UCSC genome browser image showing the alignment of ≥30 kb long reads generated using PacBio sequencing (steel blue coloured), aligned to the *Alligator mississippiensis* (GCF_030867095.1) at the syntenic location of the *CXCR6* gene remnants. Gene features track shows partial remnants of the *Pelodiscus sinensis CXCR6* gene in *Alligator mississippiensis* genome assembly. Blue-colored boxes represent the *CXCR6* exon remnants. The Repeat track shows the repeats in that region (represented by dark green boxes). Overlapping reads span the entire region at the syntenic location of the *CXCR6* gene, including the flanking region (*FYCO1* exons).

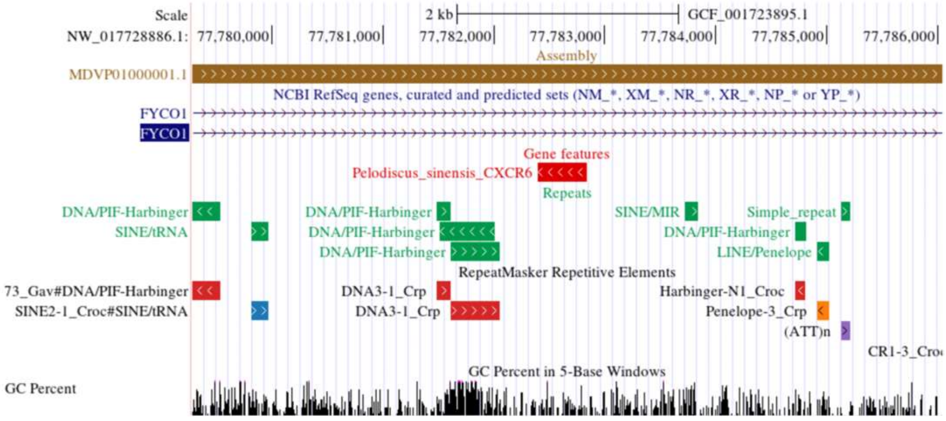


**Supplementary Figure 29:** Repeat identification in the saltwater crocodile (*Crocodylus porosus*) genome assembly (GCF_001723895.1) at the *CXCR6* gene locus using the UCSC genome browser. The Repeat track shows the repeats in that region (represented by dark green boxes). Gene features track shows partial remnants of the *Pelodiscus sinensis CXCR6* gene in *Crocodylus porosus* genome assembly.

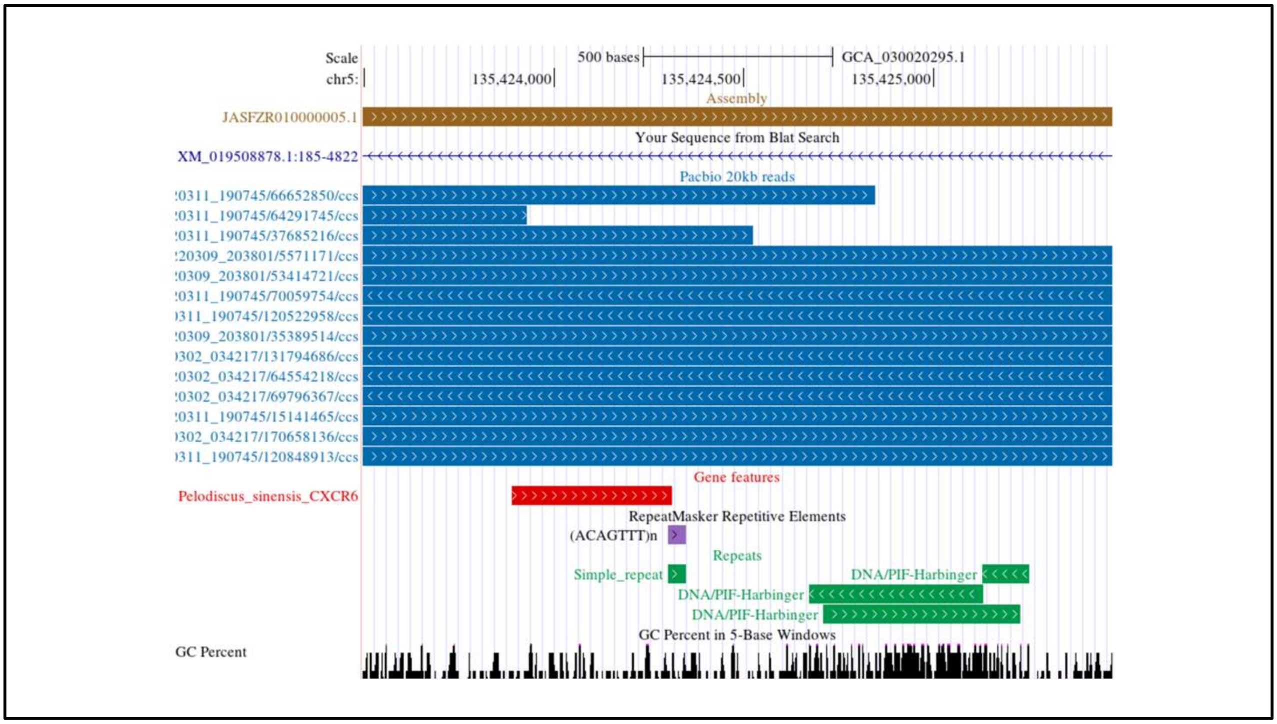


**Supplementary Figure 30:** Verification of the Gharial (*Gavialis gangeticus*) genome assembly using long reads at the *CXCR6* gene locus. The UCSC genome browser image showing the alignment of ≥20 kb long reads generated using PacBio sequencing (steel blue coloured), aligned to the *Gavialis gangeticus* (GCA_030020295.1) at the syntenic location of the *CXCR6* gene remnants. Gene features track shows partial remnants of the *Pelodiscus sinensis CXCR6* gene in *Gavialis gangeticus* genome assembly. Blue-colored boxes represent the *CXCR6* exon remnants. The Repeat track shows the repeats in that region (represented by dark green boxes). Overlapping reads span the entire region at the syntenic location of the *CXCR6* gene, including the flanking region.

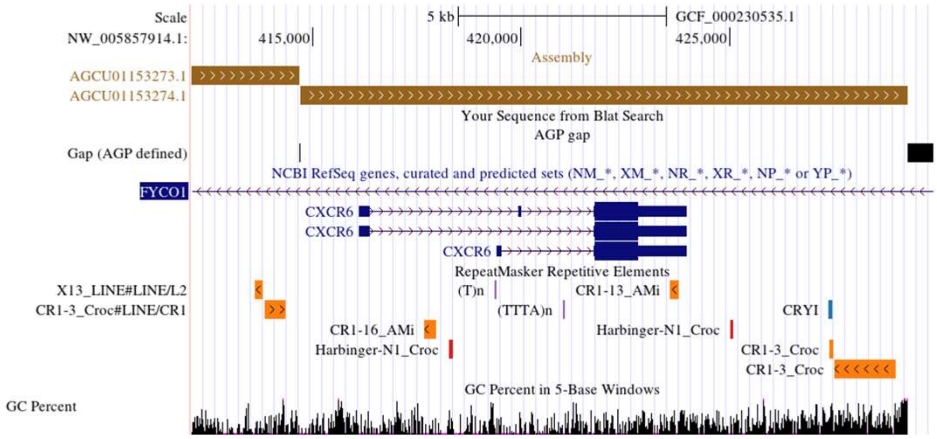


**Supplementary Figure 31:** Visualization of *CXCR6* gene locus in the Chinese softshell turtle (*Pelodiscus sinensis*) in the genome assembly (GCF_000230535.1) using the UCSC genome browser. RepeatMasker repetitive elements track shows repeat in that region. The bottom track represents GC percent.

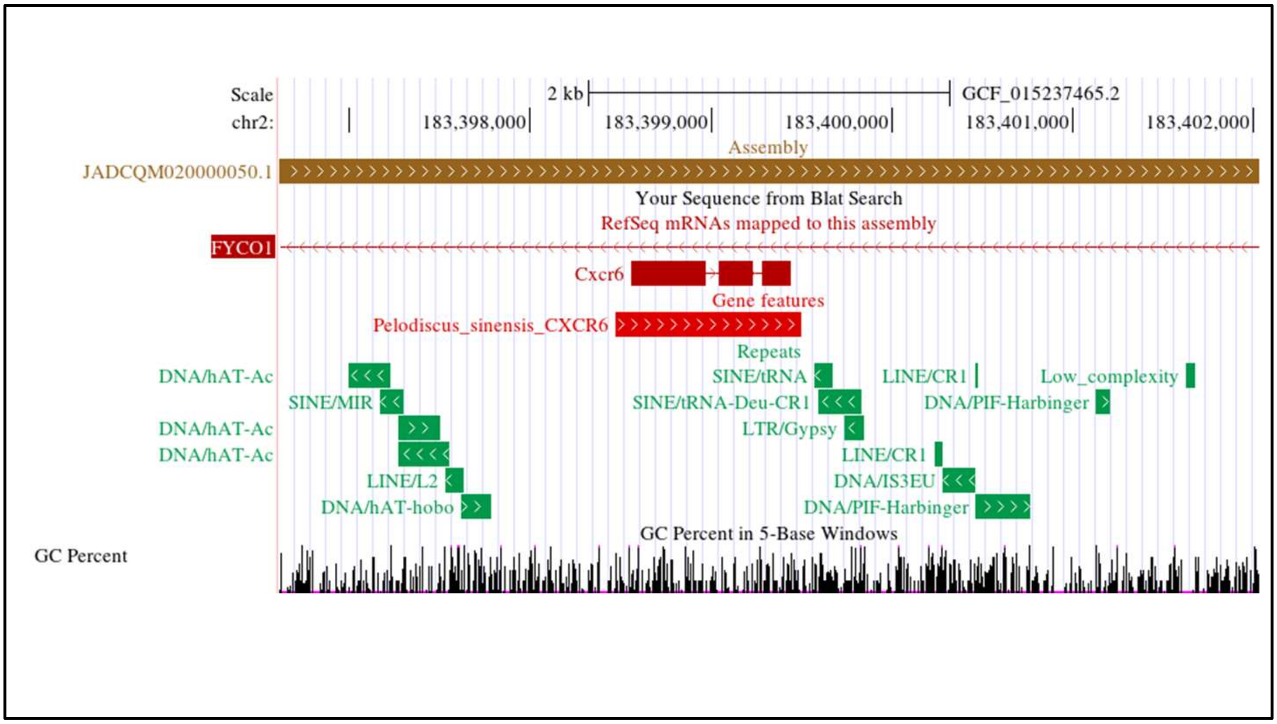


**Supplementary Figure 32:** Repeat identification in the green sea turtle (*Chelonia mydas*) genome assembly (GCF_015237465.2) at the *CXCR6* gene locus using the UCSC genome browser. The Repeat track shows the repeats in that region (represented by dark green boxes). Gene features track shows the *Pelodiscus sinensis CXCR6* gene in *Crocodylus porosus* genome assembly.

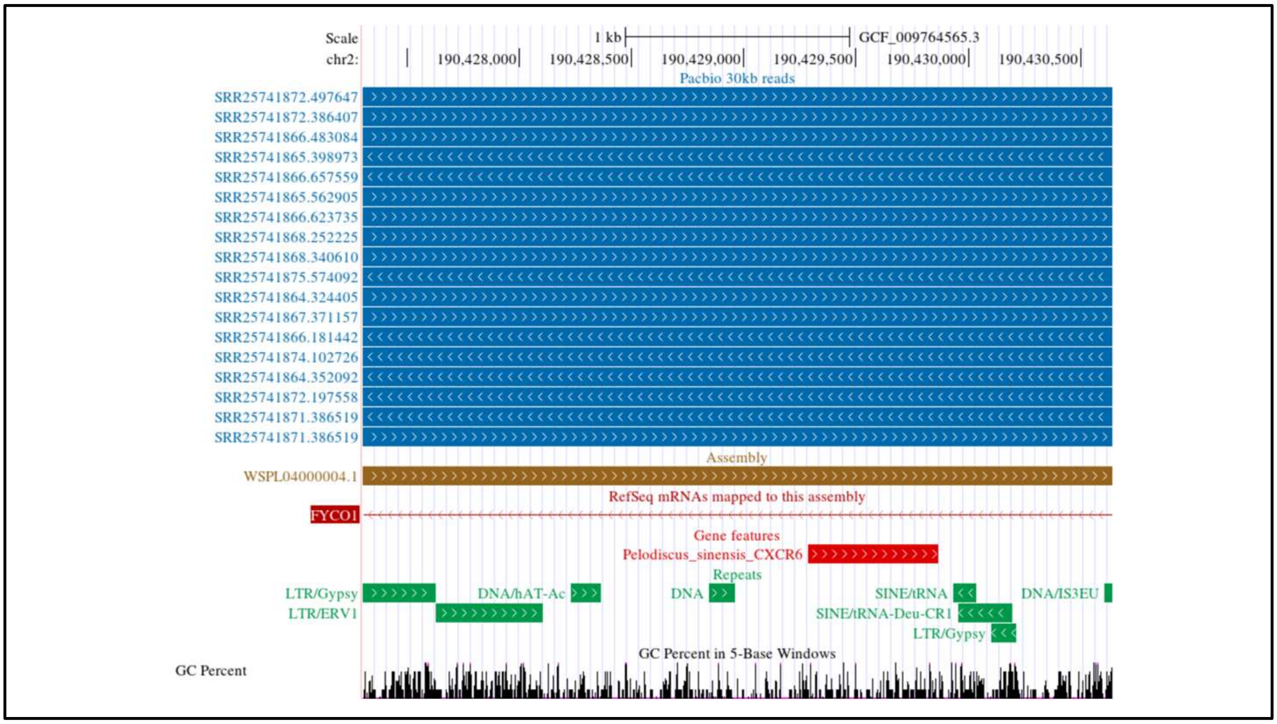


**Supplementary Figure 33:** Verification of the leatherback sea turtle (*Dermochelys coriacea*) genome assembly using long reads at the *CXCR6* gene locus. The UCSC genome browser image showing the alignment of ≥20 kb long reads generated using PacBio sequencing (steel blue coloured), aligned to the *Dermochelys coriacea* (GCF_009764565.3) at the syntenic location of the *CXCR6* gene remnants. The accession read IDs are shown on the left side of the reads. Gene features track shows partial remnants of the *Pelodiscus sinensis CXCR6* gene in *Gavialis gangeticus* genome assembly. Blue-colored boxes represent the *CXCR6* exon remnants. The Repeat track shows the repeats in that region (represented by dark green boxes). Overlapping reads span the entire region at the syntenic location of the *CXCR6* gene, including the flanking region.

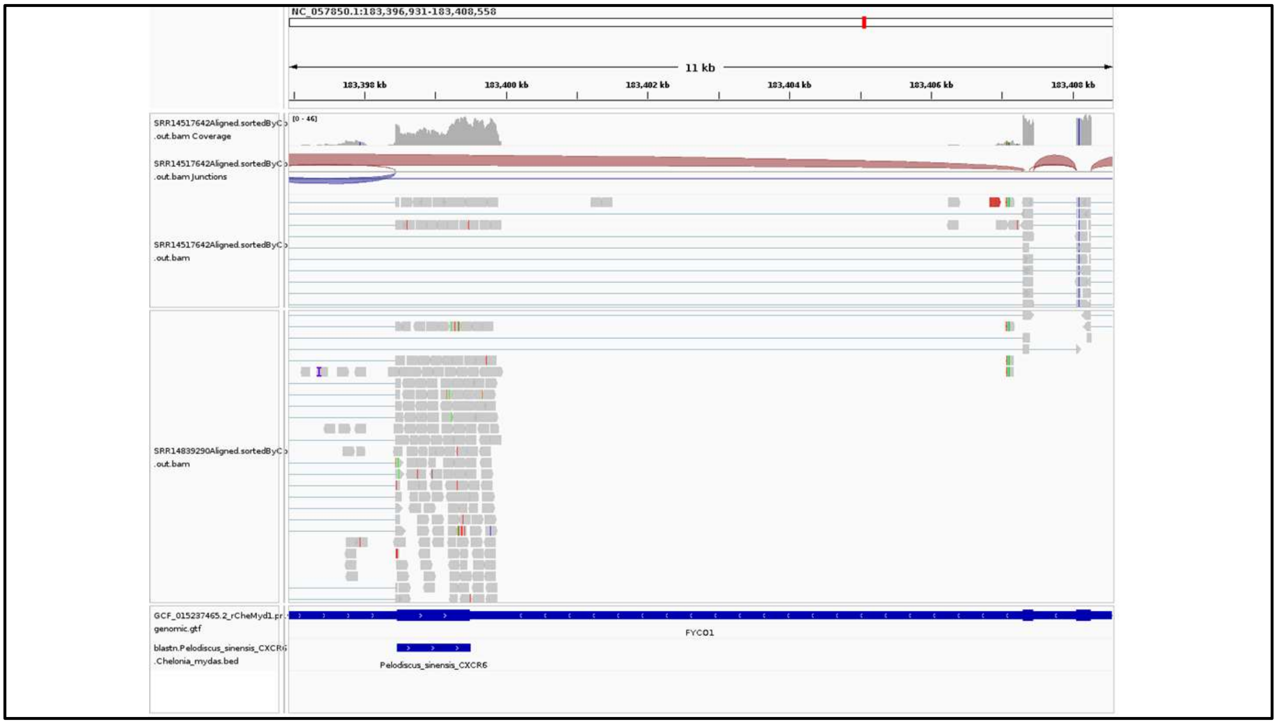


**Supplementary Figure 34:** The *CXCR6* gene expressed in the spleen of a Green sea turtle (*Chelonia mydas*). IGV screenshot for *CXCR6* expression for spleen (SRR14517642 and SRR14839290) mapped to GCF_015237465.2 genome assembly of the Green sea turtle (*Chelonia mydas*), using STAR mapper.

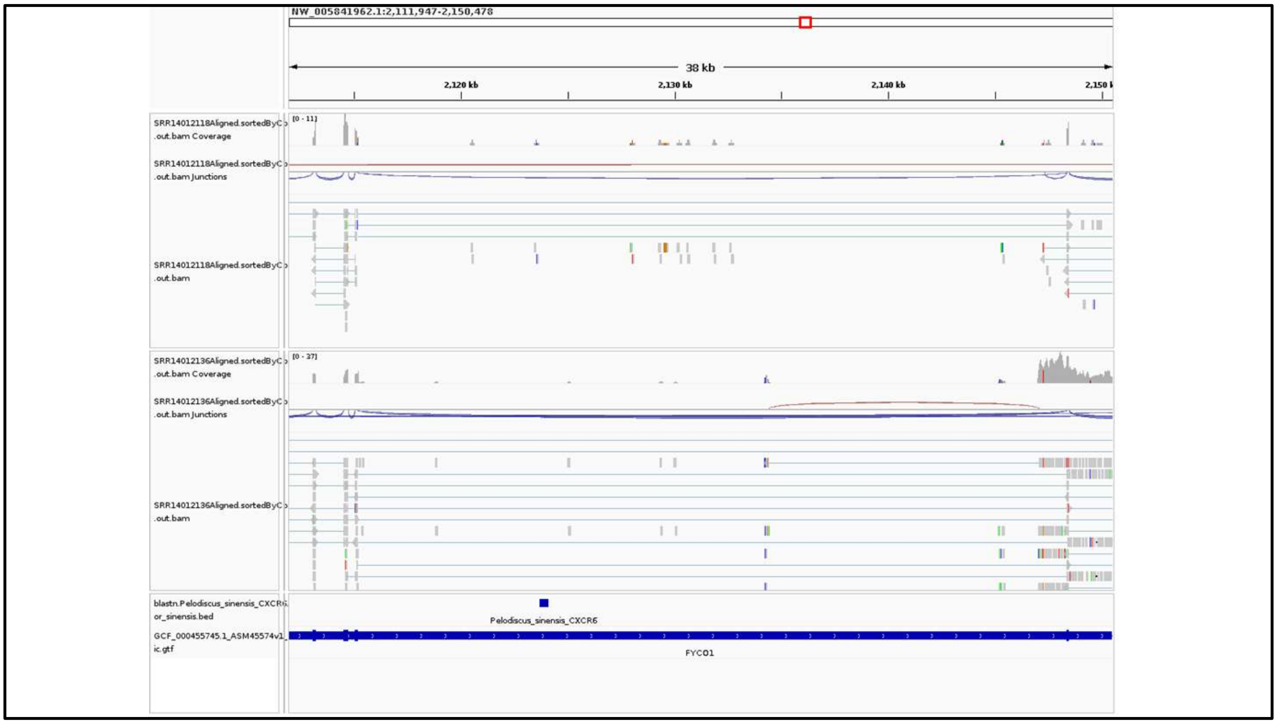


**Supplementary Figure 35:** The *CXCR6* gene lacks expression in the spleen of the Chinese alligator (*Alligator sinensis*). IGV screenshot for *CXCR6* expression for spleen (SRR14012118 and SRR14012136) mapped to GCF_000455745.1 genome assembly of the Chinese alligator (*Alligator sinensis*), using STAR mapper.

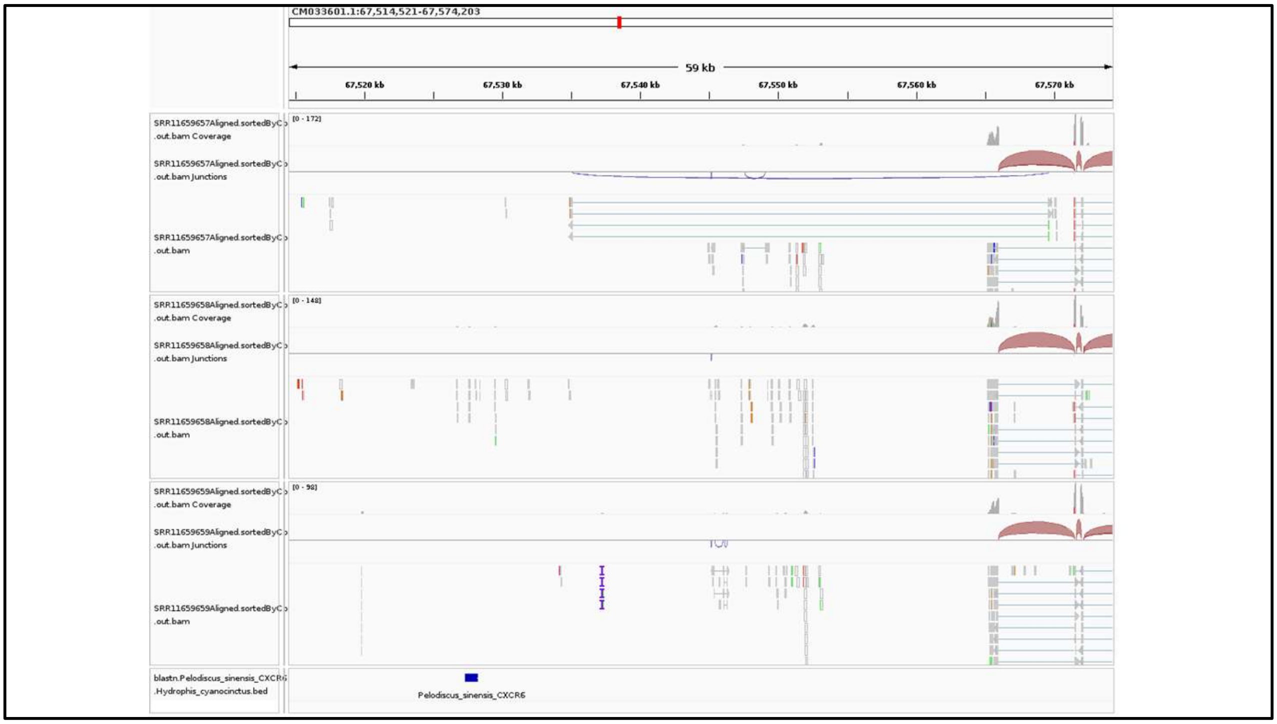


**Supplementary Figure 36:** The *CXCR6* gene lacks expression in the Blue-banded sea snake (*Hydrophis cyanocinctus*). IGV screenshot for *CXCR6* expression for transcriptomic data from SRR11659657, SRR11659658, and SRR11659659 mapped to GCA_019473425.1 genome assembly of the Blue-banded sea snake (*Hydrophis cyanocinctus*), using STAR mapper.

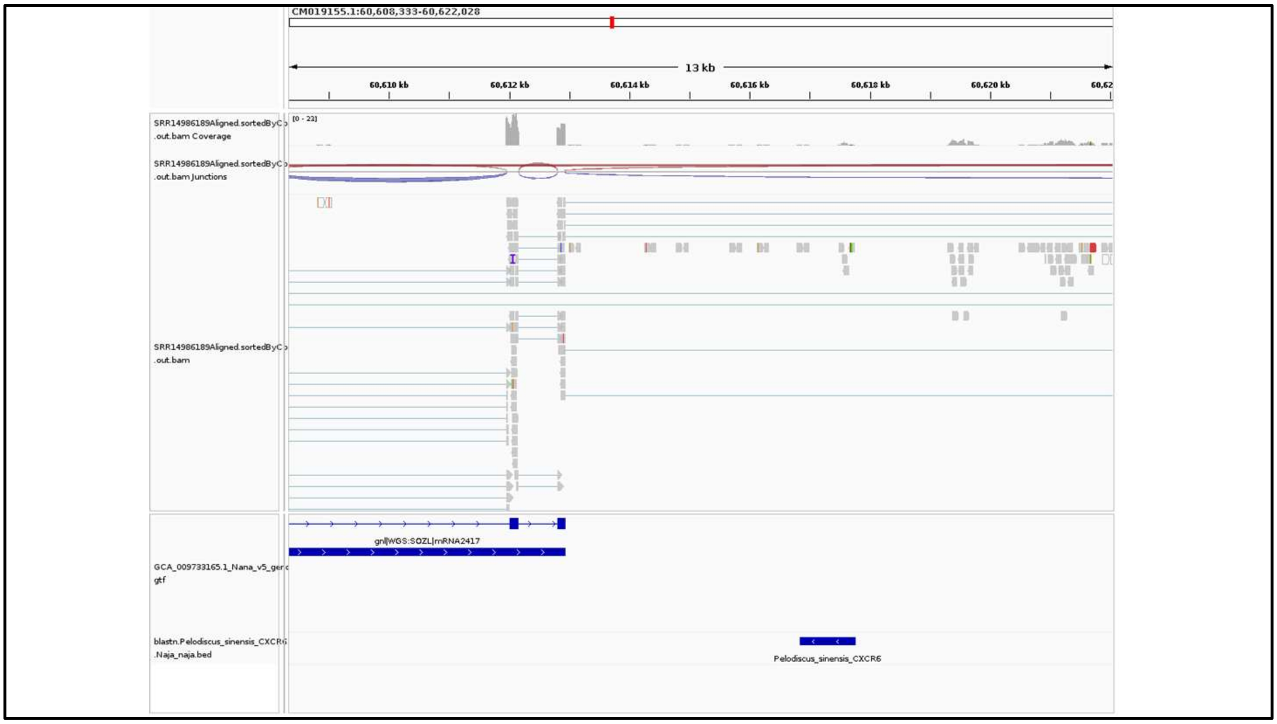


**Supplementary Figure 37:** The *CXCR6* gene lacks expression in the Indian cobra (*Naja naja*). IGV screenshot for *CXCR6* expression for transcriptomic data from SRR14986189 of venom-gland mapped to GCA_009733165.1 genome assembly of the Indian cobra (*Naja naja*), using STAR mapper.

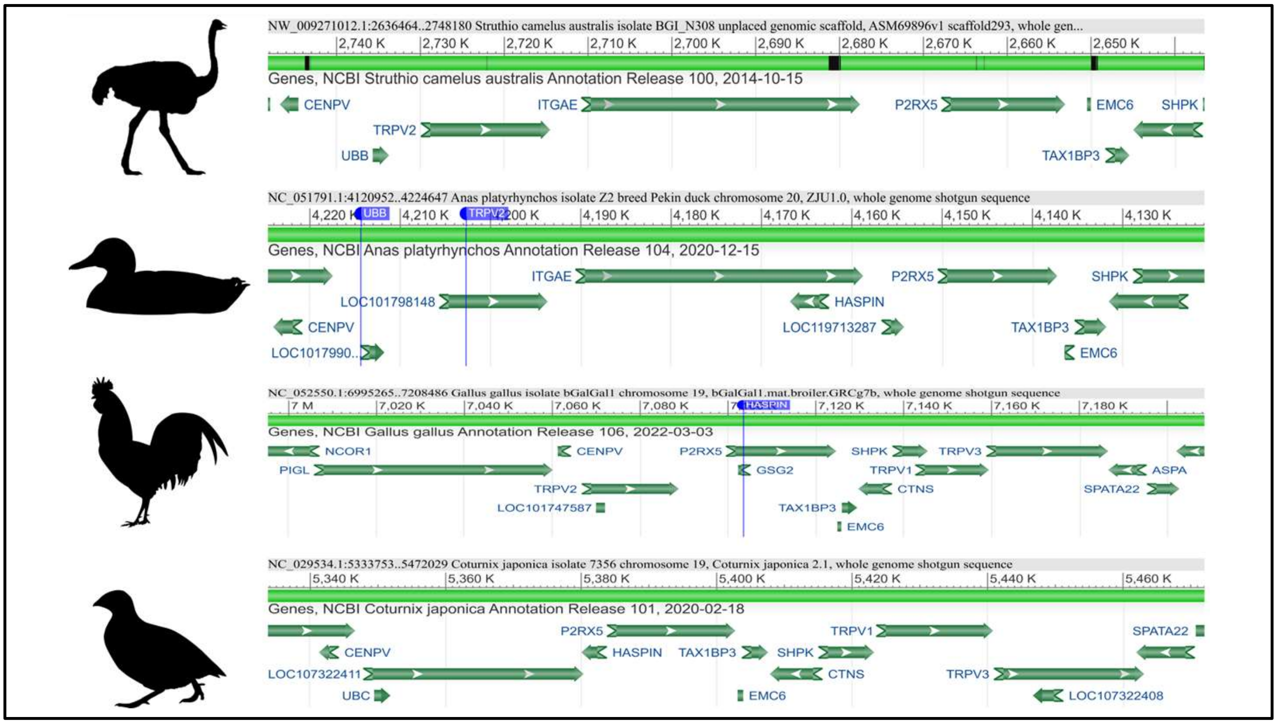


**Supplementary Figure 38:** The NCBI graphical sequence Viewer for the *ITGAE* gene for ostrich, mallard, chicken, and Japanese quail. The *CENPV*, *UBB*, and *TRPV2* flank the *ITGAE* gene on the left side, whereas *HASPIN*, *P2RX5*, and *EMC6* are on the right flank. The annotation for the *ITGAE* gene is missing in chicken and Japanese quail. The Blue marker shows the orthologs for LOC gene IDs. The Silhouette Images are taken from the PhyloPic (https://www.phylopic.org/).

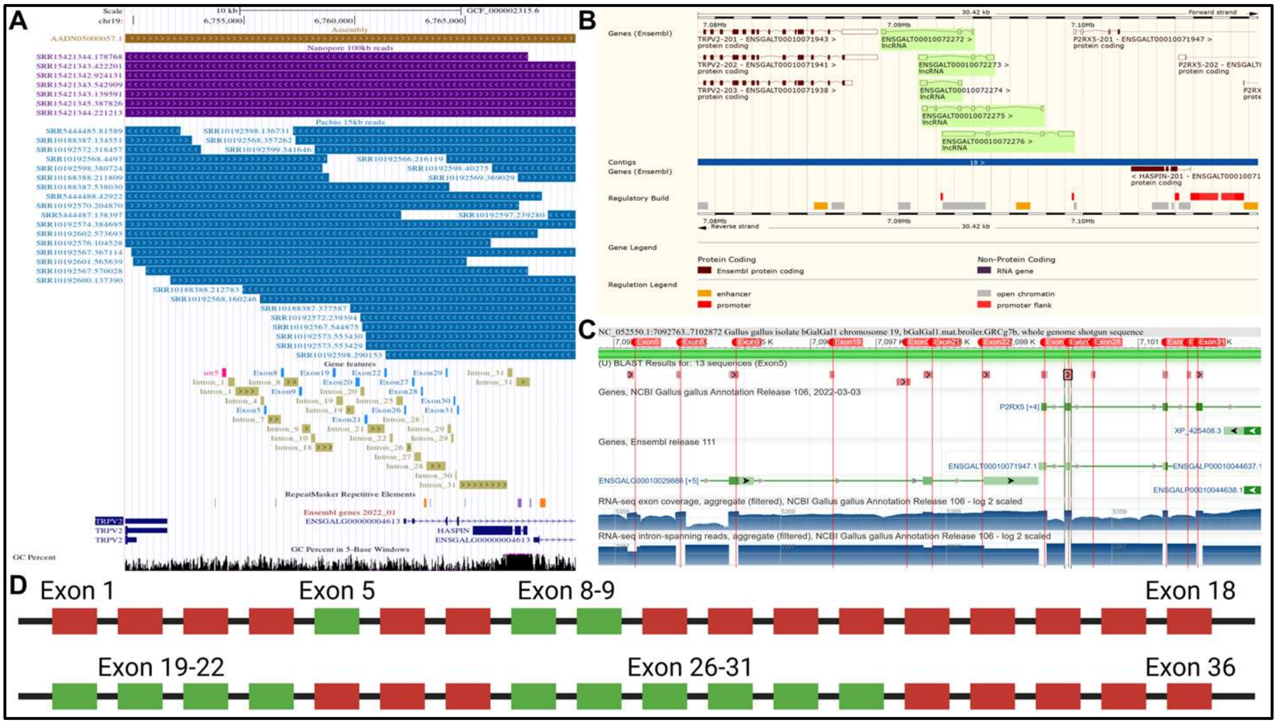


**Supplementary Figure 39: A.** Verification of the chicken (*Gallus gallus*) genome assembly using long reads at the *ITGAE* gene locus. The UCSC genome browser image showing the alignment of ≥15 kb long reads generated using PacBio sequencing (steel blue coloured) and ≥100 kb long Nanopore (purple coloured) aligned to the chicken (GCF_000002315.6) at the syntenic location of the *ITGAE* gene remnants. The accession read IDs are shown on the left side of the reads. Gene features track shows partial remnants of the mallard *ITGAE* gene in chicken genome assembly. Blue-coloured boxes represent the *ITGAE* exon remnants. **B.** ENSEMBL annotation for long non-coding RNAs at the syntenic locus of *ITGAE* in chicken. **C.** NCBI graphical sequence Viewer displaying the expressing exons of *ITGAE* in a chicken's GRCg7b genome assembly. Exons are marked with red tags. **D.** The mallard 36 exonic *ITGAE* gene sequence was used to search for *ITGAE* in chicken. Exons with no BLASTn hits are coloured red, while those with hits obtained and expressed are shown in green.

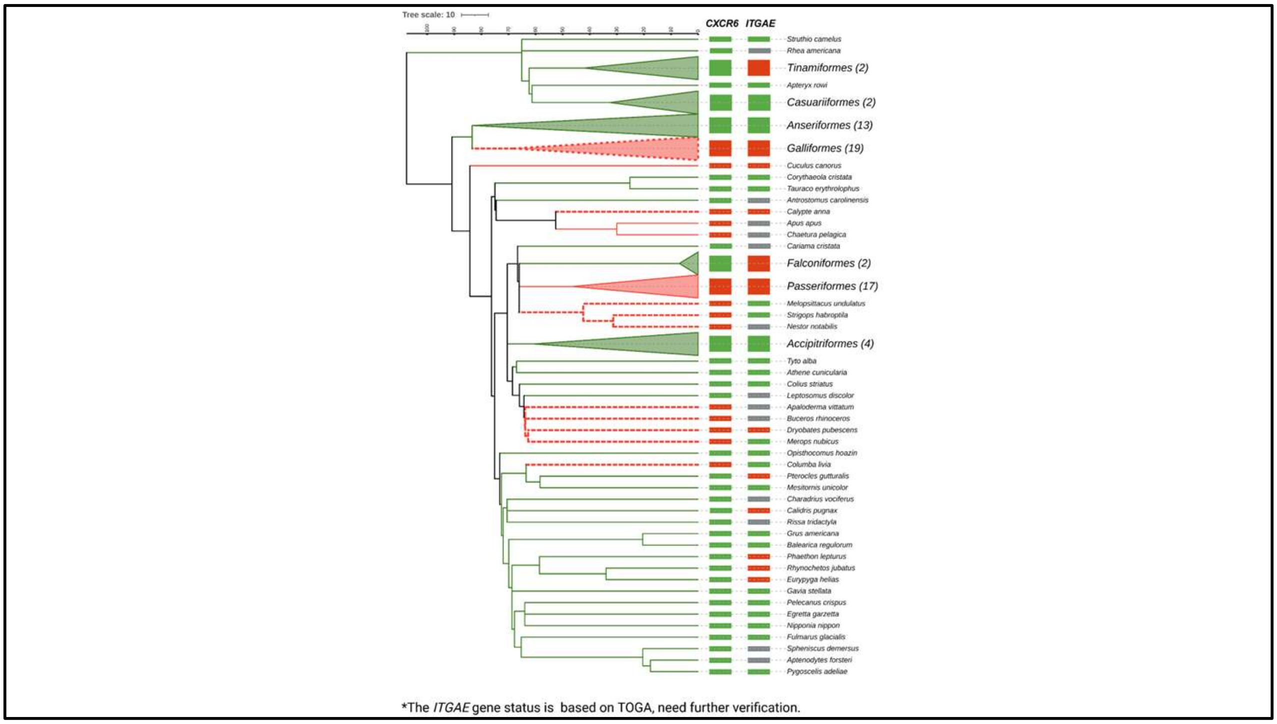


**Supplementary Figure 40:** Status of *CXCR6* and *ITGAE* gene across birds. The *CXCR6* gene is intact or lost, as shown by green- and red-filled rectangles, respectively. The solid red colour branches indicate loss of the *CXCR6* gene due to frame-disrupting changes, whereas dashed phylogenetic branches show *CXCR6* gene loss due to segmental deletion. Green and red-filled rectangles indicate the intactness or loss inference from TOGA for the *ITGAE* gene. The grey-filled rectangle represents the missing/unclear loss/partially intact status of the *ITGAE* gene. The number in brackets in front of the bird order represents the number of species. The time-calibrated phylogenetic tree is downloaded from the Timetree website. The iTOLv6 was used to annotate branches of the phylogenetic tree.

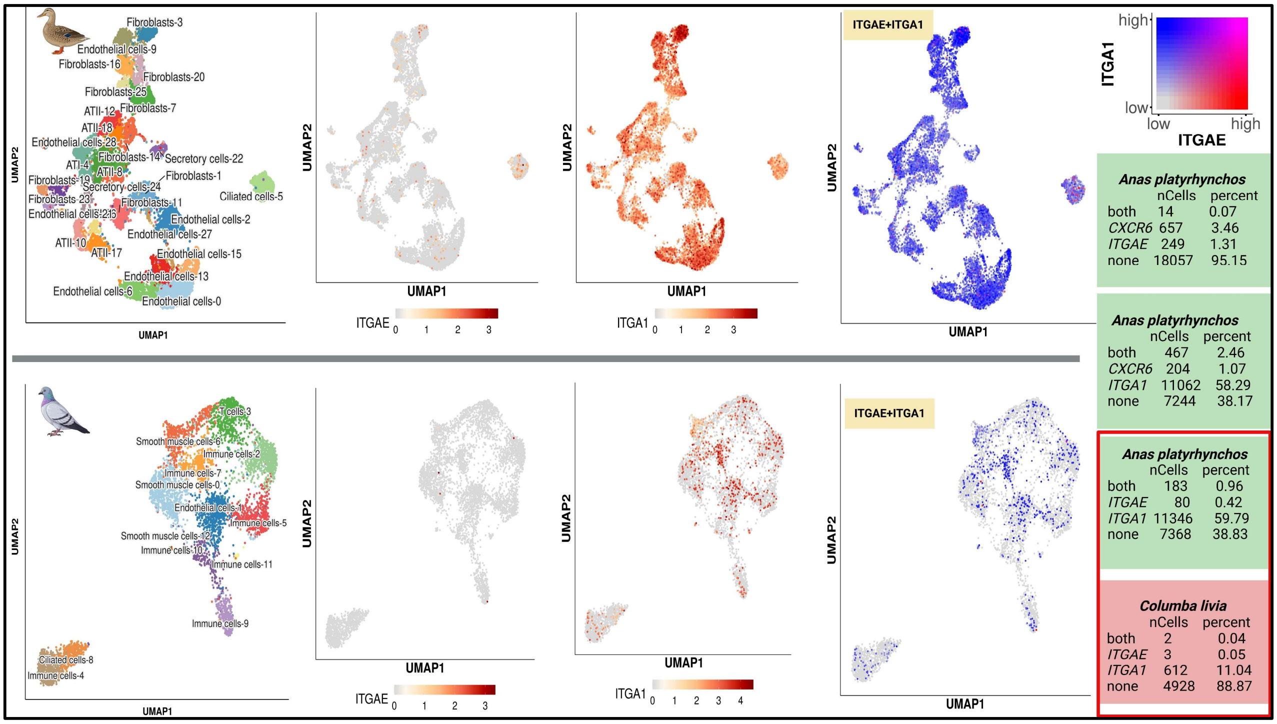


**Supplementary Figure 41:** Comparison of lung CD8 TRM cells between duck and pigeon. The comparison of TRM cell count was made based on TRM marker genes. The top panel displays the UMAP for ducks, while the bottom panel shows pigeons. Cell-type annotations are presented in the first column. The second and third columns depict cells expressing ITGAE and ITGA1, respectively, with expression scales provided below each plot. The fourth column illustrates cells with co-expression of ITGAE and ITGA1, with the expression scale at the top right corner. Light green and light red denotes the number and percentage of cells expressing different gene combinations. Rectangular red brackets highlight ITGAE and ITGA1, cells comparison for duck and pigeon. Data was obtained from the SPEED single-cell atlas (http://speedatlas.net/).

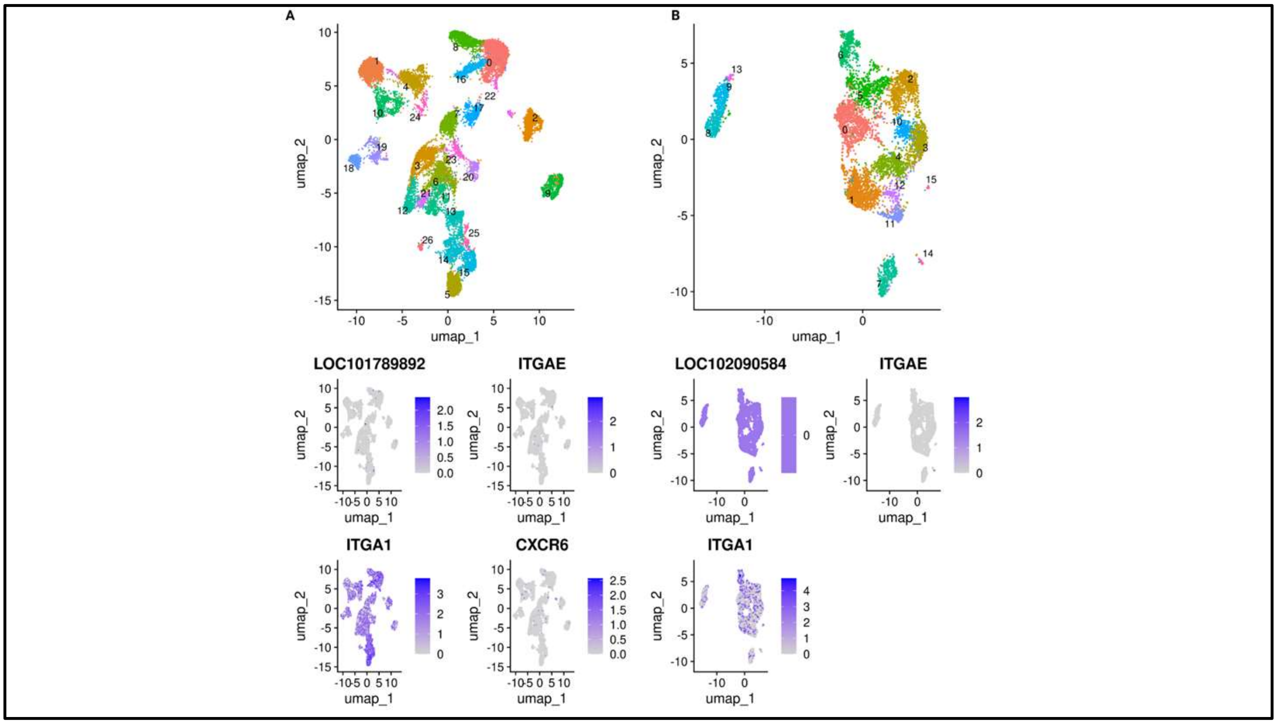

**Supplementary Figure 42:** scRNA-seq analysis of mallard and pigeon lung transcriptomes (NCBI accession: PRJNA747757). **A.** Clusters of cells in the mallard lung scRNA-seq. (top panel). The bottom panel displays feature plots for LOC101789892, *ITAGE*, *ITGA1*, and *CXCR6* for mallard. **B.** Clusters of cells in the pigeon lung scRNA-seq. The bottom panel shows feature plots for LOC102090584, *ITGAE*, and *ITGA1*. The expression scale is provided to the right of each feature plot. Cluster numbers are indicated above each cell cluster. Clusters of cells were created using the UMAP non-linear dimensional reduction method in SeuratV5.

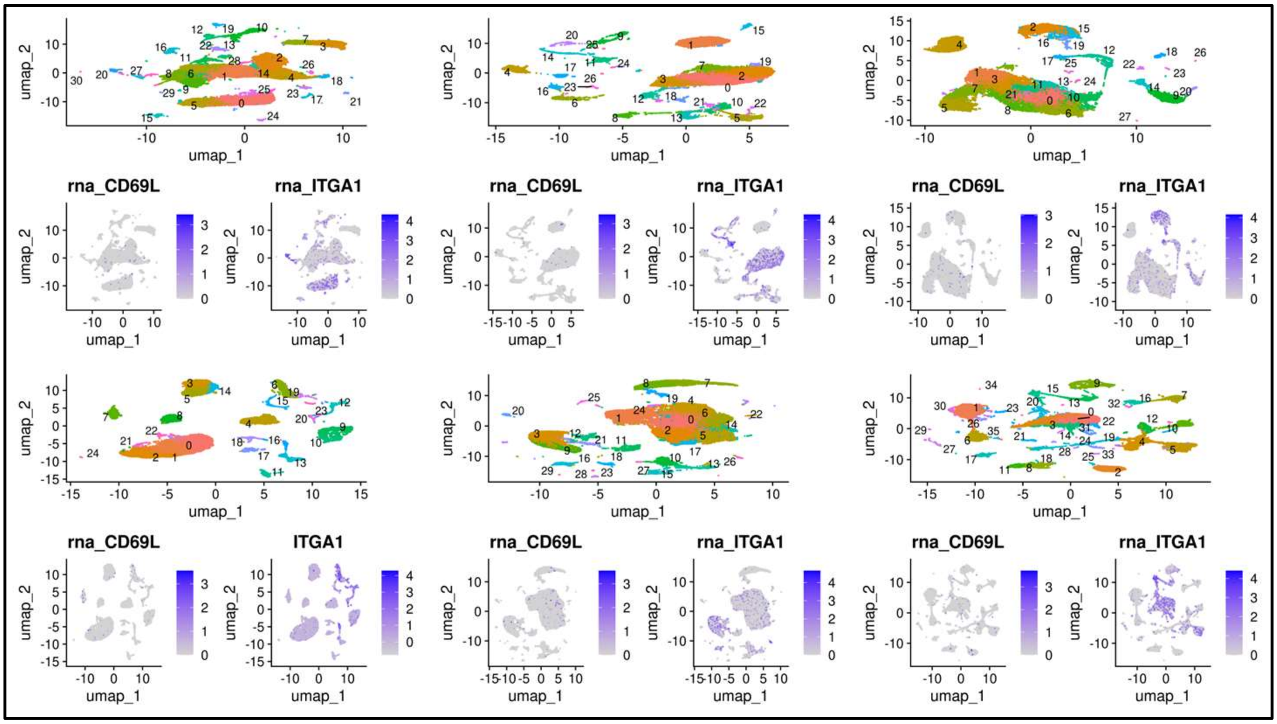


**Supplementary Figure 43:** sc-RNA seq analysis of chicken lung transcriptome for six experimental conditions (NCBI accession code PRJNA834764). Clusters of cells were made using the UMAP non-linear dimensional reduction method in Seurat V5 (Top panel). The feature plots with their expression scale are shown for *CD69L* and *ITGA1* genes. Cluster numbers are indicated above each cell cluster.
