## Supplementary Text for "Evolutionary diversity of CXCL16*-*CXCR6: Convergent Substitutions and Recurrent Gene Loss in Sauropsids"

### Supplementary Text S1: Material and Methods

#### a) Finding of *CXCR6* gene loss

##### (i) Validation of segmental deletions

The *CXCR6* gene locus is flanked by exon14 and exon15 (within intron14) of the *FYCO1* gene, missing from the chicken genome assembly while intact in mallard **(Fig. S18)**. The lack of sequence corresponding to the *CXCR6* locus in the chicken genome may be because of an assembly error, translocation event, or a segmental deletion. We validated the correctness of the genome assembly using PacBio long-read sequencing data in chickens (*Gallus gallus*) and mallards (*Anas platyrhynchos*) to rule out the possibility of an assembly error **(Fig. 6A)**. Our BLASTn search in the short-read and long-read databases **(Table S4)** using the *CXCR6* gene from the mallard genome as a query failed to recover the gene. This further rules out the possibility of translocation. Hence, the lack of such reads matching the *CXCR6* query supports the scenario of a segmental deletion in the chicken genome.

##### (ii) Deletion size estimation

We used mallard as a reference to estimate the approximate deletion size in Galliformes. We obtained the *FYCO1* gene containing chromosome/scaffold from 27

Galloanserae species. We performed a blast-based search of the focal genomes using the mallard genomic sequence from exon14 and exon15 of the *FYCO1* gene. We considered 5 Kb flanking regions beside each hit as the region of interest. The nucleotide sequence from this region was extracted using bedtools v2.27.1 (Quinlan and Hall 2010) and used as a query sequence to search the genomes of focal species. The deletion size is roughly 10 Kb, as estimated by the pairwise alignment between mallard and chicken genomes. Therefore, we used the bedtools merge command with the *-d 5000* option to merge the nearby hits to extend the alignments in regions of poor sequence identity. We found one large region of segmental deletion, orthologous to the *CXCR6* gene region lacking BLASTn hits. The size of the deletion was estimated as the length of this region lacking BLASTn hits, and the resulting bed files were loaded into the UCSC genome browser and visualized **(Fig. 6A-6B)**.

Similarly, to estimate the deletion size of the promoter region of *CXCR6* in Passeriformes compared to the golden eagle (*Aquila chrysaetos*), we followed the same approach but used *bedtools merge -d 2000*. The deletion was doubly confirmed using the dot plot. The xsltproc utility from XSLT sandbox (https://github.com/lindenb/xslt-sandbox) was used to generate dot plots. Additionally, we generated a circos plot using Circos (Krzywinski et al. 2009). GC content from genome data was calculated using the script GCcalc.py (https://github.com/WenchaoLin/GCcalc).

**(iii) Raw read-based verification of gene-inactivating mutations**

Gene loss candidates were initially identified based on the results of the TOGA (Kirilenko et al. 2023) and BLASTn (Camacho et al. 2009) search of the genome assemblies **(Table S4)**. Genome assemblies can have base pair-level errors artefactually introduced during genome assembly. Hence, we verified gene-disrupting events using raw-read datasets. The short-read datasets were searched using BLASTn with the gene sequences as queries to asses raw-read support. In the case of Passeriformes, we used the golden eagle *CXCR6* gene sequence as the query to identify gene-disrupting changes in genome assemblies, which were confirmed based on Illumina SRA BLASTn. In the case of Crocodilia (and *Dermochelys coriacea*), we used a Chinese softshell turtle as the query sequence and obtained truncated BLASTn hits. To identify gene-disrupting changes in the Elapidae family, we used the western terrestrial garter snake (*Thamnophis elegans*) as the query. Further, to find out repeat insertion, RepeatMasker v4.1.5 was used.

**b) Selection pressure analysis**

For molecular evolutionary analysis, the nucleotide sequences of *CXCR6* were downloaded from the NCBI database using the accession IDs. We used the CDSKIT utility (Fukushima and Pollock 2023) to retrieve the ORF's from NCBI GenBank and merged it with CDS for non-annotated species (annotated using TOGA v1.1.6). The nucleotide sequences of *CXCR6* were aligned using PRANK v.170427 in the Guidance program (Sela et al. 2015). In-frame stop codons and frame-disrupting changes are replaced with appropriate N's to ensure the sequences are in the same reading frame. The suitability of MSAs for molecular evolutionary analysis was evaluated using the sequence saturation test implemented in DAMBE v7.3.32 software (Xia and Xie 2001). The time-calibrated phylogenetic trees were obtained from the Timetree website (http://www.timetree.org/). As the avian genomes are strongly impacted by GC-biased gene conversion, the magnitude of gBGC was assessed using Map NH v1.3.0 (Dutheil et al. 2012) and phastBias **(Table S5)**.

We detect the selection pressure on the *CXCR6* gene across orders of vertebrate species using dN/dS metric-based software. The alignment and tree are analyzed by Hyphy v2.3.14 package **(Table S6)** (Kosakovsky Pond et al. 2005) and PAML using the codeML application **(Table S7)** (Yang 2007).

**(i) Site-specific estimation**

To estimate selection strength across the sites, i.e., the number of sites under positive selection (PS) and negative selection (NS), we used FEL (Kosakovsky Pond and Frost 2005). MEME (Murrell et al. 2012) was used to detect sites under PS. The ggplot2 package is used in the R (R Core Team 2021) v4.3.1 program to plot LRT values against codon position for visualization **(Fig. S6-S7)**.

**(ii) Estimating the strength of natural selection**

To screen branches under PS, we used aBSREL (based on the Branch-site model) (Smith et al. 2015), which tests whether a proportion of sites have evolved under PS for foreground in the phylogeny. The gene-wide screening of foreground species for PS was done using BUSTED (Murrell et al. 2015). Further, the trends and/or shifts in the stringency of natural selection on a *CXCR6* gene, we tested with RELAX program (Wertheim et al. 2015) of Hyphy package (p-Value $< 0.05$ considered as significant, $k>1$ indicates intensification whereas $k < 1$ indicates relax selection strength along the foreground branch) **(Table S6)**. Additionally, the codeML test implemented in PAML was used to compare the strength of selection on *CXCR6* across the given clade using M0 (null), b_free (PS), and b_neut (relax selection) branch models **(Table S7)**.

**c) Single-cell RNA-seq analysis**

We retrieved lung single-cell transcriptome sequencing data from the NCBI database under accession code PRJNA747757 (for mallard and pigeon) (Chen et al. 2021) and PRJNA834764 (for chicken) (Liu et al. 2023). The genome assemblies of mallard (GCF_015476345.1), pigeon (GCF_000337935.1), and chicken (GCF_016699485.2) were used along with their genome annotation files. We labelled mitochondrial genes with "MT-" as a prefix. Cell Ranger v7.2.0 (10× Genomics) was used to process these single-cell transcriptomic data, gene expression matrices were obtained, and all gene expression matrices were processed with R package Seurat v5. Briefly, quality control was performed based on the following criteria: cells with a mapped number of genes <200 or with a mitochondrial percentage higher than 10% were removed. Normalization is done with Seurat's NormalizeData function. Variable genes were determined using Seurat's FindVariableGenes function with default parameters (selection.method = "vst", nfeatures = 2000). Clusters were identified via the FindClusters in Seurat subsequently visualized using the RunUMAP function and DimPlot (reduction = 'umap' ) and FeaturePlot with *LOC101789892*, *ITGAE,* and *ITGA1* genes of mallard, *LOC102090584*, *ITGAE*, *ITGA1* genes of pigeon, and *CD69L* and *ITGA1* gene in chicken using R package Seurat v5. (Hao et al. 2024).

**d) *CXCL16* assembly verification and retrieval in birds, Crocodilia, and snakes**

We used Klumpy (Madrigal et al. 2024) to assess the assembly at the *CXCL16* locus. The long-read SRA data from PacBio and/or Nanopore were mapped to the chromosome or scaffold containing *CXCL16* or *MED11* and *CLDN7*. Minimap2 (Li 2018) was used for mapping with the *-ax map-hifi, -ax map-pb* and *-ax map-ont* options to map PacBio HiFi, PacBio sequel, and Nanopore reads, respectively. Furthermore, TOGA was used to identify an ORF using harpy eagle and great cormorant *CXCL16* sequences as references. As the *CXCL16* locus has high GC content, it may have been assembled based on long reads, which is evident from multiple frame-disrupting changes detected by TOGA. We mapped RNA-seq data using the STAR (Dobin 2019) read aligner, which was then used as input for Pilon (Walker et al. 2014) to correct the scaffold or chromosome. We used *--fix all --mindepth 0.5 --changes --verbose --minqual 20 --minmq 20*, this flags to run Pilon. The corrected FASTA output from Pilon was used again as the query sequence in TOGA, and with this approach, we recovered the ORF of *CXCL16* in the respective species.

### Supplementary Text S2: Status of *CXCL16* among vertebrates

a) **High GC-content**

Genomic clusters of "missing" genes coincide with genomic regions with high GC content (Chen et al. 2013; Huttener et al. 2021). Moreover, instances of false predictions of gene loss due to elevated GC content have been documented in avian species orthologs (Hron et al. 2015; Beauclair et al. 2019; Kim et al. 2022). Along with the high GC content, several avian genes are characterized by long G/C stretches (Hron et al. 2015). For example, the GC-rich chicken *TNF-α*, a vital cytokine in host immune defence, was initially considered absent in avians and later recovered in chickens and other birds (Rohde et al. 2018). Similarly, the *LAT* (involved in T-cell antigenic signalling), whose orthologs were missing in birds, was later identified in birds (Janusova et al. 2023). These challenges are mainly caused by the difficulty in sequencing such regions by short-read sequencing, such as Illumina (Tilak et al. 2018). High-quality chromosome-level genome assemblies generated using long-read sequencing technologies such as PacBio and Nanopore have shown promise in recovering many of these missing genes (Zhu et al. 2023). However, GC-rich and GC-poor regions have low read coverage in Illumina datasets (Chen et al. 2013). In our investigation, we observed a similar increase in GC content and G/C stretches in the case of birds and reptilian *CXCL16* orthologs compared to mammals but not in the *CXCR6* gene orthologs **(Fig 1B)**.

b) **High sequence divergence**

Interspecific divergence in immune gene expression and/or coding sequence may account for variations in immunological function between species (Zhong et al. 2021). The genes encoding pattern recognition receptors, cytokines, and chemokines had higher sequence divergence than signal transduction proteins and transcription factors (Zhong et al. 2021). In the case of *CXCL16* and *CXCR6,* such genetic variation is reported between the species of rodents and primates (Xu et al. 2018; He et al. 2020). Mice and human *CXCL16* ortholog have only 49 % amino acid identity and 70 % similarity (Matloubian et al. 2000). To find the highly divergent gene, we used a previously used approach, i.e., tBLASTn. However, we did not find *CXCL16* (spurious blast hits were obtained for other chemokines). The sequence identity of Harpy eagle *CXCL16* is compared with other vertebrate species and found to have high sequence divergence similar to *TNF-α* and *LAT* genes **(Fig. 1C)**.

**c) Physical position**

The physical location of a gene can be a deciding factor for its functionality and elusiveness (Hara and Kuraku 2023). The chemokines are often clustered, and gene density is also high in that region (Nomiyama et al. 2010, 2011). While *CXCL16* is unlike other chemokines, it is singly present in human chr 17 (Nomiyama et al. 2010). Another reason genes are reported as missing is their presence on microchromosomes (Huttener et al. 2021; Hara and Kuraku 2023). The *CXCL16* ortholog in human, mouse, loggerhead turtle, great cormorant, harpy eagle, ostrich and American alligator is located on chromosomes 17, 11, 27, 34, 26, 37 and 15, respectively **(Table S2)**.

**d) Sample contamination**

While screening for *CXCL16* in mallards short read database using human ortholog as BLASTn query, we got full-length *CXCL16* intact coding DNA sequence using a huge SRA database from the study, NCBI Accession: PRJNA896757 (Li et al. 2023) of size ~874 Gb. Upon careful verification, we found the sample contaminated with human genomic DNA **(Fig. S1)**.

**e) Recovery of *CXCL16* in birds**

Chromosome-level genome assemblies generated using PacBio, Nanopore, and Hi-C technologies provide valuable resources for identifying missing high-GC content genes. Recent advancements in producing complete, error-free genome assemblies of vertebrate species, led by the Vertebrate Genome Project, offer an essential foundation for this purpose (Rhie et al.

2021). The ortholog of *CXCL16* has been recently annotated in great cormorant (*Phalacrocorax carbo*) on NCBI (last accessed on 31st August 2024) with RefSeq transcript ID XM_064437657, located on chromosome 34 (NC_087546.1; GC-content = 65 %). The genome of the great cormorant (*Phalacrocorax carbo*) (GCF_963921805.1) was assembled using PacBio and Arima2 Hi-C sequencing methods, demonstrating their effectiveness in assembling complex genomic loci. Furthermore, our BLASTp search on the Non-redundant protein sequences (nr) database using XP_064293727.1 as the query obtained hits in two other bird species, *Harpia harpyja* (XP_052631027.1) and *Accipiter gentilis* (XP_049648608.1), both with a query cover of ≥ 80%. Notably, both GCF_026419915.1 (*Harpia harpyja*; https://vertebrategenomesproject.org, (Canesin et al. 2024)) and GCF_929443795.1 (*Accipiter gentilis*) (August et al. 2022) were assembled using PacBio and Arima2 Hi-C, highlighting the importance of such high-quality genome assemblies for recovering genes like *CXCL16*. Considering the high GC content and significant sequence divergence, we employed three approaches to recover the *CXCL16* gene in birds:

(i) **Blast search**: A BLASTn search of the great cormorant (*Phalacrocorax carbo*) *CXCL16* sequence XM_064437657.1 against the NCBI nt database yielded hits in five species with query coverage ≥ 80%: (1) *Phalacrocorax carbo* (chromosome 34; OY997636.1; Query cover 100%), (2) *Buteo buteo* (chromosome 12; OZ076931.1; Query cover 94%), (3) *Harpia harpyja* (chromosome 26; NC_068965.1; Query cover 90%), (4) *Calonectris borealis* (chromosome 36; OZ077845.1; Query cover 87%), (5) *Accipiter gentilis* (chromosome 35; OV839396.1; Query cover 81%), and (6) *Haliaeetus albicilla* (chromosome 13; OX381650.1; Query cover 80%) (BLASTn results, last accessed on 31st August 2024).

(ii) **Based on synteny with *MED11***: Given the high sequence divergence of *CXCL16* with its orthologs, we utilized *MED11* to identify the chromosomes in the respective species using NCBI datasets with Gene ID 400569. We obtained chromosome data for 32 bird species (see **Supplementary Table S2**) by using the eSearch utility of NCBI to retrieve the relevant chromosomes from the nucleotide database (Sayers 2018). In most birds, *MED11* is singly annotated. However, for cases where syntenic genes were identified, we employed TOGA (Kirilenko et al. 2023) for orthologs gene annotation and screened the *CXCL16* in those scaffolds. For instance, based on *MED11* annotation, we identified *CXCL16* in Darwin's rhea (*Rhea pennata*), a species within the Palaeognathae infraclass.

(iii) **Species-specific approaches**:

a. **Chicken** (*Gallus gallus*): Based on a BLASTn search for *MED11* in chicken genome assemblies using the great cormorant sequence, we found hits on chromosome 35 of the chicken Huxu breed genome (GCA_024206055.2). Notably, no BLAST hits were detected for *CXCL16* and *ZMYND15*. The genome assembly was verified by mapping long Nanopore reads and visualized using the UCSC Genome Browser and Klumpy. We observed that multiple Nanopore reads (SRA datasets SRR15421342 and SRR15421343) spanned the entire syntenic locus of *CXCL16*, ruling out the possibility that *CXCL16* was missing in the chicken genome due to an assembly artefact (**Supplementary Figure S2)**. Furthermore, no BLASTn hits for the *CXCL16* sequence (of other bird species) were recovered, and TOGA analysis inferred its loss. Thus, *CXCL16* has been concurrently lost with *CXCR6* in chickens.

b. **Ostrich** (*Struthio camelus*): We screened the genome assembly of ostrich (GCA_040807025.1) for *CXCL16* using BLASTn and found hits on chromosome 37 (CM081622.1). Synteny is conserved, with *ARRB2* and *MED11* on the left flank of *CXCL16* and *ZMYND15* on the right flank. We mapped long-read PacBio HiFi datasets (SRR29936073, SRR299360734, and SRR299360735) using Minimap2 to verify the assembly. Additionally, to assess the expression status of *CXCL16*, we used transcriptomic data from ostrich SRA IDs SRR9031299 (spleen lymphocyte) and SRR12237020, mapping it to the reference genome using the STAR read mapper. *CXCL16* is expressed in ostrich (**Supplementary Figure S1)**. Furthermore, the assembly was verified using Klumpy, and the *CXCL16*'s intactness and coding sequence were inferred based on TOGA. The GC content of the recovered coding sequence is 74.83%.

c. **Mallard** (*Anas platyrhynchos*): We screened the available genome assemblies of mallard for *CXCL16*, but no BLASTn hits were detected. However, we created a database using PacBio HiFi data from ERR12875150, in which we identified the read ERR12875150.3600316, which gave BLASTn hits for *CXCL16*, as well as *MED11* and *CLDN7*. We polished this read with RNA sequencing data from ERR13493936 and SRR23561719-38 using Pilon and obtained the orthologous *CXCL16* sequence through TOGA, using *Harpia harpyja* as the reference. TOGA detected a splice site change and a two-base deletion in the fourth exon, although confirming this is challenging due to the high GC content of the obtained CDS (73.68%).

d. **Swan goose** (*Anser cygnoides*): We recovered a partial transcript from chromosome NC_089903.1 of GCF_040182565.1 using TOGA. The GC content of the partially recovered sequence is 79.34%.

e. **Zebra finch** (*Taeniopygia guttata*): Based on the annotation of the flanking gene *MED11* (XM_041712301.1) in the zebra finch genome on chromosome

32 (NC_054764.1, GCF_003957565.2), we found that synteny is conserved on the left flank with *PELP1* (XM_041712172.1) and *MED11* (XM_041712301.1), while the right flank contains *CTDNEP1* (XM_041712291.1), *ELP5* (XM_041712300.1), and *TAF6* (XM_041712173.1). The genome assembly has a gap between *MED11*-*CTDNEP1* and *ELP5*-*TAF6*. We used the UCSC Genome Browser and Klumpy to assess this region. The genes *CXCL16*, *ZMYND15*, and *CLDN7* are missing. However, we found a common read, SRR25741339.4564582, in the PacBio-HiFi datasets (SRR25741332, SRR25741333, SRR25741334, SRR25741339, and SRR25741347), which gave BLASTn hits to *MED11* and *ELP5*. No BLASTn hits were found for *CXCL16* (using sequences from other birds), *CLDN7*, or *ZMYND15* from Phalacrocorax carbo on SRR25741339.4564582. Notably, we obtained BLASTn hits for *CXCL16*, *CLDN7*, and *ZMYND15* from the PacBio-HiFi database (SRR25741332, SRR25741333, SRR25741334, SRR25741339, and SRR25741347). The hits may be from other chemokines. Therefore, it is difficult to infer the status of *CXCL16* in the zebra finch due to assembly artefacts due to noisy PacBio-HiFi data.

f. **Green sea turtle** (*Chelonia mydas*): We screened the *CXCL16* (XM_048834045.1), *MED11* (XM_048834055.1), *ZMYND15* (XM_048834056.1), and *PRSS55* (XM_048834057.1) genes of Loggerhead sea turtle (*Caretta caretta*) in the genome assembly of the green sea turtle (GCA_015237465.2). None of these genes showed BLASTn hits with more than 80% query coverage. Given the high GC content and sequence divergence, we screened for *CXCL16* and syntenic genes in the PacBio SRA data (SRR25743018, SRR25743022, SRR25743023, SRR25743024,

SRR25743025, SRR25743026, and SRR25743027) using BLASTn. We identified a common read, SRR25743026.493011, with a read length of 48,485 bp, which gave BLASTn hits to the *CXCL16*, *MED11*, and *ZMYND15* genes. The read was corrected using datasets SRR12540953, SRR14517642, and SRR14839290, yet TOGA detected multiple frame-disrupting changes. Additionally, we performed BLASTn with the TOGA-recovered sequence against Hi-C sequencing data (SRR25743019) and 10X linked-reads data (SRR25743020, SRR25743021, SRR25743029, and SRR25743030). No significant similarity was found in the Hi-C data. However, most of the gene was covered by the 10X linked reads. Nonetheless, the entire reading frame could not be reconstructed using the 10x linked reads. Thus, it is difficult to infer the gene status due to limitations in the available data.

**f) Status of *CXCL16* in Crocodilia**

1. American alligator (*Alligator mississippiensis*): *MED11* (XM_006262906.4) and *ZMYND15* (XM_059717839.1) are annotated in the genome assembly of *Alligator mississippiensis* (GCF_030867095.1) on chromosome 15 (NC_081838.1), whereas *CXCL16* is not annotated. Additionally, no BLASTn hits for *CXCL16* were found between these genes when searching with bird *CXCL16* as the query sequence. We verified the genome assembly by mapping long-read data (SRR30180798 and SRR30180799) using Minimap2 and visualized the results with IGV and the UCSC Genome Browser (**Supplementary Figure S3)**. TOGA screening revealed the deletion of all *CXCL16* exons. Since the flanking genes of *CXCL16* are present and the assembly is accurate, it appears that the region containing *CXCL16* has been deleted, suggesting the loss of *CXCL16* due to a segmental deletion.

2. Chinese alligator (*Alligator sinensis*): *ARRB2* and *MED11* are annotated in the *Alligator sinensis* genome assembly (GCF_000455745.1) on scaffold NW_005842389.1. The assembly gaps are found on the UCSC genome browser. Also, the TOGA analysis indicates that the exons of *CXCL16* and *ZMYND15* are missing, suggesting an assembly gap.
3. Gharial (*Gavialis gangeticus*): We found BLASTn hits for *MED11* and *ZMYND15* on chromosome 15 (CM057603.1) of the *Gavialis gangeticus* genome assembly (GCA_030020295.1). Analysis of spanning long-read data and Klumpy alignment suggests the assembly is accurate. TOGA screening revealed the deletion of all *CXCL16* exons. Since the flanking genes of *CXCL16* are present and the assembly is correct, it appears that the region containing *CXCL16* has been deleted, indicating the loss of *CXCL16* due to a segmental deletion.

**g) Status of *CXCL16* in Snakes**

We screened *CXCL16* in the genome assemblies of *Pseudonaja textilis*, *Hydrophis cyanocinctus*, and *Naja naja*, species from the Elapidae family of snakes where *CXCR6* is lost. In the cases of *Pseudonaja textilis* (GCF_900518735.1) and *Hydrophis cyanocinctus*, no BLASTn hits were found for *CXCL16* (XM_039324894.1) with *Crotalus tigris* query with a query cover ≥ 10%.

We screened the genome assembly of *Naja naja* (GCA_009733165.1) using BLASTn with the *CXCL16* sequence from *Crotalus tigris* (XM_039324894.1). BLASTn hits on the scaffold SOZL01001329.1 could not be investigated due to missing genes on the right flank. However, BLASTn hits were also obtained on scaffold SOZL01001525.1 with a query coverage of 36%. Gene synteny is conserved at this locus, with *MED11* (KAG8147999.1) on the right flank and *PRSS55* (KAG8147998.1) and *ZMYND15* (KAG8147997.1) on the left flank (**Supplementary Figure S4)**. TOGA identified multiple frame-disrupting changes in the *Naja naja* genome, but data limitations make it challenging to corroborate these findings with SRA data, particularly regarding the frame-disrupting changes. However, deletion verification was possible with noisy long-read data. The overlapping and spanning reads from the Nanopore (SRR10418060-SRR10418075) and PacBio (SRR10418082-SRR10418089) data rule out the possibility of an assembly artefact. Using long-read databases from Nanopore (SRR10418060-SRR10418075) and PacBio (SRR10418082-SRR10418089), no BLASTn hits were found for Intron 2 and Exon 3 in either database (**Supplementary Figure S5)**. These suggest that the *CXCL16* is lost in the Indian cobra (*Naja naja*) concurrently with *CXCR6*.

We identified the receptor loss (*CXCR6*) in three major lineages (i.e., birds, Crocodilia, and elapids). Our extensive re-analysis using all currently available data could establish that the loss of the ligand (*CXCL16*) is concurrent with the loss of the receptor in at least one species from each of the three lineages. However, despite these advances, limitations in the genome assembly quality and the lack of high-coverage data from multiple sequencing platforms restrict the inference of confident gene loss/presence. The future availability of karyotype-complete gap-free chromosomal-level telomere-to-telomere haplotype-phased assemblies will provide a conclusive answer as the tools to perform genome annotation and infer gene loss are currently available.
